# Challenges and considerations for reproducibility of STARR-seq assays

**DOI:** 10.1101/2022.07.27.501795

**Authors:** Maitreya Das, Ayaan Hossain, Deepro Banerjee, Craig Alan Praul, Santhosh Girirajan

## Abstract

High-throughput methods such as RNA-seq, ChIP-seq and ATAC-seq have well-established guidelines, commercial kits, and analysis pipelines that enable consistency and wider adoption for understanding genome function and regulation. STARR-seq, a popular assay for directly quantifying activity of thousands of enhancer sequences simultaneously, has seen limited standardization across studies. The assay is long with >250 steps, and frequent customization of the protocol and variations in bioinformatics methods raise concerns for reproducibility of STARR-seq studies. Here, we assess each step of the protocol and analysis pipelines from published sources and in-house assays, and identify critical steps and QC checkpoints necessary for reproducibility of the assay. We also provide guidelines for experimental design, protocol scaling, customization, and analysis pipelines for better adoption of the assay. These resources will allow better optimization of STARR-seq for specific research needs, enable comparisons and integration across studies, and improve reproducibility of results.

## Introduction

Enhancers are *cis*-acting DNA elements that regulate gene expression (Banerji et al. 1981). The ability of enhancers to recruit transcription factors through specific binding motifs to regulate the expression of target genes in a cell, tissue, and developmental stage-specific manner make them critical components of gene regulatory networks (Shlyueva et al. 2014). While enhancer-reporter assays (Banerji et al. 1981) and comparative genomics (Pennacchio et al. 2006) enabled initial discoveries, enhancers are usually mapped within nucleosome-free open chromatin regions that are hypersensitive to DNase I (Gross and Garrard 1988) or accessible to transposase (Buenrostro et al. 2013), and to sequences bound by specific transcription factors (Visel et al. 2009) or histone modifications (Heintzman et al. 2007). Such assays that are used to detect putative enhancer regions throughout the genome include DNase-seq (Boyle et al. 2008), ATAC-seq (Buenrostro et al. 2013), and ChIP-seq (Robertson et al. 2007). However, these methods are limited to only providing the location of candidate enhancers and do not assess their functional activity.

Self-Transcribing Active Regulatory Region sequencing or STARR-seq, like other episomal massively parallel reporter assays (MPRA) (Patwardhan et al. 2012; Melnikov et al. 2012) directly quantifies enhancer activity by relying on transcription factors within a host cell system, thereby removing any chromatin-associated biases (Arnold et al. 2013). STARR-seq takes advantage of the property of enhancers to act bidirectionally, and therefore candidate enhancer fragments cloned downstream of a minimal promoter sequence transcribe themselves in a host cell. A comparison between the final read count of self-transcribed fragments with the initial number of transfected or transduced fragments provides a quantifiable measure of enhancer activity. By assessing candidate enhancer libraries through massively parallel sequencing, researchers have built genome-wide enhancer activity maps (Liu et al. 2017b), functionally validated enhancers identified by other methods such as ChIP-seq (Barakat et al. 2018), ATAC-seq (Wang et al. 2018; Hansen and Hodges 2022), and FAIRE-seq (Chaudhri et al. 2020; Glaser et al. 2021), tested the impact of non-coding variants within or near enhancer sequences (Liu et al. 2017a; Zhang et al. 2018; Kalita et al. 2018; Schöne et al. 2018), and assessed changes in enhancer activity due to external factors such as drug or hormone treatment (Shlyueva et al. 2014; Johnson et al. 2018). Furthermore, Peng and colleagues identified a subset of enhancers in mouse embryonic stem cells that showed activity with STARR-seq but were not associated with active chromatin marks (Peng et al. 2020), suggesting that functional assessment using STARR-seq can reveal novel chromatin-masked enhancers in specific cellular contexts.

Thus, STARR-seq has emerged as a state-of-the-art method for functional interrogation of the non-coding genome.

While STARR-seq is powerful and versatile, there are several challenges associated with successfully carrying out this assay. *First*, the experimental protocol is laborious with over 250 steps, involving construction of a STARR-seq plasmid library, delivery of the library into a host through methods such transfection or transduction, sequencing of delivered (input) and self-transcribed (output) fragments to sufficient read depth, and bioinformatic analysis to identify peaks at enhancer sites (Neumayr et al. 2019). Several of these steps also require preliminary experiments, *for example*, optimizing library construction to achieve sufficient <u>complexity</u>, testing transfection efficiency, mitigating host-specific limitations, and assessing target coverage for reproducible discovery of enhancer location and activity. *Second*, variation across studies and a lack of benchmarking for selecting sequencing and data analysis parameters such as optimal read depth, choice of peak caller, cut-off scores for characterizing enhancer activity, and methods for data validation complicate comparisons across studies. *Third*, most published studies do not provide sufficient details for researchers to adapt or scale their protocols for specific needs such as modifying target library size and choosing enhancer fragment length. Furthermore, lack of quality control (QC) details for critical intermediate steps raises significant concerns for <u>replicability</u> and <u>reproducibility</u> of results from STARR-seq assays.

Several studies have addressed challenges associated with reproducibility in biology, including reports on best practice guidelines for various genomic analysis pipelines, such as RNA-seq (Conesa et al. 2016), ChIP-seq (Landt et al. 2012), and ATAC-seq (Yan et al. 2020), as well as the Reproducibility in Cancer Biology project (Center of Open Sciences and Science Exchange). *For example*, Errington and colleagues attempted 193 experiments from published reports, but were able to replicate only 50 due to failure of previous studies to report descriptive and inferential statistics required to assess effect sizes, as well as a lack of sufficient information on study design. Several of their experiments required modifications to the original protocols with results showing significant deviations from previous findings (Errington et al. 2021). These studies illustrate the difficulties in successfully repeating published functional genomics experiments.

Here, we discuss four major aspects of a STARR-seq assay including *(i)* pre-experimental assay design, *(ii)* plasmid library preparation, *(iii)* enhancer screening, and *(iv)* data analysis and reporting. We also delineate several features specific to STARR-seq in the context of general reproducibility of biological experiments (Freedman et al. 2015). We demonstrate how each of these features vary across previous studies and score them based on details provided in the original publications (**Box 1**). Furthermore, through a series of carefully designed STARR-seq experiments **(Supplemental Materials)** we identify challenges, rate-limiting conditions, and critical checkpoints. Finally, we reanalyze multiple published STARR-seq datasets along with our own data to illustrate limitations of available analysis pipelines. We provide recommendations for study design, library construction, sequencing, and data analysis to help future researchers conduct robust and reproducible STARR-seq assays.

### Different assay goals result in varying experimental designs

The general strategy for STARR-seq consists of cloning a library of fragments selected from either a list of putative enhancers or the entire genome into a STARR-seq plasmid, which is transfected or transduced into a host, such as cultured cells or live tissues. The “input” library and the transcribed “output” fragments from the host is sequenced to high read depth, followed by bioinformatic analysis for quantification of enhancer peaks. In our assessment of published STARR-seq studies, we found that the original protocol (Arnold et al. 2013) was modified in myriad ways to fit various study goals and underlying biological contexts, including altering design features such as target library size, DNA source, fragment length, sequencing platform, choice of STARR-seq vector, and choice of host (**Fig. 1A**). Reporting of these features are crucial for understanding the rationale behind downstream strategies such as protocol scaling, choice of QC measures, and selection of optimal parameters for assay performance and data analysis. Here, we discuss STARR-seq experimental design features, how they vary across studies and affect study reproducibility (**Table 1**), and suggest rationale for selecting different designs **(Supplemental_Table_S1)**.

**Figure 1:**
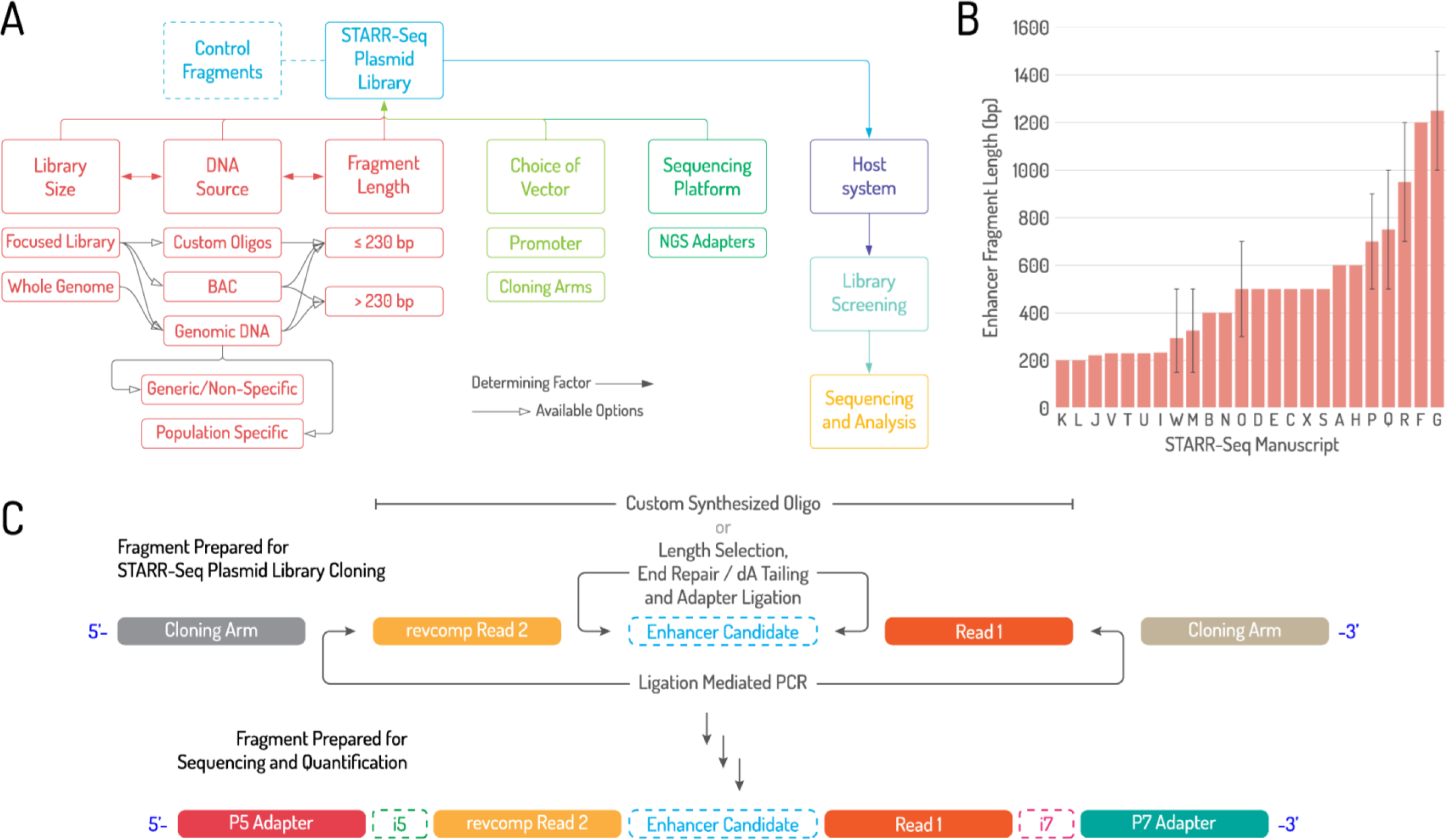
Experimental design features of a STARR-seq experiment. **(A)** Variations of pre-experimental design features and their impact on library preparation protocol are shown. **(B)** Fragment length (mean and range shown) of published STARR-seq libraries reported by de-identified papers A to X on X-axis and reported fragment length on Y-axis. Note that studies W, M, O, P, Q, R and G reported the range for their fragment lengths indicated by the error bars and other studies reported the exact length of the fragment. **(C)** Read architecture schematic of a STARR-seq plasmid library (top) and ‘input’ and ‘output’ sequencing libraries (bottom) are shown.

**Table 1.** Reproducibility features for biological assays with a focus on STARR-seq.

|  | Reproducibility factors for a biological assay | Best practices for reporting reproducibility of a biological assay | Considerations and best practices for reporting STARR-seq specific features to maximize assay reproducibility |
| --- | --- | --- | --- |
| 1 | Experimental Design | • Clear definition of the biological question. | • Biological question that would be addressed using STARR-seq such as genome-wide/targeted screening for enhancer activity, or comparison of enhancer activity due to changes in the host cell environment or due to mutations within or near enhancer sites. |
|  |  | • Rationale for assay design to address the biological question. | • Rationale for choosing specific experimental design parameters including library size, library source, fragment length, vector and basal promoter, sequencing platform, host cell line and library controls. |
| 2 | Protocol | • Details of optimization of protocols, types of kits and reagents used.<br>• Details of protocol scaling, number of reactions used, customizations carried out and data validations. | STARR-seq plasmid library construction<br>• Selection and adaptation of all kits, reagents, and protocols used for insert preparation, library amplification, and confirming sufficient library complexity. Steps such as LM_PCR, library cloning and transformation may require additional optimization to achieve consistency across replicates.<br>• Details of reaction parameters, such as PCR, primer details, number and details of utility of replicates, results of intermediate steps to ensure the protocol is working, and final sample library architecture (with enhancer fragment and sequencing adapters). |
|  |  |  | STARR-seq screening<br>• Optimization of host cell culture conditions, transfection methodology, transfection parameters, RNA isolation and reverse transcription kit protocols and associated modifications, and number of replicates required for each step for successful representation of all target sites, and report potential issues accounted for each step and mitigation strategies used (such as fragment loss).<br>• Reaction details (such as reverse transcription), replicates used, primer details, and results for intermediate QC steps.<br>• For libraries using UMI's, details on UMI ligation strategy and illustration of sequence architecture with UMI position. |
|  |  |  | Sequencing<br>• Optimization of parameters for sequencing PCRs to prevent over-amplification, details on selection of multiplexing strategy used, and library pooling. |
|  |  |  | • Details of sequencing parameters such as read length, single- or paired-end, and read depth based on library size and coverage, and read loss due to filtering was accounted.<br>• Complete sequencing information including experimental and sequencing run parameters and read details. |
| **3** | **Quality Control** | • Sample clean-up and purity assessments.<br>• QC and validation of intermediate steps.<br>• QC and validation of results. | STARR-seq experimental protocol<br>• Sample clean-up followed by concentration and purity after each experimental step for all replicates.<br>• Assessment of fragment length distribution for insert and vector prior to cloning to check for primer or adapter dimers and fragments of irregular which may lead to PCR or cloning biases.<br>• Assessment of transformation efficiency and library complexity through CFU assessment by plating of serially diluted transformant and control plasmids.<br>• Optimization of transfection efficiency, host cell line immune responses and RIN analysis of isolated RNA.<br>• Assessment of final fragment distribution of libraries prior to sequencing.<br>STARR-seq data processing and analysis<br>• Processing and filtering steps for raw reads, including sequence mapping and deduplication.<br>• Assessment of read depth and library coverage as well as read quality across replicates.<br>• All read and data QC parameters used to enable replication of the analysis. |
| **4** | **Data analysis** | • Analysis pipeline based on biological question and experimental parameters.<br>• Thresholds and cut-offs for data validation and statistical inferences. | • Read processing steps used for sequence mapping, filtering of bad or unmapped reads, and read deduplication methodology (depending on the use of UMIs)<br>• Peak caller to analyze enhancer activity.<br>• Peak validation method and assessment of control regions for data QC.<br>• Detailed pipelines for replication of complete analysis to ensure data reproducibility. |
| **5** | **Data reporting** | • Experimental design features, protocol, and analysis pipelines.<br>• Intermediate steps to check if the protocol is working and final data. | • Details of library design and pre-experimental optimization steps.<br>• Experimental method with scaling guidelines, protocol parameters, and analysis pipeline.<br>• Details of quality assessment at critical stages of the protocol, and final raw and processed data. |

### Target library size

The versatility of STARR-seq allows for enhancer screening across both whole genomes and targeted regions. Studies screening for enhancer activity across numerous targets typically require multiple reactions for all library preparation steps, and high cloning and transformation efficiencies to achieve adequate representation of all fragments compared to studies with fewer targets. *For example*, a library generated by Johnson and colleagues to assess the effect of glucocorticoids on genome-wide enhancer activity contained over 560 million unique fragments (Johnson et al. 2018). The authors performed 60 reactions of sequencing adapter ligations and 72 transformations and grew the pooled libraries in 18-liter LB broth. In contrast, Klein and colleagues synthesized a targeted library of 2,440 unique fragments along with customized adapter and cloning sequences in three reactions to compare context-specific differences among seven different MPRA designs, including two variations of STARR-seq (Klein et al. 2020).

Cloning and transformation of each tested design was performed in duplicates, and each reaction was grown in 100 ml of LB broth. While adequately scaled protocols exist for whole genome STARR-seq in *Drosophila melanogaster* and humans (Neumayr et al. 2019; Arnold et al. 2013), scaling guidelines for focused libraries are not well reported since their sizes can vary from >7 million unique fragments (Wang et al. 2018) to a few hundred (Vockley et al. 2015) or thousand fragments (Klein et al. 2020). We provide rationale for custom designing STARR-seq assays in **Supplemental_Table_S1** and scaling guidelines based on library size, fragment length, and read coverage in **Supplemental_Table_S2**.

### DNA source

A strong advantage of STARR-seq is its ability to screen random fragments of DNA from any source for enhancer activity. To this effect, DNA can be sourced from commercially available DNA repositories (Liu et al. 2017b), from specific populations carrying non-coding mutations or SNPs to be assayed (Liu et al. 2017a), cultured bacterial artificial chromosomes (BACs) (Arnold et al. 2013), or custom oligo pools (Kalita et al. 2018). Different sources have their own advantages and drawbacks that are important to consider for efficient library building **(Supplemental_Table_S1)**. For example, custom synthesized oligo pools are generally designed to be of uniform length and may include sequencing and cloning adapters as part of the design.

These libraries may not require fragment length selection and adapter ligation and are also devoid of any representation bias caused by fragments of non-uniform lengths. However, we note that synthesized fragments may also have sequence-specific biases due to factors such as repetitiveness, single stranded DNA structures, and mappability issues associated with the underlying sequence (Halper et al. 2020). Furthermore, most STARR-seq studies using synthesized oligos have a fragment synthesis-based length restriction of 230 bp; though recent advances can allow up to 300 bp. Therefore, current limitations for custom-made fragments include synthesis-based length restrictions, higher costs, and a smaller library size as opposed to larger focused or whole genome libraries using genomic DNA. Of note, genomic DNA derived from host cell is recommended for focused studies that aim to assess enhancers captured through chromatin-based techniques such as ATAC-STARR-seq (Wang et al. 2018), ChIP-STARR-seq (Barakat et al. 2018) and FAIRE-STARR-seq (Chaudhri et al. 2020). Using the host cell DNA for these methods enables selection of cell-type specific candidate enhancers which can then be functionally validated with STARR-seq upon re-delivery into the host. While in theory any DNA fragment can be tested in any cell line, this recommendation can be advantageous to test cell-type specific effects.

### Fragment length

Length of each candidate enhancer fragment that is compatible with the assay goal, library size, and DNA source (**Fig. 1A**) determines critical experimental parameters such as the extension temperature of ligation-mediated PCR (LM_PCR), sample volume-to-bead ratio for clean-up, insert-to-vector ligation ratio, and fragment molarity for cloning reactions. Most STARR-seq studies either use ∼500 bp fragments sourced from sheared whole genome DNA or DNA isolated from BACs, or use ≤230 bp oligo pools (**Fig. 1B**). However, an ‘ideal’ fragment length is dependent on the assay goal, since different fragment lengths have different benefits and limitations **(Supplemental_Table_S1)**. Fragments >500 bp may be more economical and better suited for genome-wide screens or enhancer discovery within larger targets. Notably, larger fragments may not have the resolution to detect the activity of individual enhancers but are useful to identify the compounding effect of multiple closely-located enhancers or the influence of flanking sequences on transcription factor binding (Schöne et al. 2018; Klein et al. 2020). In contrast, focused studies using shorter fragments allow for fine mapping the enhancer effects of individual TF binding sites but may not uncover synergistic effects detectable by longer fragments. Interestingly, Klein and colleagues compared the effects of fragment length on enhancer activity by extending the flanking genomic sequences of the same candidate enhancer sites to create 192 bp, 354 bp, and 678 bp fragments using hierarchical multiplex pairwise assembly (Klein et al. 2020). The authors found less concordance in enhancer activity for the same candidate enhancers when fragments of different lengths were compared, with higher correlation between smaller fragments than larger fragments. Schöne and colleagues demonstrated that the flanking sequences have an effect on DNA shape that affect TF binding, thus impacting enhancer activity (Schöne et al. 2018). Hansen and Hodges reported that differently sized fragments around the same genomic region can identify distinct active regions (Hansen and Hodges 2022). Library preparation from sheared DNA require accurate size selection methods as fragments of non-uniform length are more prone to uneven representation in the final library due to PCR and cloning biases. In fact, Neumayr and colleagues suggest keeping the fragment lengths within ∼300 bp of each other to avoid extreme biases (Neumayr et al. 2019). If the assay requires variable fragment lengths, then similarly-sized fragments can be batched together in reactions and pooled prior to enhancer screening.

### Choice of vector and basal promoter

The choice of STARR-seq vector and basal promoter primarily depends on the host type, the class of candidate enhancers being assayed, and the biological question. The original STARR-seq vector included a backbone sequence derived from vectors previously used for reporter assays or MPRA, and featured a species-specific minimal or core promoter i.e., a promoter that binds a set of general transcription factors and RNA Polymerase II to initiate transcription (Haberle and Stark 2018). While most studies have continued to use a similarly designed vector, the backbone has been modified to fit different biological contexts. *For example*, Inoue and colleagues developed LentiMPRA, that included candidates cloned upstream of the minimal promoter on a lentiviral construct. This allows the enhancer-promoter sequence to get integrated into the host genome, thus providing a genomic context to the assay (Inoue et al. 2017). Lambert and colleagues adapted a recombinant Adeno Associated Virus (AAV) vector to enable library delivery into mouse retina and brain (Lambert et al. 2021). Muerdter and colleagues observed that enhancer signals from the original human STARR-seq minimal promoter, SCP1 (Super Core Promoter 1) were not consistent due to interference from the ORI promoter that was present on the same vector. The authors mitigated this issue by redesigning the vector to carry just the ORI promoter and demonstrated a more robust assay (Muerdter et al. 2018). Sahu and colleagues used a minimal δ1-crystallin gene (Sasaki) promoter for their experiments across multiple libraries.

They also used a CpG-free vector backbone containing the Lucia reporter gene driven by EF1α promoter to assess the effect of CpG methylation on enhancer activity (Sahu et al. 2022). Klein and colleagues compared seven different MPRA and STARR-seq designs and concluded that LentiMPRA and the ORI vector showed the highest consistencies in activity across replicates (Klein et al. 2020). Thus, the vector can be designed based on the assay goal and modified to be compatible with the host genome.

Studies have also shown that core promoters are not only specific to a particular species, but also specific to different co-factors (Haberle et al. 2019), genomic environments (Hong and Cohen 2022), and enhancers targeting genes of particular function (Zabidi et al. 2015). For example, Zabidi and colleagues performed seven whole genome STARR-seq assays in different *Drosophila* derived cell lines using different core promoters and found two distinct promoter classes, housekeeping and developmental, based on promoter-enhancer specificities (Zabidi et al. 2015). Jores and colleagues demonstrated that the composition of core promoter elements and the presence of distinct TF binding sites determined the basal strength of the promoter as well as its specificity to different enhancers (Jores et al. 2021). In fact, low strength of basal promoters adds limitations to measuring repressive functions of enhancers (Tewhey et al. 2016). Recent MPRA studies have also reported automated methods for design and synthesis of non-repetitive promoter sequences (Hossain et al. 2020) and proposed models to predict transcription initiation rates for different promoters (LaFleur et al. 2022). Martinez-ara and colleagues tested pairwise enhancer-promoter combinations in mouse embryonic stem cells and reported that enhancer-promoter pairs targeting housekeeping genes show more consistent activity than pairs targeting other genes (Martinez-Ara et al. 2022). Bergman and colleagues also tested pairwise combinations of 1000 enhancers with 1000 promoters in human K562 cells using ExP-STARR-seq (Bergman et al. 2022). However, they found that promoters were activated uniformly by most enhancers, with enhancer-promoter pairs for housekeeping genes only showing subtle differences compared to other tested combinations. Barakat and colleagues tested their ChIP-STARR-seq libraries with multiple promoters including SCP1, CMV and AAV on human embryonic stem cells, and they did not find significant differences between the promoters (Barakat et al. 2018). While core promoters may have specificity to certain enhancers, there is still ambiguity in our understanding of promoter-enhancer interactions. Therefore, variable effects of a basal promoter must be considered when designing MPRA or STARR-seq assays (Mulvey et al. 2021).

### Control fragments

The plasmid library can also be designed to include control sequences to validate STARR-seq results. Commonly used control sequences include fragments previously validated to be active (positive control) or inactive (negative control) by MPRA or classical reporter assays (Klein et al. 2020), or ‘scrambled’ sequences can be used as negative controls (Martinez-Ara et al. 2022). While currently available whole genome STARR-seq datasets may serve as a repository for shortlisting control sequences, factors such as cell-type and enhancer-promoter specificities make it difficult to identify controls specific to a cell type or minimal promoter. To this end, Neumayr and colleagues suggest running a small-scale focused STARR-seq screen prior to an experiment to identify potential control regions (Neumayr et al. 2019). Alternatively, studies have also used regions predicted to be inactive, such as coding sequences of genes (Arnold et al. 2013) or CTCF-binding regions (Vanhille et al. 2015), as a proxy for negative controls.

### Sequencing methodology

The final set of considerations for library design include the choice of sequencing platform and the compatible adapters and kits required for library preparation. Almost all STARR-seq libraries use Illumina-compatible designs, where the putative enhancer fragment is flanked by two adapters consisting of read 1 and read 2, followed by unique index sequences or barcodes (i5 and i7), and P5 and P7 sequences to facilitate the sequencing-by-synthesis method (**Fig. 1C**). The two adapters flanking the candidate fragment serve as “constant” sequences for primer annealing during library amplification as well as primer recognition sites during Illumina sequencing.

Cloning overhang arms are designed based on the selected STARR-seq vector and are ligated adjacent to the adapters through LM_PCR, and used for library cloning. The overhangs are subsequently replaced by the index barcodes i5/i7 and adapters P5/P7 to facilitate sequencing (**Fig. 1C, Supplemental_Fig_S1)**. These adapter sequences are especially important for designing oligo libraries, LM_PCR primers, and blocking oligos during hybridization and capture of focused libraries.

### Roadmap to successful library construction

With all the design details in place, construction of a STARR-seq plasmid library includes several optimization and QC steps with each step contributing to varying levels of assay reproducibility. Here, we highlight key elements of the protocol that strongly influence the outcome and offer clarity and explanations behind various optimizations. The features that impact library building include insert preparation, vector linearization, library cloning, and library amplification (**Fig. 2**). Additionally, focused libraries also require a library enrichment step to capture the fragments of interest. This enrichment can be done prior to insert preparation using ATAC-seq (Wang et al. 2018), ChIP-seq (Barakat et al. 2018), FAIRE-seq (Chaudhri et al. 2020), fragments extracted from specific BACs, or fragments synthesized as custom oligo pools. Alternatively, library enrichment can also be carried out after insert preparation using techniques like capSTARR-seq, which uses special microarrays (Vanhille et al. 2015), or by using custom hybridization and capture probes (Liu et al. 2017a).

**Figure 2:**
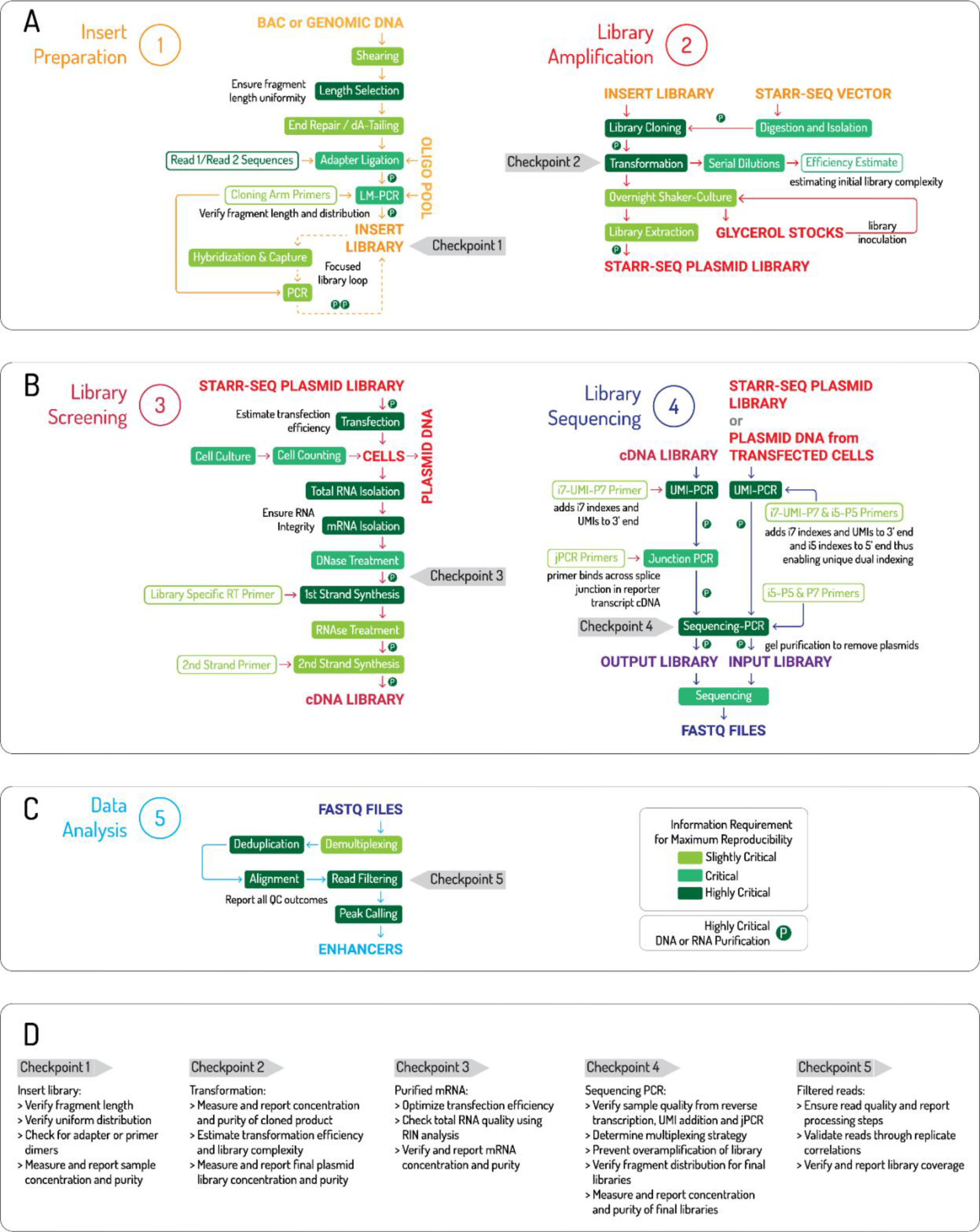
Roadmap to a successful STARR-seq assay: Schematic illustrates five major sections of STARR-seq **(A)** Insert preparation and Library amplification, **(B)** Library screening and library sequencing, **(C)** Data analysis. Individual steps in each section are categorized as **slightly critical (**light green**), critical (**green**), or highly critical (**dark green) based on the importance of reporting the methodological detail and intermediate results for a reproducible assay. **(D)** Assay checkpoints. Each section also has a checkpoint, which may serve as stopping points in the protocol to perform validation of previous steps prior to moving to the next step. Key QC measures that may be carried out at these checkpoints are also provided in the bottom panel. Of note, Panel B, section 4: ‘Library sequencing’ illustrates a methodology for adding Unique Molecular Identifiers (UMIs) and Unique Dual Indexes. A more detailed schematic along with sequence information is provided in **Supplemental_Fig_S1** and **Supplemental_Table_S3**. Alternative methods for UMI addition have also been reported in several studies.

While these steps typically follow kit-based protocols, there are limited scaling guidelines for differently sized libraries. To ensure replicability of the scaled protocols, details such as the number of reactions performed, sample concentration and purity, details of the purification method used for intermediate product stages, use of control reactions, and the experimental parameters must be reported. Additionally, reporting details of validations for intermediate steps such as length verification of fragments following length selection as well as adapter and cloning arm ligation steps provide checkpoints for researchers attempting similarly designed experiments. It is often difficult to understand whether the library being constructed is successful or not without sequencing the library, which is both time consuming and expensive, and it is also hard to detect and pinpoint experimental errors due to the length of the protocol.

Therefore, having validation checkpoints at intermediate stages help address these problems. While performing STARR-seq, we shortlisted five such checkpoints in the protocol that require product assessment and validation before proceeding to the next step (**Fig. 2**). We also categorized intermediate assay steps as ‘highly critical’, ‘critical’ and ‘slightly critical’ based on the requirements of data reporting to replicate those steps (**Fig. 2**).

The final determinant of library coverage is the efficiency of molecular cloning and bacterial transformation. Cloning strategies such as In-fusion HD, Gibson assembly, and NEBuilder HiFi DNA Assembly allow for fast and one-step reactions that use complimentary overhang sequences on the inserts and the vector. While these methods have comparable accuracy and efficiency, the chosen method needs to be optimized prior to cloning the final library pool. The critical parameters to consider, optimize, and report include *(i)* purity and concentration of the insert and the vector, *(ii)* insert-to-vector ratio by mass or moles and the maximum molarity per reaction supported by the cloning enzyme, *(iii)* the number of cloning reactions required and pooled, *(iv)* details of negative control used for cloning (typically using a linearized vector-only reaction) *(v)* competent cell strain and volume per reaction, *(vi)* the amount of cloned product transformed per reaction and the total number of reactions performed and pooled, *(vii)* details of positive (such as supercoiled plasmids like pUC19) and negative (such as un-ligated, linearized vector) controls used for transformation, *(viii)* the volume of culture used to grow the library, and *(ix)* the number of unique colonies observed in a dilution series for estimating transformation efficiency and library complexity. *For example*, Barakat and colleagues used genomic fragments obtained from chromatin immunoprecipitation to perform multiple PCR reactions and generate inserts for cloning via Gibson assembly. Each reaction was performed in duplicate, which were pooled prior to transformation in DH10β cells. The transformed product was grown in SOC medium, serially diluted, plated, and grown in LB to obtain between 8 and 31 million unique colonies (Barakat et al. 2018). In contrast, Kalita and colleagues used synthesized oligos for two rounds of nested PCRs to obtain the inserts for cloning using Infusion HD strategy, followed by multiple transformation reactions (Kalita et al. 2018). For each of these steps, the authors provided the exact volumes for each reaction, elution steps, the number of reactions used, and the number of colonies obtained from the dilution series, enabling accurate replication of these experiments.

Prior to final library transformation, pilot transformations may also be carried out using a positive control plasmid and the cloned library to estimate the transformation efficiency by calculating the CFU/μg (Colony Forming Units per microgram of plasmid). This step helps estimate the number of transformations required to achieve the required complexity **(Supplemental_Table_S2)**. The final complexity of the plasmid library is typically assessed by evaluating sequencing read representation of the target regions. *For example*, Johnson and colleagues performed MiSeq and used the preseq tool (Daley and Smith 2013) to assess library complexity for their whole genome library (Johnson et al. 2018). The verified STARR-seq plasmid library is then transfected or transduced into the host for screening. For sequencing, candidate fragments need to be re-amplified from the plasmid pool and fitted with complete P5 and P7 adapters and index barcodes replacing the cloning arms (**Fig. 1C**). To compare self-transcribed transcripts (output) with fragments present in the plasmid library (input), two sets of sequencing libraries (output and input) are prepared. While there is a consensus on strategy for building the output library, the input library preparation varies across studies. *For example*, some studies built the input library by directly amplifying the plasmid pool in a second round of LM_PCR, independent of library transfection (Muerdter et al. 2018; Neumayr et al. 2019). Other studies have isolated the input fragment DNA from a fraction of the transfected cells (Arnold et al. 2013; Peng et al. 2020) or extracted DNA and RNA from the same cells to account for fragment loss during transfection (Klein et al. 2020). An advantage of the latter strategy is that it helps identify fragments that were retained following library delivery, thus allowing for discriminating inactive fragments from fragments lost during library delivery. This is especially useful for hosts with lower transfection or transduction efficiency causing high fragment loss.

### Enhancer screening using STARR-seq

Enhancer screening involves preparation of the STARR-seq ‘output’ library followed by deep sequencing of the input and output libraries in parallel. The output library is derived from STARR-seq transcripts generated by functional enhancers when delivered into a host, chosen based on the desired cellular environment. For studies using cultured cells, the choice of host cell line determines multiple transfection parameters such as *(i)* transfection methodology, *(ii)* amount of plasmid per transfection, *(iii)* the total number of cells transfected per replicate, and *(iv)* incubation time of transfected cells. These details can vary across studies and determine downstream experimental features for screening. *For example*, for focused STARR-seq, Wang and colleagues electroporated 120-130 million Lymphoblastoid Cell Lines (LCLs) per replicate at 5 µg per one million cells, and used five replicates for their library comprising of >7 million unique fragments (Wang et al. 2018). Kalita and colleagues also electroporated LCLs, but used 3 µg per 7.5 million cells per replicate in nine replicates for their library of 75,501 unique fragments (Kalita et al. 2018). For whole genome STARR-seq, Sahu and colleagues transfected 1 µg of their library per one million cells and used two replicates comprising 35-500 million cells per replicate in multiple cell lines (Sahu et al. 2022). In contrast, Lee and colleagues electroporated 8 µg of their whole genome library per one million cells and used two replicates of 700 million to 1 billion cells per replicate in multiple cell lines (Lee et al. 2020). Therefore, it is important to consider the exact cell line-specific transfection method and its nuances. For instance, transfection through electroporation is more efficient for harder-to-transfect cells such as Embryonic Stem Cells (ESC) or LCLs and involves shorter incubation time of about 6 hours. However, this process requires higher amounts of plasmid and more cells per library due to the loss of cells during the electroporation process. Methods using lipofectamine may be recommended for various mammalian cell lines such as HEK293T, as they are more efficient and gentler on the cells, but this procedure involves longer incubation times of 12 to 24 hours. Furthermore, higher transfection efficiency reduces the need for multiple transfections for each biological replicate. Another important consideration for selecting the cell line is to test whether there is a need for kinase inhibitors to prevent interferon response from the host. Muerdter and colleagues demonstrated that cell lines such as HeLa S3 trigger a type I interferon (INF-1) response upon library transfection which disrupts enhancer signals and leads to false results. The authors suggest the use of kinase inhibitors BX-795 hydrochloride to inhibit TBK1 and Imidazolo-oxindole PKR inhibitor C16 to inhibit PKR to mitigate the situation (Muerdter et al. 2018). Our assessment of recent studies showed enrichment of interferon gene motifs within enhancer peaks, illustrating the significance of interferon response on specific cell types (Johnson et al. 2018; Sahu et al. 2022). While STARR-seq and most MPRAs have predominantly been performed on cultured cells, some studies have also used *in-vivo* or *ex-vivo* systems such as mouse retina explants (White et al. 2013), cortex (Shen et al. 2016), and post- natal mouse brain (Lambert et al. 2021). Here, considerations include the mode of library delivery such as injections or electroporation (Montana et al. 2011) and transduction efficiency for AAV-based libraries.

The factors that determine the extent and scalability of enhancer screening include library size and the number of biological replicates. The library size determines the multiplicity of reactions required per biological replicate. Here, the term “reactions” indicate smaller experimental setups that are pooled for a single large experiment and “replicates” typically refers to additional experiments conducted for validation and reproducibility. While reactions may be pooled at intermediate steps of the protocol, replicates may only be pooled post-hoc during peak calling. The impact of the number of replicates on final output has been assessed previously. *For example*, Hansen and Hodges found that an increase in the number of replicates increased enhancer calls in their ATAC-STARR-seq assays (Hansen and Hodges 2022). However, the authors also caution that increasing replicates may also increase the number of false positive calls. Thus, it is important to consider the number of replicates especially for studies that pool reads from replicates for peak calling.

Following library delivery, the next series of steps involve total RNA isolation, mRNA isolation, and reverse transcription of STARR-seq transcript-specific mRNA using a reporter-specific primer for first strand synthesis to build a cDNA library. A majority of studies have used well-described kits for these steps and follow standardized scaling guidelines. However, in order to replicate these steps, authors should report reaction parameters like sample concentrations, amount of starting material for each step, replicate information, and sample pooling details to ensure replicability of the steps. Additional contributors to reproducibility here include intermediate sample validations and QC assessments including RNA integrity (RIN) analysis and RNA purification using TURBO DNase and RNAClean XP beads, all of which increase library quality. The cDNA library is then amplified using junction PCR primers to remove plasmid artifacts followed by ligation of sequencing adapters and index barcodes using LM_PCR, similar to the input sequencing library (**Fig. 2**). An important consideration here is to prevent overamplification of the library, which leads to sequencing bias, by adjusting the number of amplification cycles during the final PCR step. Sequencing libraries may be visualized on an agarose gel for a characteristic “*smear*” at the expected fragment length as opposed to a tight band (Neumayr et al. 2019), unless the library is built using synthesized fragments of uniform length.

While preparing the screening library is simpler compared to the plasmid library, certain anomalies may occur that hamper library quality and replicability. For example, batch effects during library preparation can be minimized by constructing biological replicates independently and on different days. For example, Klein and colleagues transfected each replicate on separate days and prepared the libraries independent of each other (Klein et al. 2020). Another common occurrence in STARR-seq data is the presence of PCR duplicates due to the use of PCR-based sequencing library preparation methods. Self-transcribed mRNA may resemble these PCR duplicates, leading to data anomalies. Previous studies have typically removed PCR duplicates by filtering reads originating from a single fragment of DNA using computational techniques such as the “*MarkDuplicates*” function in Picard (Peng et al. 2020). However, this may not be ideal as it runs the risk of also removing STARR-seq transcripts, thus resulting in reduced enhancer signals and high false negatives (Liu et al. 2017b). Therefore, for accurate removal of PCR duplicates, Kalita and colleagues added Unique Molecular Identifiers (UMIs) during the reverse transcription step, such that PCR duplicates possess the same UMI sequence, unlike the self-transcribed mRNA (Kalita et al. 2018). There are now several published variations of the UMI method. For instance, Neumayr and colleagues suggested adding UMIs following an extra second strand synthesis step for cDNA in a single cycle PCR reaction (Neumayr et al. 2019).

Any UMI-based STARR-seq experiment can use relevant UMI-based read deduplication algorithms such as UMI-tools (Smith et al. 2017), Picard’s “*UmiAwareMarkDuplicatesWithMateCigar*” function (Broad Institute. 2019), or the calib library (Orabi et al. 2019).

The final step is sequencing of the input and output libraries. Libraries are usually pooled and sub-sampled prior to sequencing. While these are standardized procedures, small variations in these steps can largely impact data reproducibility. *For example*, while running multiple STARR-seq samples on the same lane of an Illumina-based sequencer, a phenomenon called ’index hopping’ may occur, where index barcodes are assigned to the wrong libraries leading to inaccurate sequencing data (Kircher et al. 2012). To mitigate this, each library can be assigned two indexes or unique dual indexes (UDI) to detect these incidents and increase the accuracy of <u>demultiplexing</u> individual libraries from a library pool (**Fig. 1C**, **Fig. 2**) (MacConaill et al. 2018).

### STARR-seq analysis and reporting

Despite strictly following experimental design and protocols for reproducing an assay, lack of data analysis guidelines can result in inconsistent findings. The main features that contribute to reproducibility here include sequencing depth, read processing, and read QC, as well as choice of peak caller, cut-offs used for analysis, data QC, and validations. While there are well described guidelines and data standards set for sequencing techniques such as RNA-seq (Conesa et al. 2016), ChIP-seq (Landt et al. 2012), Hi-C (Lajoie et al. 2015) and ATAC-seq (Yan et al. 2020), data analysis for STARR-seq is not standardized.

### Sequence depth, coverage, and library complexity

A major factor for any high-throughput study is to attain sufficient coverage of the sequenced library. *For example*, the Sequence QC project (SEQC) demonstrated that read depth and choice of analysis pipelines are key aspects for reproducibility of RNA-seq experiments (Su et al. 2014). Read depth and coverage requirements for techniques including ChIP-seq (Landt et al. 2012) and RNA-seq (Tarazona et al. 2011; Sims et al. 2014; Conesa et al. 2016) have been thoroughly discussed, and multiple data standards have been set by consortiums such as ENCODE (Birney et al. 2007). On the other hand, STARR-seq read depth requirements are vague. Even calculating library coverage of STARR-seq data using available RNA-seq guidelines results in inaccurate estimates and faulty inferences of current studies.

Whole genome STARR-seq assays typically have lower coverage due to technical limitations, although these assays are highly useful for screening entire genomes for strong and distinct enhancer signals. *For example*, Johnson and colleagues obtained 59X coverage of the human genome in their library (Johnson et al. 2018) while Liu and colleagues reported that 74.3% of their library were covered by at least 10 reads (Liu et al. 2017b). Whole genome studies, which primarily focus on mapping the entire enhancer landscape of the organism, may not require a high-resolution view of enhancer activity. In contrast, smaller focused libraries such as those quantifying the effect of non-coding mutations require a higher coverage to reduce false negative signals. *For example*, Schone and colleagues obtained over 100X coverage of the glucocorticoid receptor binding sites they assessed for activity (Schöne et al. 2018). Hence, there needs to be an established minimum sequencing depth requirement for STARR-seq experiments and guidelines for reporting read length, number of sequenced reads per library, and sequencing parameters used to generate reproducible data. Computational tools such as preseq (Daley and Smith 2013) can be used to estimate the number of sequenced reads required to obtain sufficient coverage for all the unique fragments present in a sequencing library, but they require a preliminary “shallow” sequencing run of the STARR-seq plasmid library. Based on the preliminary run, preseq estimates a library complexity curve that can be used to determine the efficacy of deeper sequencing runs. Alternatively, the required number of reads can also be estimated from the library size, required library complexity, and expected <u>dynamic range</u> (i.e., range of enhancer activity) of the library, as shown in **Supplemental Table_S2**.

There is also a large variation in how sequence depth is reported by different studies. *For example*, some studies have reported the percentage of unique library fragments with at least ‘N’ reads as a measure of sequencing depth, while others have reported the average number of reads that aligned to all library fragments. We suggest reporting both these metrics since they each summarize sequencing depth distinctly; the former presents an atomistic view of read coverage of the fragment whereas the latter indicates library coverage as a whole.

### Read processing and quality control

After completion of sequencing, raw reads generated from the sequencer are first demultiplexed and the correct sample labels are assigned. Reads are then mapped to a reference genome and filtered to remove unmapped reads, off-target reads, PCR duplicates, and reads with low mapping quality based on the assigned Mapping Quality (MAPQ) scores. Read filtering results in the loss of a large percentage of reads and reduces the overall coverage of the library. Loss of coverage will reduce the strength of enhancer signals and may result in incorrect quantification of target regions*. For example*, Johnson and colleagues reported up to a 40% loss of their reads after QC filtering (Johnson et al. 2018). To compare the read loss statistics and QC metrics across studies, we analyzed multiple published STARR-seq datasets in addition to our own using a custom analysis pipeline **(Supplemental Methods)**. In our assay, which included a modified sequencing library design to include UMIs and UDIs (**Fig. 3A, Supplemental_Fig_S1)**, we filtered out ∼75% of all reads after QC (**Fig. 3B, Supplemental Methods)**. Since the reanalyzed datasets did not contain UMIs, precluding the use of our deduplication strategy, we instead modified our pipeline to use Picard (Broad Institute. 2019) to remove duplicate reads. We observed that the percentage of reads lost after read QC for focused STARR-seq libraries ranged between 70 to 80%, while whole genome libraries lost approximately half as much **(Supplemental_Fig_S2)**. We also observed that the percentage of reads lost due to deduplication was significantly lower in our study compared to the reanalyzed data. It is possible that Picard’s deduplication strategy inadvertently eliminated self-transcribing STARR-seq fragments along with PCR duplicates. Therefore, it is not only important to consider read loss after sequencing when designing the experiment, but also to report read QC steps such as filtering parameters and cut-offs used, on public repositories such as GitHub to enable replication of the analysis.

**Figure 3:**
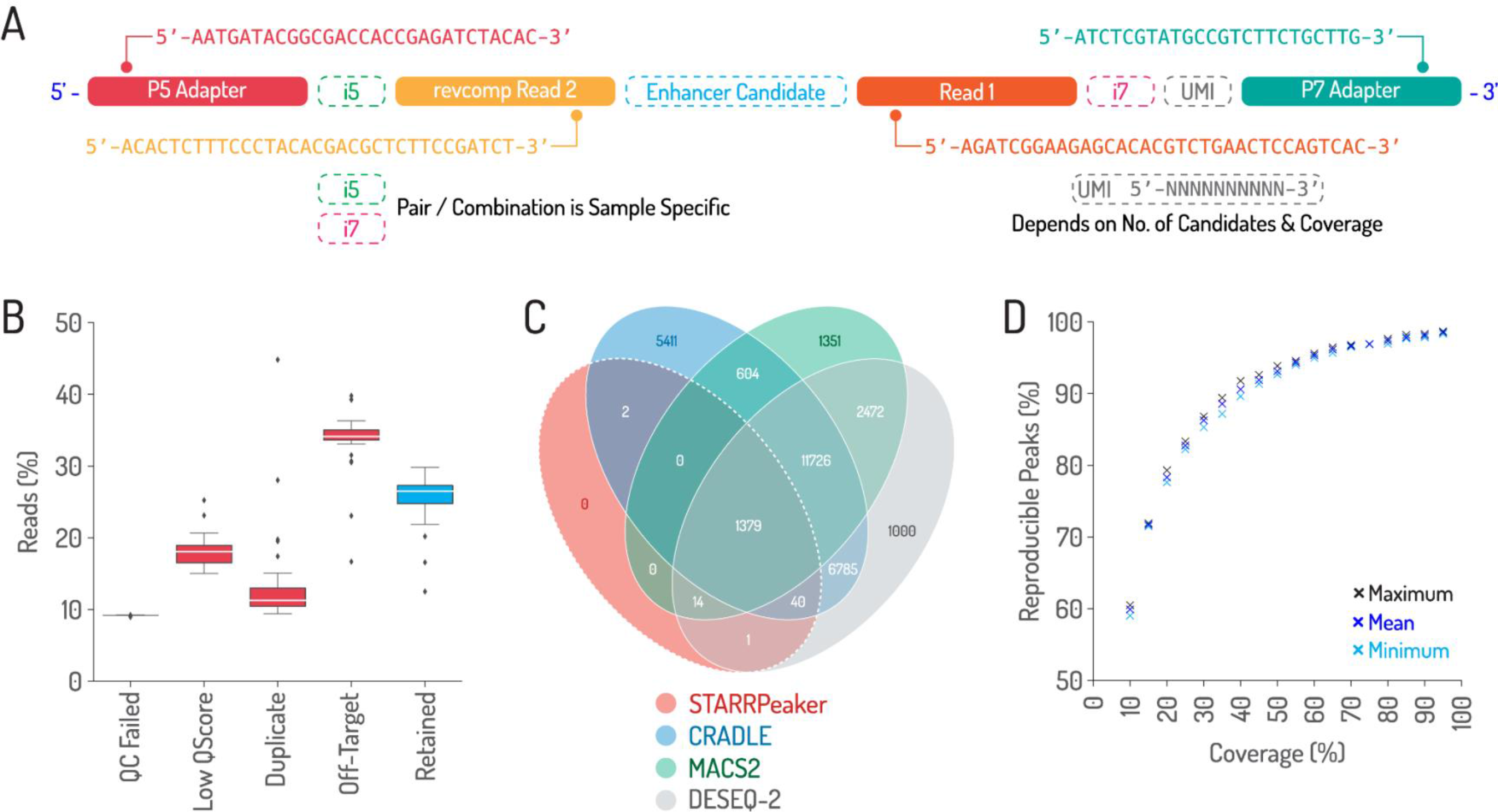
Read architecture, read depth loss, and peak calling comparisons (A) Schematic illustrating read architecture of final STARR-Seq sequencing library (input and output) prior to sequencing. (**B)** Box plots show the percentage of STARR-Seq reads retained and those removed due to various QC filters observed during analysis of in-house STARR-seq assays. (**C)** Overlap of STARR-Seq peaks identified from the same experiment by four different callers. (**D)** Percentage of peaks that were reproduced from in-house STARR-Seq dataset after down- sampling to various coverage thresholds. Analysis details provided in **Supplemental Methods**.

After ensuring read quality, studies often evaluate data quality and reproducibility by calculating correlation coefficients for read counts between library replicates. Some studies also calculate correlation of fold change between read counts of RNA from output compared to DNA from input libraries, across sample replicates to check for consistency of STARR-seq activity.

Our analysis of published datasets and our own data showed strong correlations between library replicates **(Supplemental_Fig_S3, Supplemental_Fig_S4, Supplemental Methods)** similar to correlations (Pearson’s correlation coefficients >0.8) reported by other STARR-seq studies.

Correlation values can vary across studies depending on *(i)* the type of correlation metric (Pearson or Spearman) and type of correlation measured (input versus input, output versus output, or output/input versus output/input), *(ii)* normalization of reads (RPM, RPKM or not normalized), and *(iii)* the read scaling factor used (log scale or not scaled). Additionally, data quality can be assessed for batch effects using Principal Component Analysis (PCA) for all library replicates, where clustering reflects library consistency. For instance, PCA of our libraries showed strong clustering of library replicates and were consistent with the reanalyzed datasets from published studies **(Supplemental_Fig_S5, Supplemental Methods)**.

### Enhancer calling

Following data QC assessment, the next step involves comparing the input and output libraries to quantify enhancer activity. Here, read counts for candidate regions are compared between the output and input libraries to determine a ‘peak’ of concentrated reads that are indicative of enhancer activity. Previously, peak calling was made using either the MACS2 algorithm which was originally developed for ChIP-seq analysis (Arnold et al. 2013), the DE-seq2 algorithm (Love et al. 2014), or enhancer activity was detected by calculating statistically significant fold-change in reads per region between the input and output libraries. Recently, STARR-seq-specific peak calling algorithms such as STARRPeaker (Lee et al. 2020) and CRADLE (Kim et al. 2021) have also been developed. The primary difference between the peak calling algorithms lies in the probabilistic read density distribution used to estimate expected STARR-seq read density across the regions of interest. MACS2 and CRADLE use a Poisson distribution to model region-wide read density whereas DESeq2 and STARRPeaker use negative binomial distribution.

Additionally, CRADLE and STARRPeaker use regression models to account for sequence-based biases before modelling the read density distribution. We also note that each of these callers have distinct utilities based on the data obtained and the assay design. For example, CRADLE may be better suited for studies with low correlation and irregular clustering across replicates, as it takes variance between replicates into account prior to merging reads for peak calling.

To assess reproducibility of enhancer calls, we used four methods on our STARR-seq data and assessed overlap of peaks between callers (**Fig. 3C, Supplemental Methods)**. While peaks called by STARRPeaker overlapped with all other peak callers, the number of peaks varied across different callers, suggesting higher experimental noise than enhancer signal for a subset of enhancers. Akin to other next-generation sequencing experiments, signal-to-noise ratio in STARR-seq can be measured by evaluating the ratio between the mean and the standard deviation of enhancer activity values across multiple replicates. Furthermore, STARR-seq data might also be confounded by DNA sequence related biases which cannot be removed solely by increasing the number of replicates (Kim et al. 2021). Thus, we recommend using STARR-seq specific callers such as STARRPeaker and CRADLE that can identify and eliminate sequence-associated biases.

To investigate the effect of read coverage on peak calling, we sub-sampled our reads and used STARRPeaker to call peaks to delineate the percentage of peaks retained at different levels of sequencing coverage (**Fig. 3D, Supplemental Methods)**. We observed a steady increase in peaks with increase in coverage up to 280X, demonstrating the importance of read depth for STARR-seq. *For example*, we found that only 60% of the peaks remained at ∼28X coverage and 90% at 112X coverage, indicating the need to benchmark these parameters. After peak calling, the next step involves peak validation and assessment of control regions. Comparing peaks at candidate regions with control regions ensures robust detection of enhancer activity. For example, in our study, we compared the peaks at candidate enhancer regions to those within exons and noticed significantly reduced activity across the exonic regions **(Supplemental_Fig_S6),** in line with our assumption that coding regions of genes on an average display lower activity than candidate enhancer sites **(Supplemental Methods)**.

Another component of STARR-seq data analysis is to determine the sequence features of the regions designated as peaks. Traditionally, Motif Enrichment Analysis (MEA) tools such as HOMER (Heinz et al. 2010) or MEME (Bailey and Elkan 1994) have been used to detect enriched binding sites of known TF motifs within a set of active enhancer peaks. However, machine learning-based classification models can also serve a similar purpose. *For example*, Sahu and colleagues used logistic regression to predict enhancer activity based on the presence of TF binding motifs (Sahu et al. 2022). Furthermore, deep learning methods such as the Convolutional Neural Networks that predict enhancer activity by automatically learning the underlying sequence features can be powerful (de Almeida et al. 2022). Insights derived from these models add biological contexts to the identified peaks.

## Conclusion

Successful functional genomic studies take years to design and perform, and each step goes through repeated iterations in multiple replicates to optimize parameters for meaningful outcomes. Even steps involving established kits and protocols need to be scaled according to the study design and often require customizations and intermediate step validations. While each of these factors contribute to overall assay reproducibility, most published studies focus solely on the final outcome and tend to underexplain the methods, optimizations, and intermediate validations, resulting in large gaps of knowledge for researchers attempting to reproduce the results or tailor the study for their own biological questions. In fact, a comparison of human whole genome STARR-seq datasets across studies showed variations **(Supplemental_Fig_S7)**. Therefore, standardization of protocols and data reporting guidelines would benefit researchers and significantly improve the quality of the conducted assays. Here, we highlighted the different challenges in performing STARR-seq, a particularly long and difficult assay with huge potential to identify detailed enhancer landscapes and validate enhancer function. We emphasize the importance of reporting details related to biological context, underlying hypothesis, and experimental design, and protocol and bioinformatic analysis parameters to ensure replicability of each step and outcome of the assay.

### Box 1:Reproducibility across STARR-seq studies

To assess and quantify reproducibility for STARR-seq based studies, we systematically compared 24 studies (Arnold et al. 2013; Liu et al. 2017b; Barakat et al. 2018; Wang et al. 2018; Chaudhri et al. 2020; Liu et al. 2017a; Zhang et al. 2018; Kalita et al. 2018; Schöne et al. 2018; Johnson et al. 2018; Muerdter et al. 2018; Klein et al. 2020; Sahu et al. 2022; Peng et al. 2020; Vanhille et al. 2015; Lee et al. 2020; Verfaillie et al. 2016; Brandt et al. 2018; Klein et al. 2018; Van Ouwerkerk et al. 2020; van Weerd et al. 2020; Selvarajan et al. 2021; Glaser et al. 2021; Vockley et al. 2015) that performed STARR-seq experiments on mammalian or *Drosophila* cells (**Fig. 4**). We further identified 15 features critical for assay success and assessed their potential for reproducibility across the published studies. We scored each feature from 0 to 4 based on reporting of methodological details, design rationale, and QC measures, with 0 indicating no detail and 4 indicating reporting of complete detail for each feature that would allow for replication of the assay. The scoring was carried out independently by four individuals and aggregated. A detailed scoring rubric for each feature is provided in **Supplemental_Table_S4**. Our approach provides a method for quantifying the extent of reproducibility of published studies. In particular, we observed that authors typically mentioned steps such as read filtering, mapping tools, peak callers and QC experiments, but fail to provide associated intermediate data or parameters and cut-off thresholds used for their analyses, thereby reducing replicability of the steps. Furthermore, critical intermediate steps for sequencing such as library multiplexing and pooling that may be carried out at shared sequencing cores are often overlooked, leading to additional disparities in data quality. Therefore, standardization of these assay features and implementation of uniform data reporting guidelines will strongly improve reproducibility of STARR-seq assays.

**Figure 4:**
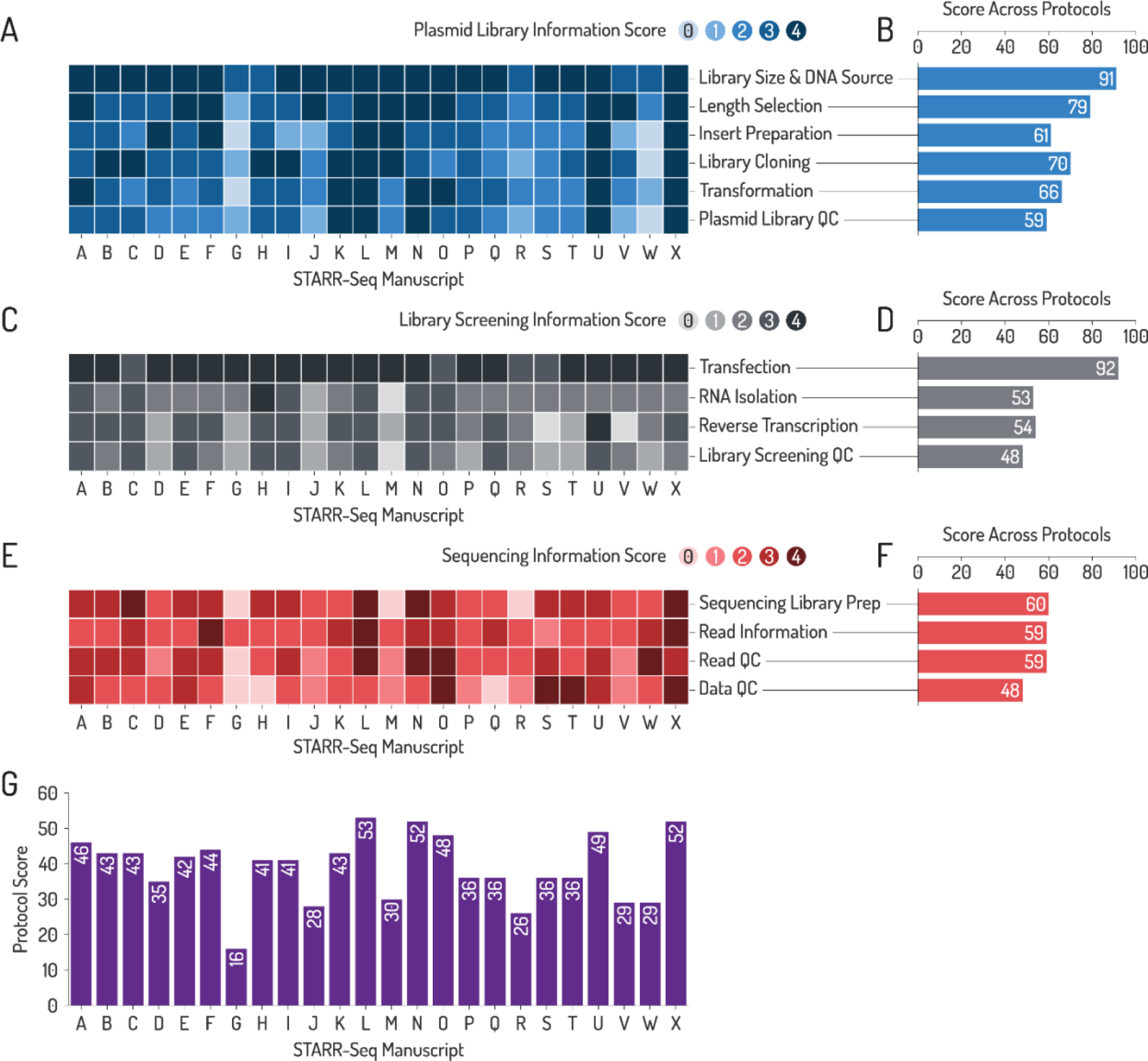
Reproducibility in current STARR seq studies. (A, C, E) Heatmap plot between feature scores for de-identified studies A-X illustrating scores for **(A)** assay design and plasmid library preparation, **(C)** library screening and **(E)** sequencing information. **(B, D, F)** Cumulative scores for each feature across studies illustrating reporting trends and highlighting features that are typically well explained or under-explained. **(G)** Cumulative scores for each assay (A-X) for all features enabling direct comparisons of reproducibility. We observed that critical design factors such as library scale and source and fragment length or experimental features such as transfection scored ≥3 across 87.5% - 100% of publications, with studies missing only minor details and explanations. In contrast, features such as library screening QC, data QC and data transparency scored ≥3 across 16.6% - 37.5% of publications.

### Box 2: Definitions of frequently used terminology in sequencing-based studies

1. **Assay reproducibility:** Measure of how accurately and consistently assay results can be re-obtained to re-affirm previous conclusions.
2. **Bead clean-up:** Magnetic DNA or RNA purification beads designed to be selective for DNA or RNA fragments. In DNA purification, the ratio of volume of beads to the volume of sample is selective of the length of DNA being purified. Lower bead volumes fail to bind longer DNA fragments, thereby filtering them out. This is a method for fragment length selection.
3. **Cloning arms or overhangs:** Sequences added to adapter-ligated (read1/read2) inserts that are complementary to cloning sites of the STARR-seq vector. Ligation is accomplished using LM_PCR to facilitate cloning of the fragments into the vector.
4. **De-duplication:** Bioinformatic process for removing PCR duplicates from sequenced reads.
5. **De-multiplexing:** Bioinformatic process by which all NGS reads from pooled libraries are separated into sample-specific reads, based on sample-specific i5/i7 barcodes.
6. **Episomal assay:** Assays that transfect one or more DNA sequences into a host cell whereby the transfected sequences do not integrate within host cell genome, and remain independent of the influence from the host cell genome.
7. **Experimental replicability:** Measure of how accurately an experiment can be repeated with the exact parameters to obtain the same output data.
8. **i5/i7 barcodes or indexes:** These are 6-8 bp sequences ligated to sequencing libraries to provide a unique identity for each sample when pooled with libraries from other samples. Libraries can use single indexing (either i5 or i7 index) or dual indexing (both i5 and i7).
9. **Input library:** STARR-seq sequencing library constructed either directly from plasmid library pool or from plasmid DNA extracted from transfected cells. The number of reads from the input is used to normalize the number of “output” reads to quantify enhancer activity.
10. **Library complexity:** The total number of unique fragments present in a library.
11. **Library delivery**: The process of delivering the prepared STARR-seq library into a host system through processes like transfection or transduction.
12. **Library dynamic range:** The ratio of read counts between the most active fragment in the output library to the least active fragment in the same library.
13. **LM_PCR: <u>L</u>**igation **<u>M</u>**ediated PCR is a type of PCR that uses primers carrying extended sequences (not having complementary sequence on the target strand) and ligates those extended sequences to the region being amplified.
14. **Massively parallel sequencing:** Next-generation sequencing (NGS) techniques that can sequence billions of reads in parallel based on a sequencing library.
15. **MPRA: <u>M</u>**assively **<u>P</u>**arallel **<u>R</u>**eporter **<u>A</u>**ssays are high-throughput assays that quantify the activity of a test fragment, typically cloned downstream of a minimal promoter, via massively parallel sequencing of barcoded reporter transcripts.
16. **Multiplexing:** A process where multiple libraries are pooled and sequenced on the same lane of the sequencer.
17. **Output library:** STARR-seq sequencing library built using self-transcribed reporter transcripts of candidate enhancers extracted from the host cell.
18. **P5/P7 Adapters:** Sequences flanking index (i5/i7) barcodes that help hybridize the fragment to the flow cell to enable sequencing.
19. **Peak calling:** Automated method by which sequenced STARR-seq reads from the output library are compared to reads from the input library to identify regions with significantly increased read counts or “peaks”.
20. **Read 1/Read 2 adapters:** Sequences immediately flanking the fragment of interest that are sequenced as “read 1” and “read 2” on a standard Illumina flow cell. These regions serve as recognition sequences for Illumina sequencing primers.
21. **Sequencing library:** Libraries containing fragments of amplicon DNA or cDNA to be sequenced using NGS. Each fragment consists of a core region of interest flanked by read 1 and read 2 adapter sequences, index barcodes, and P5 and P7 flow cell adapters.
22. **STARR-seq plasmid library:** Library comprising of candidate enhancer fragments cloned downstream of a minimal promoter in a STARR-seq vector.
23. **STARR-seq: <u>S</u>**elf **<u>T</u>**ranscribing **<u>A</u>**ctive **<u>R</u>**egulatory **<u>R</u>**egion **<u>Seq</u>**uencing, is a specialized episomal MPRA that is used to directly measure enhancer activity by comparing output transcripts produced by a candidate enhancer sequence cloned downstream of a minimal promoter to the number of copies of the enhancer fragments used as input prior to transcription.
24. **UMIs: <u>U</u>**nique **<u>M</u>**olecular **<u>I</u>**dentifiers are randomly synthesized oligos (unknown sequences) of fixed length that can be added to fragments. If fragments undergo amplification via PCR, UMIs added to those fragments prior to PCR are also duplicated during PCR. This enables detection of PCR duplicates, as opposed to identical mRNA self-transcripts.

## Declarations

### Consent for publication

All authors agree and consent for publication of the manuscript.

### Competing interests

The authors declare that no competing interests exist in relation to this work.

## Acknowledgements

We acknowledge Drs. Matthew Jensen, Yasuhiro Kyono, Istvan Albert, Aswathy Sebastian, Howard Salis, Ross Hardison, Corrine Smolen, and the Penn State Genomics Core Facility for technical support for this project. This work was supported by the NIH grants R01-GM121907, R21-NS122398, and resources from the Huck Institutes of the Life Sciences to SG.

## Supplemental Materials

### Supplemental Text

#### STARR-seq experiment

We built a STARR-seq plasmid library spanning approximately 33 Mbp of the human genome and performed multiple STARR-seq runs on HEK293T cells. Our target regions were shortlisted from existing ChIP-seq data on HEK293 or HEK293T cells available on ENCODE (Birney et al. 2007). In brief, we first overlapped ChIP-seq sites binding to the histone modifications H3K27Ac (for active enhancers) and H3K4Me1 (for active or poised enhancers). Next, we intersected these regions with all TF-ChIP-seq sites on HEK293 (from ENCODE) to obtain a comprehensive enhancer catalog spanning 46,010 ChIP-seq sites. The selected regions comprise TF-binding sites that overlap with recognized enhancer marks. We captured our target library from commercially available human whole genome DNA using hybridization and capture probes and cloned the captured fragments into the human STARR-seq vector to build a STARR-seq plasmid library **(see Supplemental_Fig_S1, STARR-seq Protocol)**. To assess the impact of different genomic mutations on enhancer activity, we transfected the library into seven different mutant HEK293T lines and one wild-type line in three biological replicates and isolated reporter specific mRNA to build 24 STARR-seq output screening libraries. We directly amplified the STARR-seq plasmid library in three replicates for the input. We sequenced 24 output screening libraries and 3 input libraries using a NextSeq2000 sequencer with approximately 45 million reads per sample.

While conducting the assays we came across various design and protocol inconsistencies that enabled us to generate a list of STARR-seq design considerations, best practice guidelines and quality control checkpoints **(see Supplemental Tables)**. Our analysis pipeline is also posted on GitHub (links provided in **Supplemental Methods**). We used our data to demonstrate the effects of read filtering, significance of read depth, and compared different peak callers. All sequence data generated in this study have been submitted to the NCBI BioProject database (https://www.ncbi.nlm.nih.gov/bioproject/) under **accession number PRJNA879724**. We also reanalyzed datasets from existing STARR-seq studies and assessed them for their data quality control steps and compared them to our data (see **Supplemental methods**). We also evaluated 24 STARR-seq studies and scored important assay features for each study based on the number of details reported by the authors **(see Supplemental_Table_S4)**.

### Supplemental Methods

#### Read deduplication, alignment, and filtering

We used custom scripts to remove all PCR duplicates from raw FASTQ files of STARR-seq input and output libraries obtained after demultiplexing. We then aligned the paired-end reads to GRCh38 human genome assembly using BWA-MEM (Li 2013) with default settings. Next, we removed reads that were *(i)* unaligned, *(ii)* low quality (mapping quality score<30), *(iii)* multi-mapped (reads that mapped to multiple locations with equal confidence), and *(iv)* off-target using SAMtools (Li et al. 2009) with the following parameters: -F 2828 -f 2 -q 30.

To compare read loss across published datasets with our data, we reanalyzed ATAC-STARR-Seq data from Wang and colleagues and human whole-genome STARR-seq data from Johnson and colleagues using our computational pipeline with modifications to the deduplication strategies (Johnson et al. 2018; Wang et al. 2018). We first aligned paired-end reads using BWA-MEM. Next, we filtered PCR duplicates using Picard (Broad Institute. 2019). Finally, we removed (i) unaligned, (ii) low quality (mapping quality score<30), (iii) multi-mapped (reads that mapped to multiple locations with equal confidence) reads and compared reads across the three datasets **(Supplemental_Fig_S2)**. Analysis pipelines are posted on GitHub: https://github.com/deeprob/starrseq_dedup_align_filter.

#### Data quality control

We assessed data quality and replicability by calculating correlation of filtered read counts between input and output library replicates. Additionally, we calculated correlation of Reads per Million (RPM) normalized output-over-input fold changes between replicates **(Supplemental_Fig_S3)**. The output sequencing libraries were generated from the wild-type control HEK293T line. To compare quality of published datasets with our data, we assessed the correlation of filtered read counts between input and output replicates for our STARR-seq libraries and the reanalyzed datasets (Johnson et al. 2018; Wang et al. 2018) **(Supplemental_Fig_S4)**. We also performed Principal Component Analysis (PCA) and visualized the first two principal components of input and output replicate filtered read counts for all reanalyzed datasets including our own **(Supplemental_Fig_S5)**.

#### Peak calling

We called peaks using previously published tools, MACS2 (Zhang et al. 2008), STARRPeaker (Lee et al. 2020), CRADLE (Kim et al. 2021) and DESeq2 (Love et al. 2014) using default settings. We merged the input and output replicates before peak calling using MACS2 and STARRPeaker. For CRADLE and DESeq2, the replicates were kept separate since they both utilize the variance between replicates to adjust p-value estimates of peaks. Before peak calling with DESeq2, we fragmented all regions using a sliding window of size 500 bp and a stride of 50 bp. Next, we calculated the input and output library coverage for each of these windows using BEDTools (Quinlan and Hall 2010). The input and output replicate-wise library coverage for each window was used by DESeq2 to identify differentially active regions. We used BEDTools to intersect CRADLE- and DESeq2-called “active” peaks with peaks called by other callers.

For comparing the number of peaks by varying library coverage, we randomly subsampled the input and output control libraries from 10% to 90% with increments of 5% using SAMtools (Li et al. 2009). For each subsample, we also created three replicates by changing random seed parameter. We called peaks in the subsampled libraries using STARRPeaker with the default settings. Our peak calling pipeline is posted on GitHub: https://github.com/deeprob/starrseq_peak_call.

#### Activity comparison between peaks and exonic regions

To compare activity between peaks and exonic regions, we first identified regions in our library which overlapped with known exonic regions from the reference human genome (GRCh38).

Next, we calculated the RPKM normalized coverage of filtered reads fold changes between output and input libraries for both STARRPeaker called peaks and the identified exonic regions. Finally, we compared the fold change distributions between peaks and exonic regions using t-test **(Supplemental_Fig_S6)**.

#### Assessing reproducibility of published STARR-seq datasets

To compare variation in enhancer activity for a given fragment between different STARR-seq studies, we used processed bigwig files from Johnson and colleagues and Lee and colleagues for their respective whole genome studies (Johnson et al. 2018; Lee et al. 2020). We used output over input fold change signals from the two whole genome libraries reported by Lee and colleagues, submitted as bigwig files. Johnson and colleagues reported separate bigwig files for input and output read signals for their whole genome library. In this regard, we first normalized the input signal by Z-score normalization method. Next, we calculated fold change of output over normalized input. Finally, we measured Pearson and Spearman correlation of fold changes between the three libraries **(Supplemental_Fig_S7)**.

## Supplemental Figures

**Supplemental_Fig_S1:**
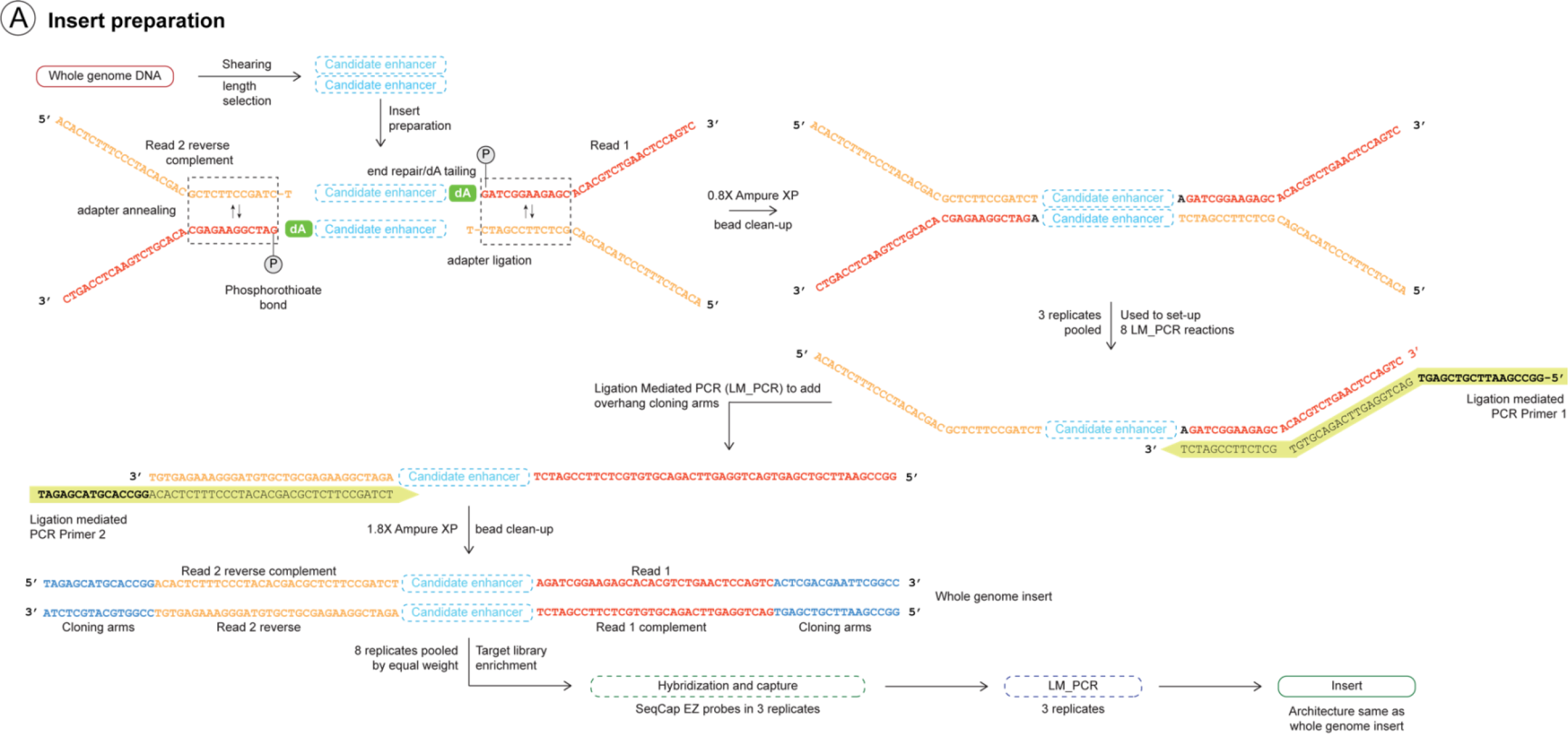
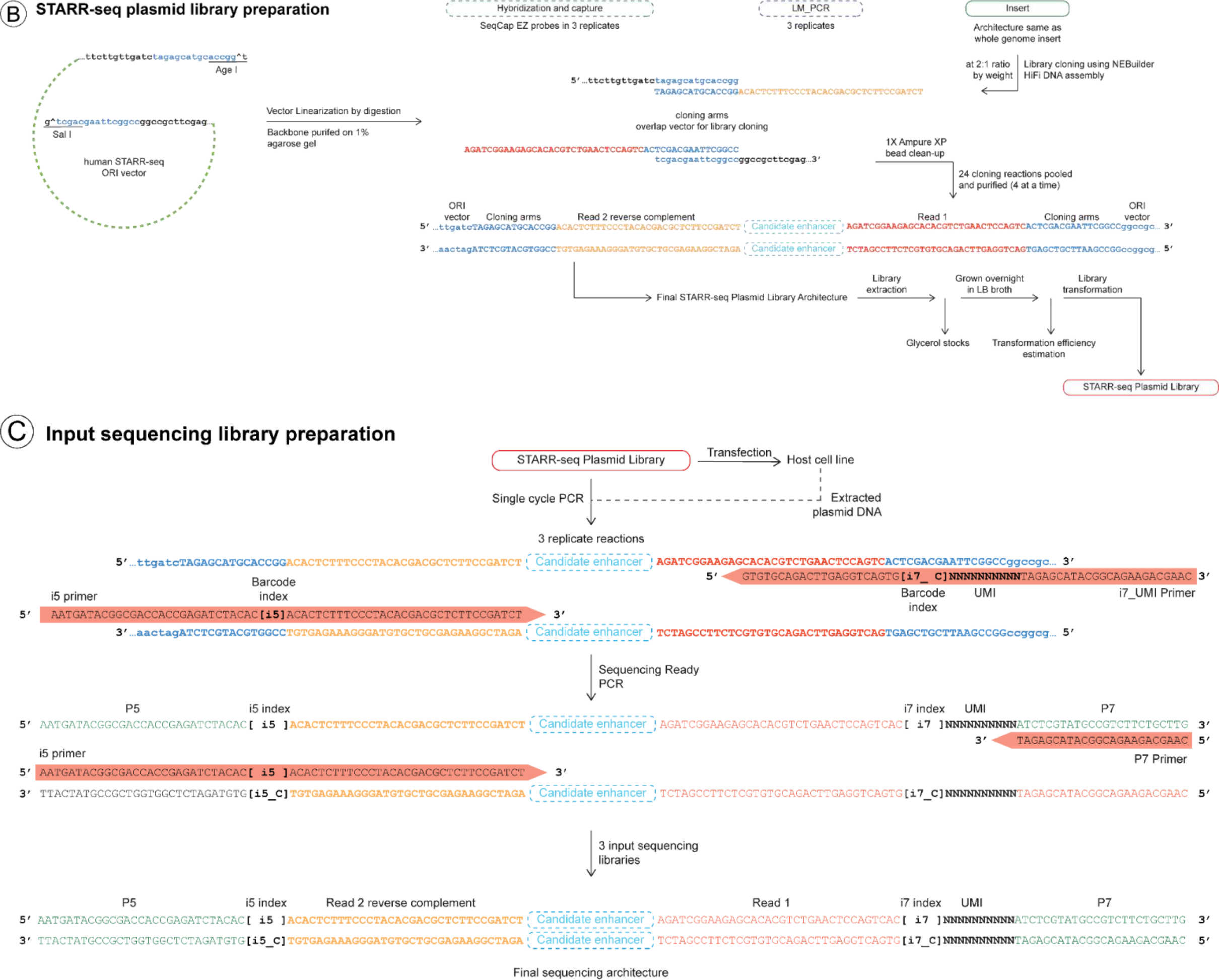
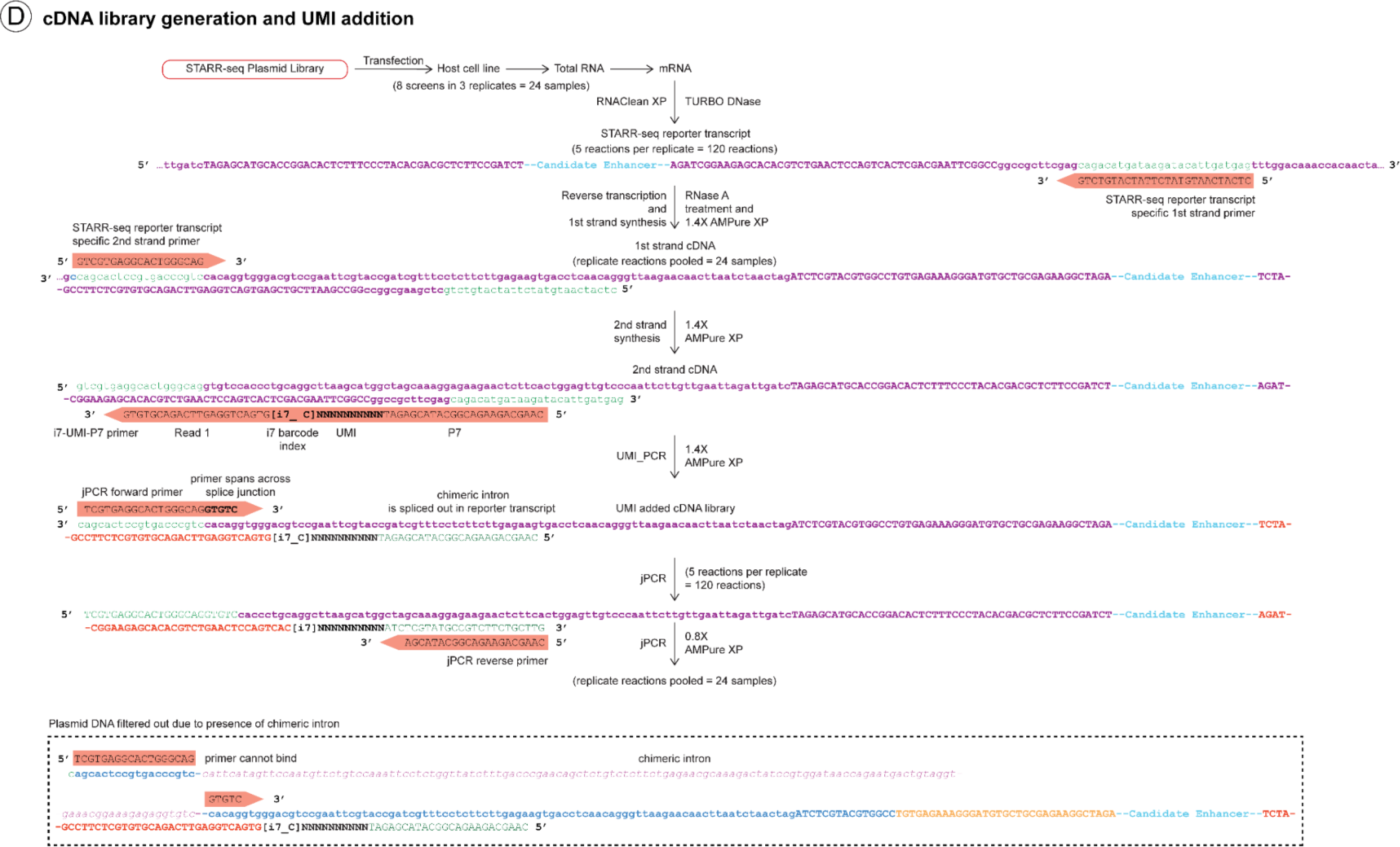
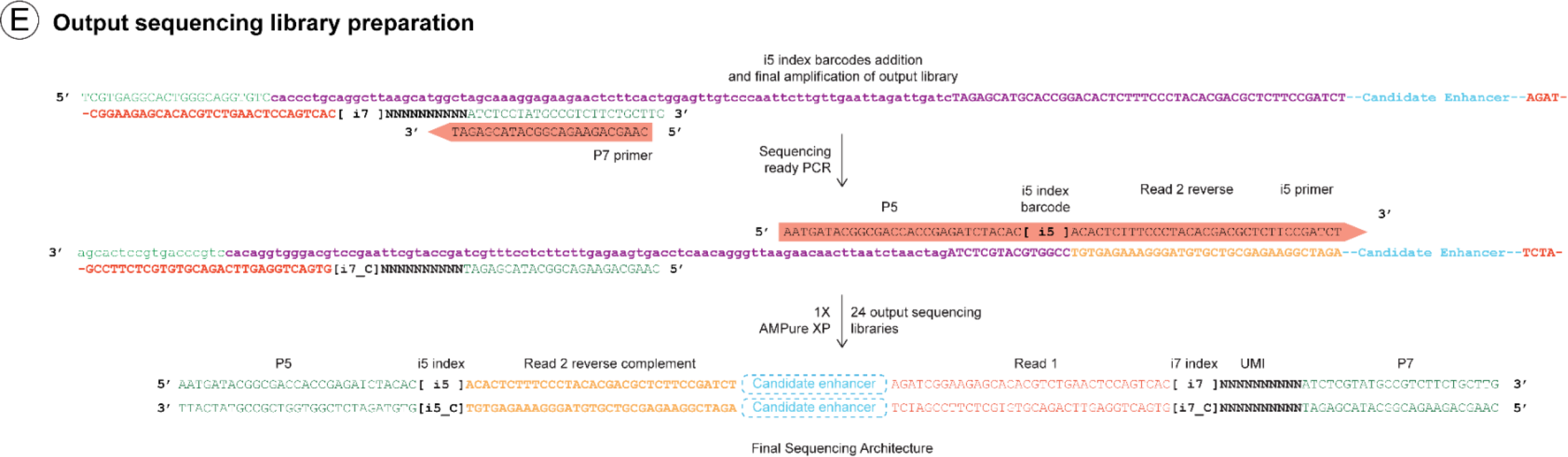
A schematic of STARR-seq experimental protocol and complete library sequence information are shown. **(A)** Insert preparation including addition of sequencing adapters and cloning arms to library fragments to facilitate sequencing and library cloning. **(B)** STARR-seq plasmid library preparation including vector linearization and cloning of library inserts into human STARR-seq vector (Muerdter et al. 2018) followed by library amplification through transformation. **(C)** Preparation of ‘input’ sequencing library either directly from plasmid library or from DNA extracted from library-transfected host cells. This step adds on UMIs and index barcodes to the library prior to sequencing to sort for PCR duplicates and to sequence multiple libraries on the same sequencing lane (multiplexing). **(D)** cDNA library generation for STARR-seq screening includes transfection of the plasmid library into a host cell and reverse transcription of self-transcribed reporter transcripts. This step also involves adding UMIs for detecting and removing PCR duplicates. **(E)** Preparation of ‘output’ sequencing library involves adding sequencing barcode indexes to the screening library before sequencing to enable multiplexing. Both ‘input’ and ‘output’ libraries are pooled and sequenced in parallel for enhancer screening.

**Supplemental_Fig_S2:**
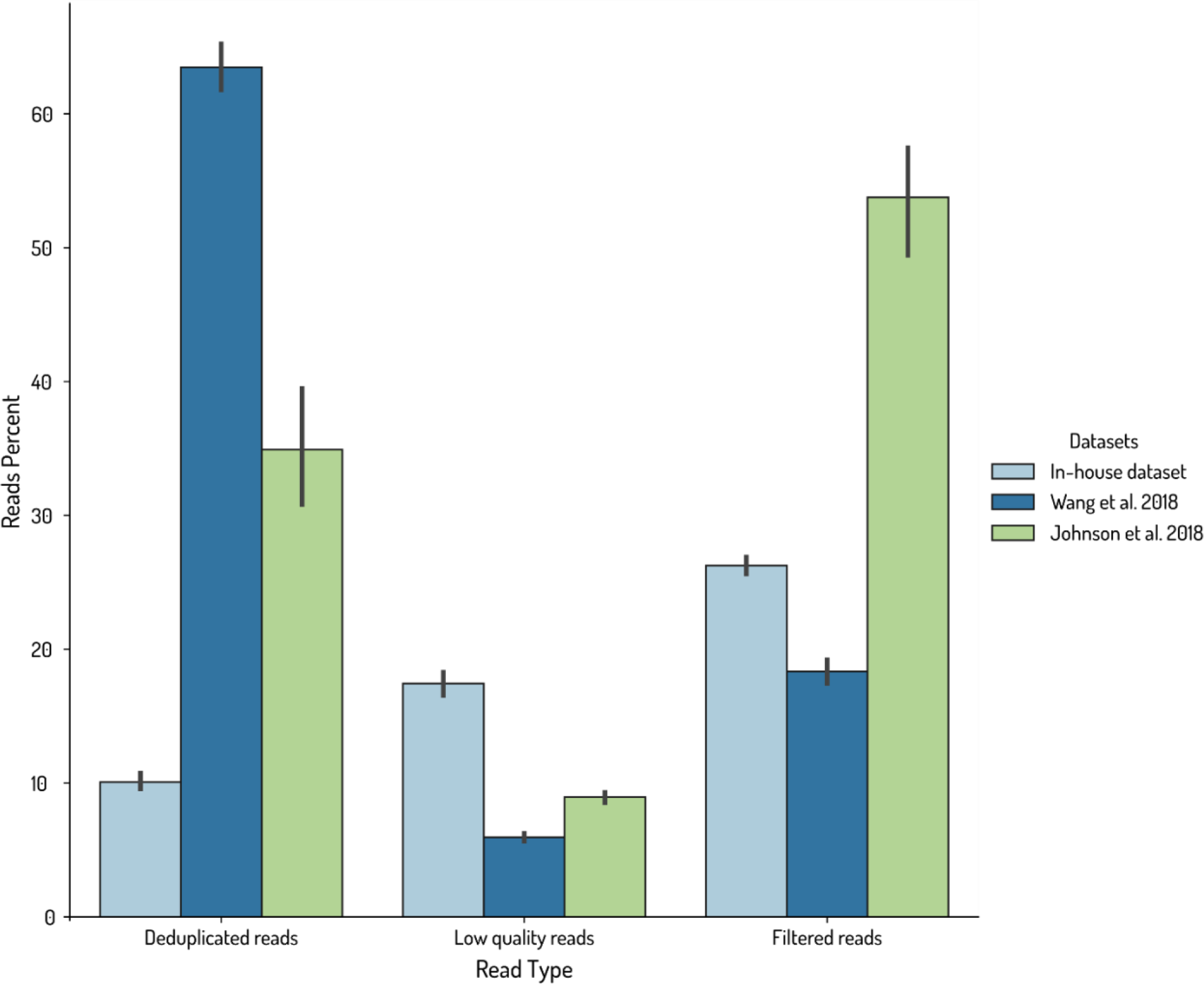
Bar plots illustrating read loss observed across published STARR-seq datasets compared to our in-house STARR-seq assay are shown. Study by (Wang et al. 2018) and in-house STARR-seq includes focused assays, while (Johnson et al. 2018) includes data from a whole genome STARR-seq assay.

**Supplemental_Fig_S3:**
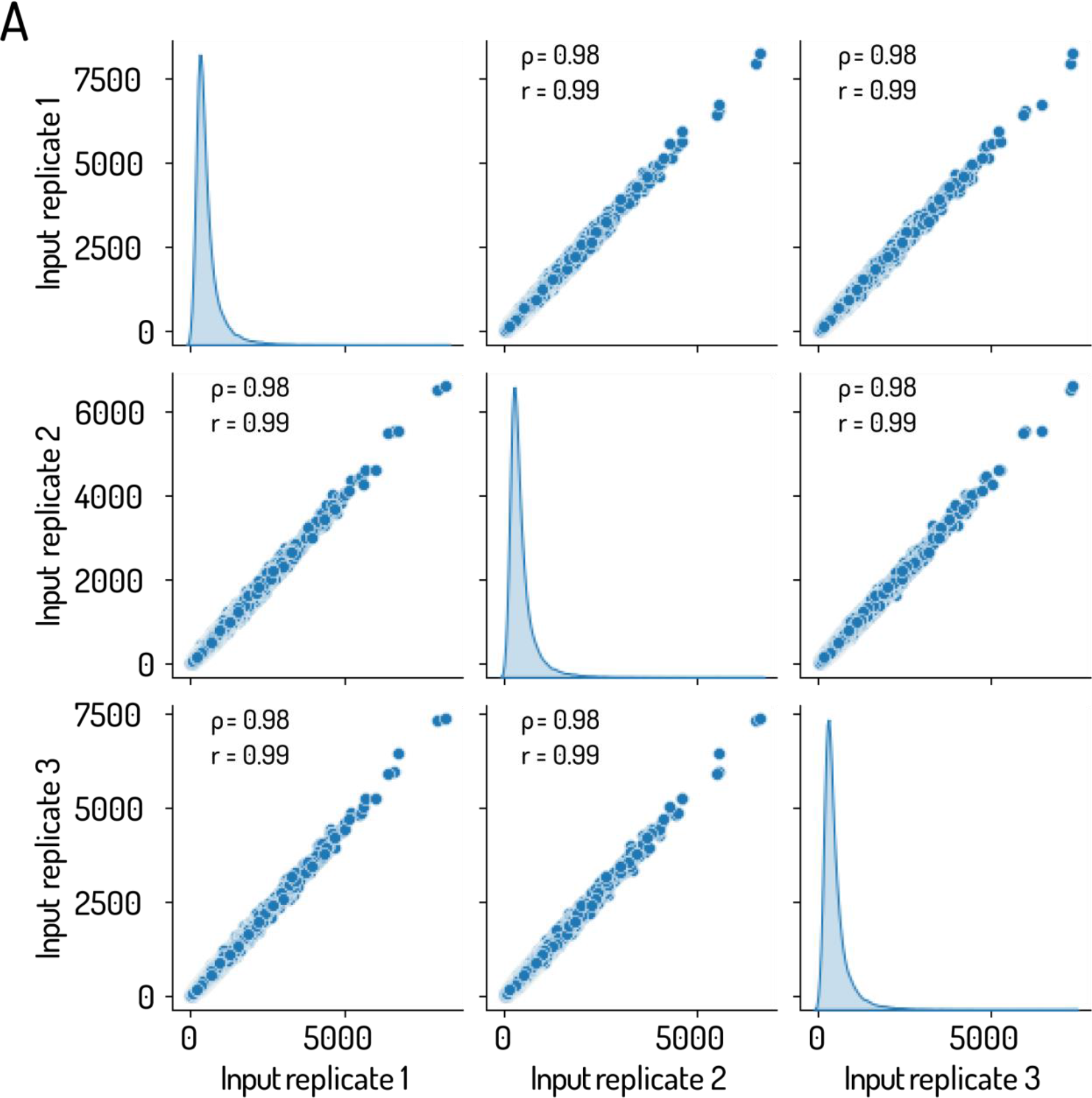
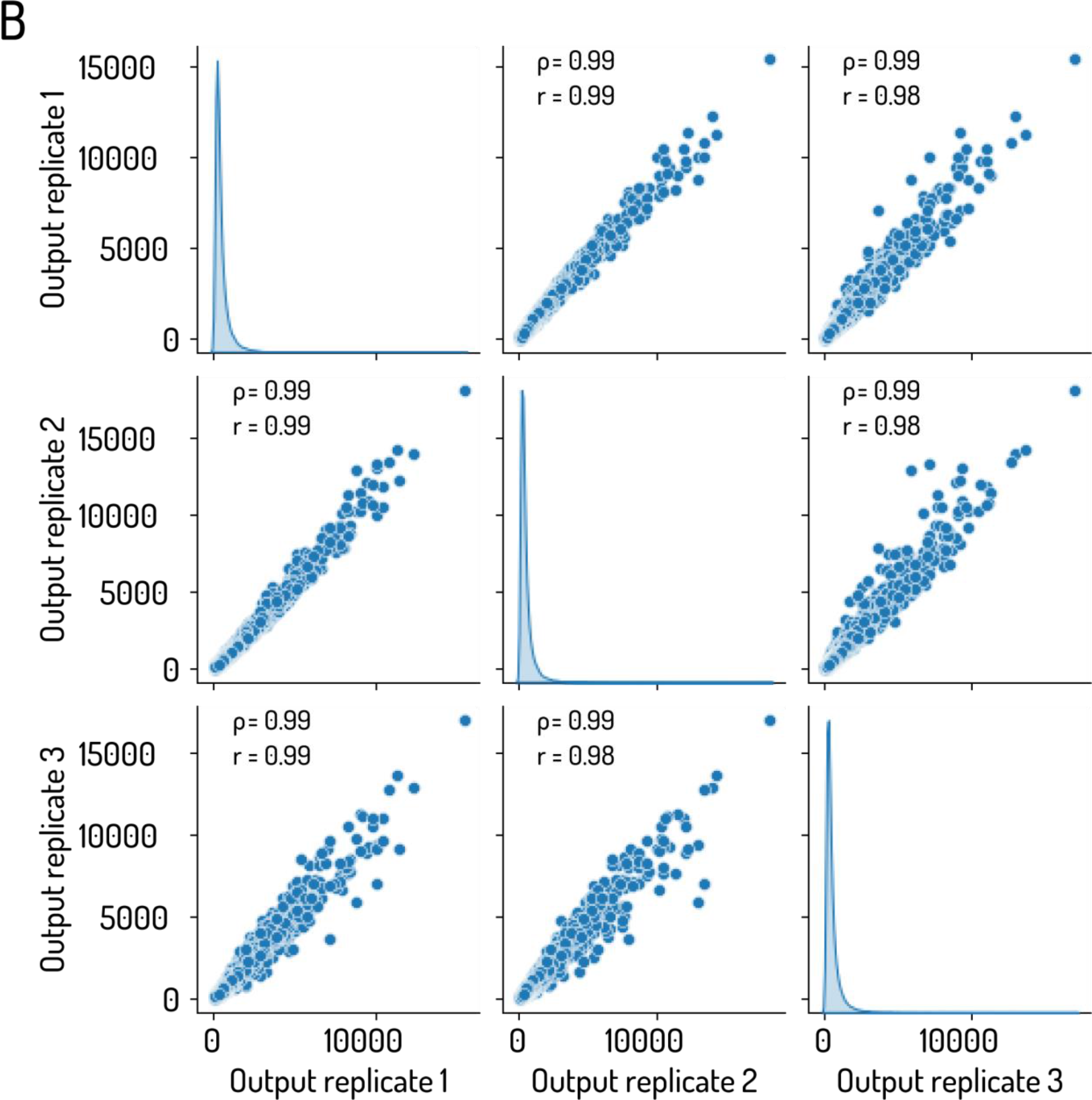
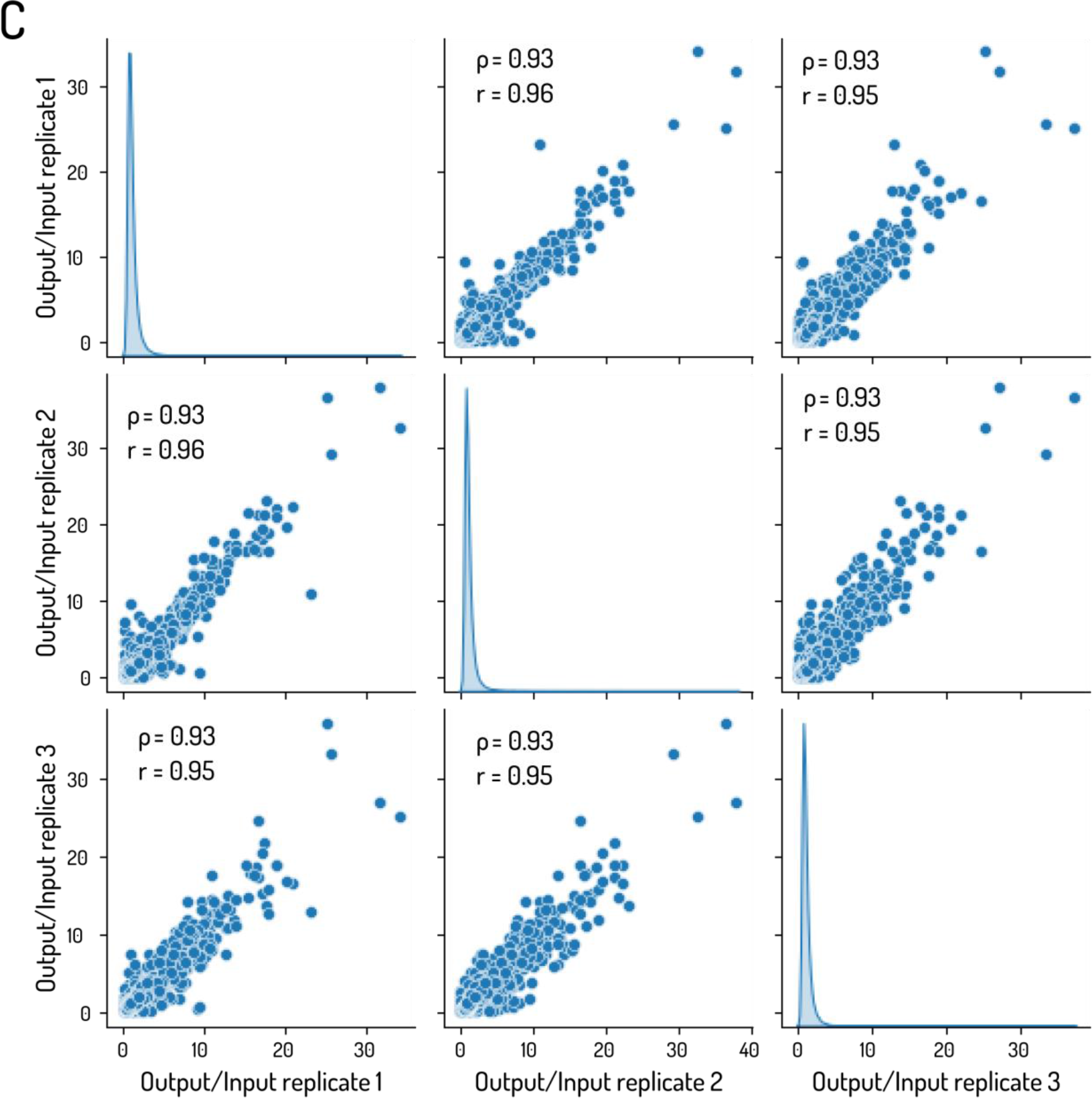
**Scatter plots and correlation (Spearman and Pearson co-efficient values indicated) across in-house STARR-seq libraries**. **(A)** Correlation observed across three replicates of input sequencing libraries prepared from direct amplification of in-house STARR-seq plasmid library is shown. **(B)** Correlation observed across three replicates of output sequencing libraries is shown. The output library consists of the STARR-seq experiment conducted on control HEK293T cell line. **(C)** Correlation of output over input fold changes across input and output libraries is shown. Input replicates assigned randomly to output replicates 1, 2 and 3

**Supplemental_Fig_S4:**
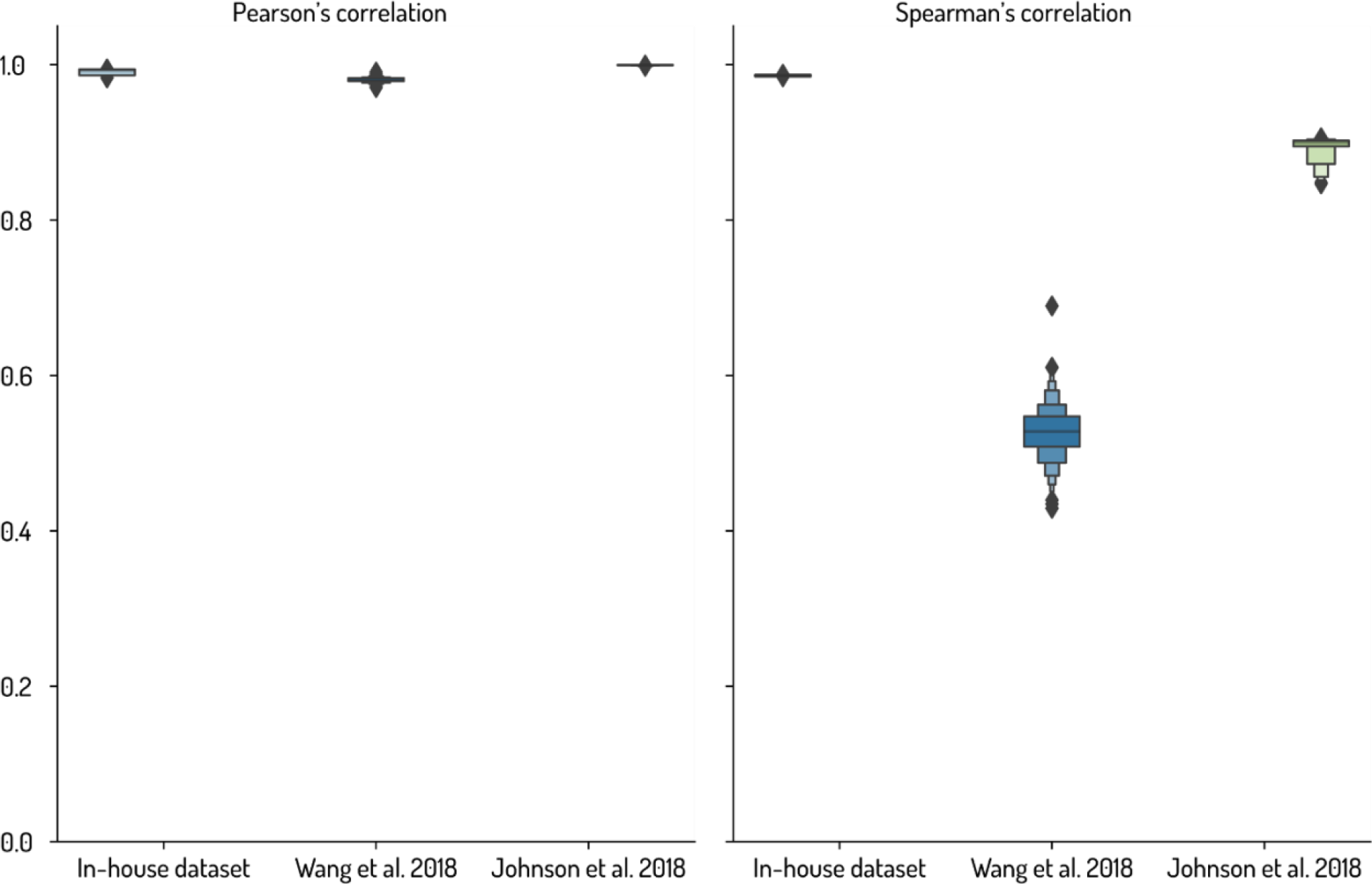
Comparison of Pearson’s and Spearman’s correlation of input and library read counts across published studies (Wang et al. 2018; Johnson et al. 2018) and in-house STARR-seq assay is shown. Study by Wang and colleagues and in-house dataset are focused assays while Johnson and colleagues report a whole genome STARR-seq assay.

**Supplemental_Fig_S5:**
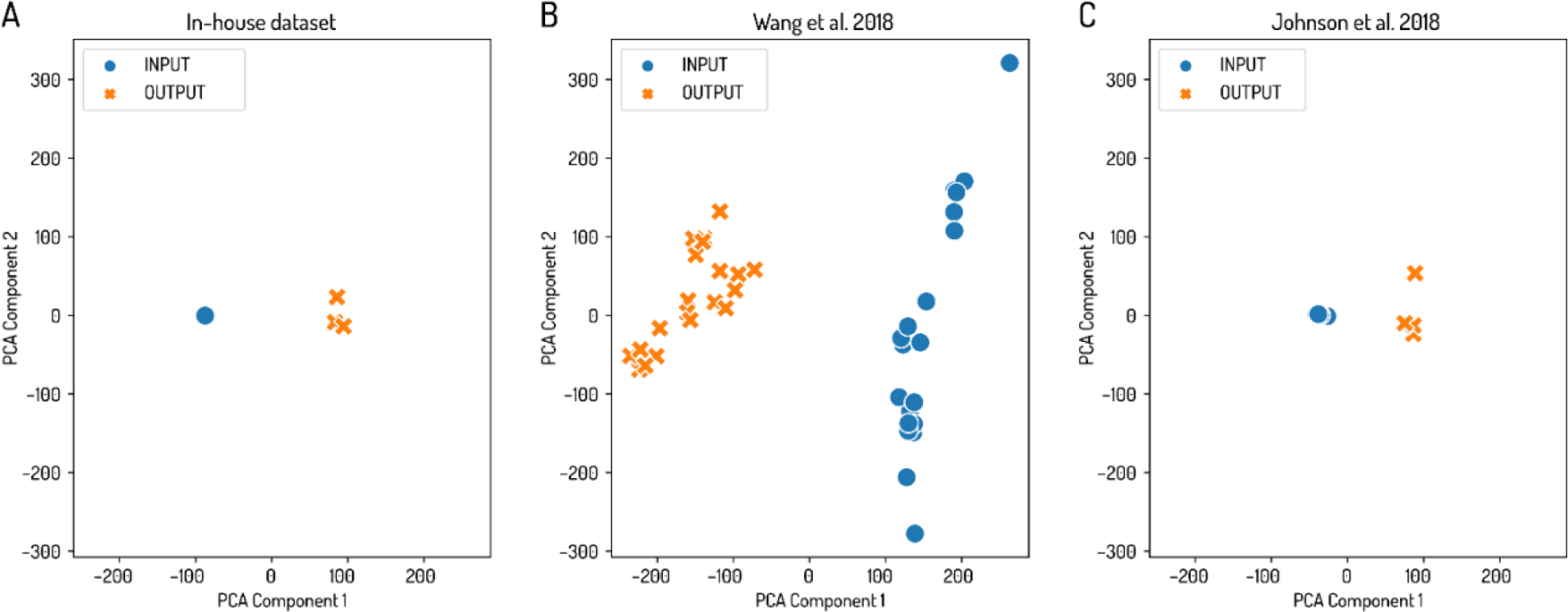
PCA plots demonstrating successful clustering of input and output STARR-seq library replicates for **(A)** Our in-house STARR-seq input and control library **(B)** Dataset from (Wang et al. 2018) **(C)** Dataset from (Johnson et al. 2018). Please note in **(A)** all three input library replicates (blue dots) cluster together.

**Supplemental_Fig_S6:**
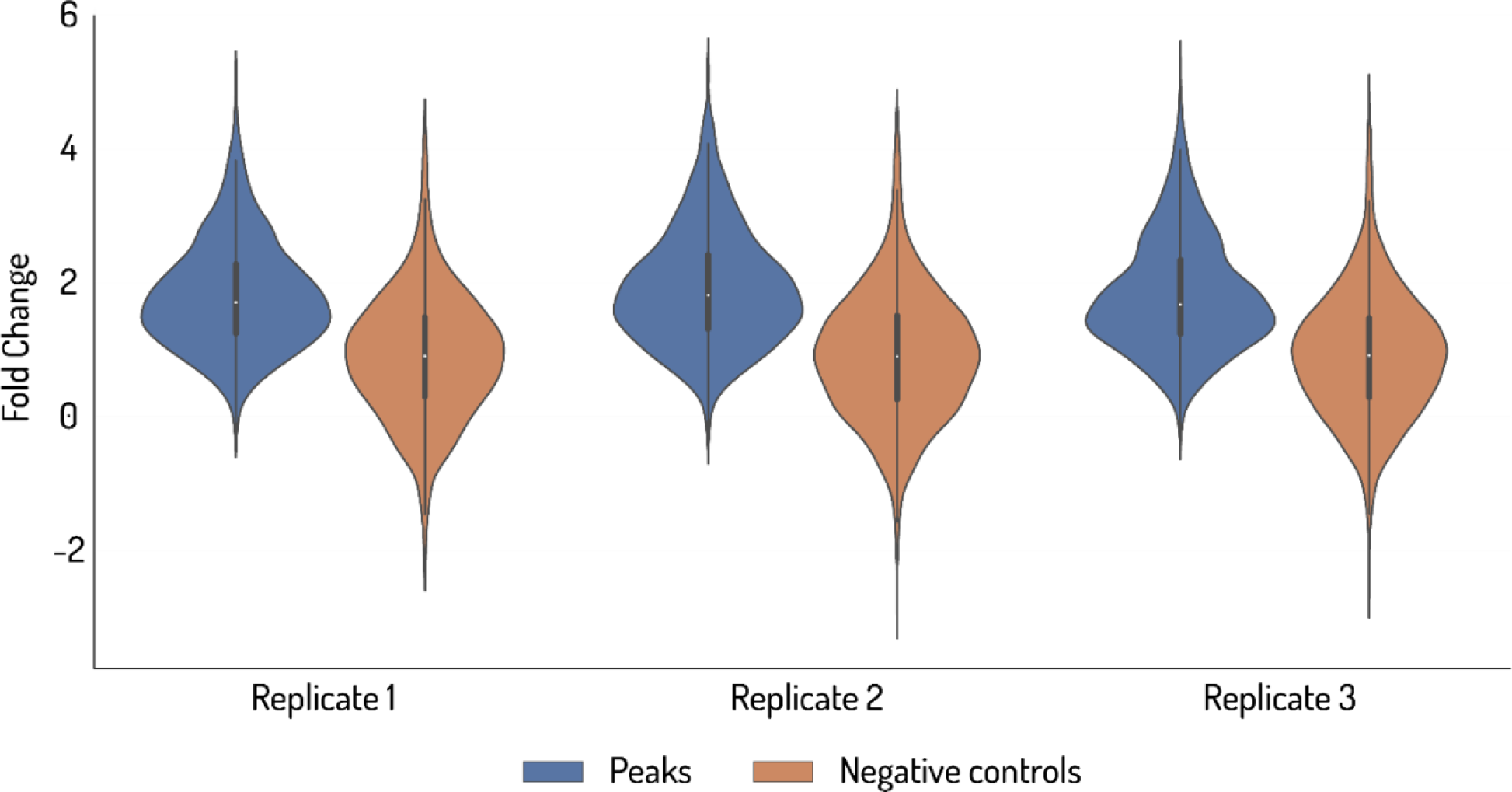
Violin plots comparing the log2 fold change of normalized reads (Output over Input Reads Per Kilobase Million) observed between the STARRPeaker-called peaks and exonic regions from our in-house STARR-seq dataset as negative controls are shown. STARR-seq activity of exons was significantly lower than that of the peaks from enhancer regions (t-test statistic: 53.14; p-value: 0.0).

**Supplemental_Fig_S7:**
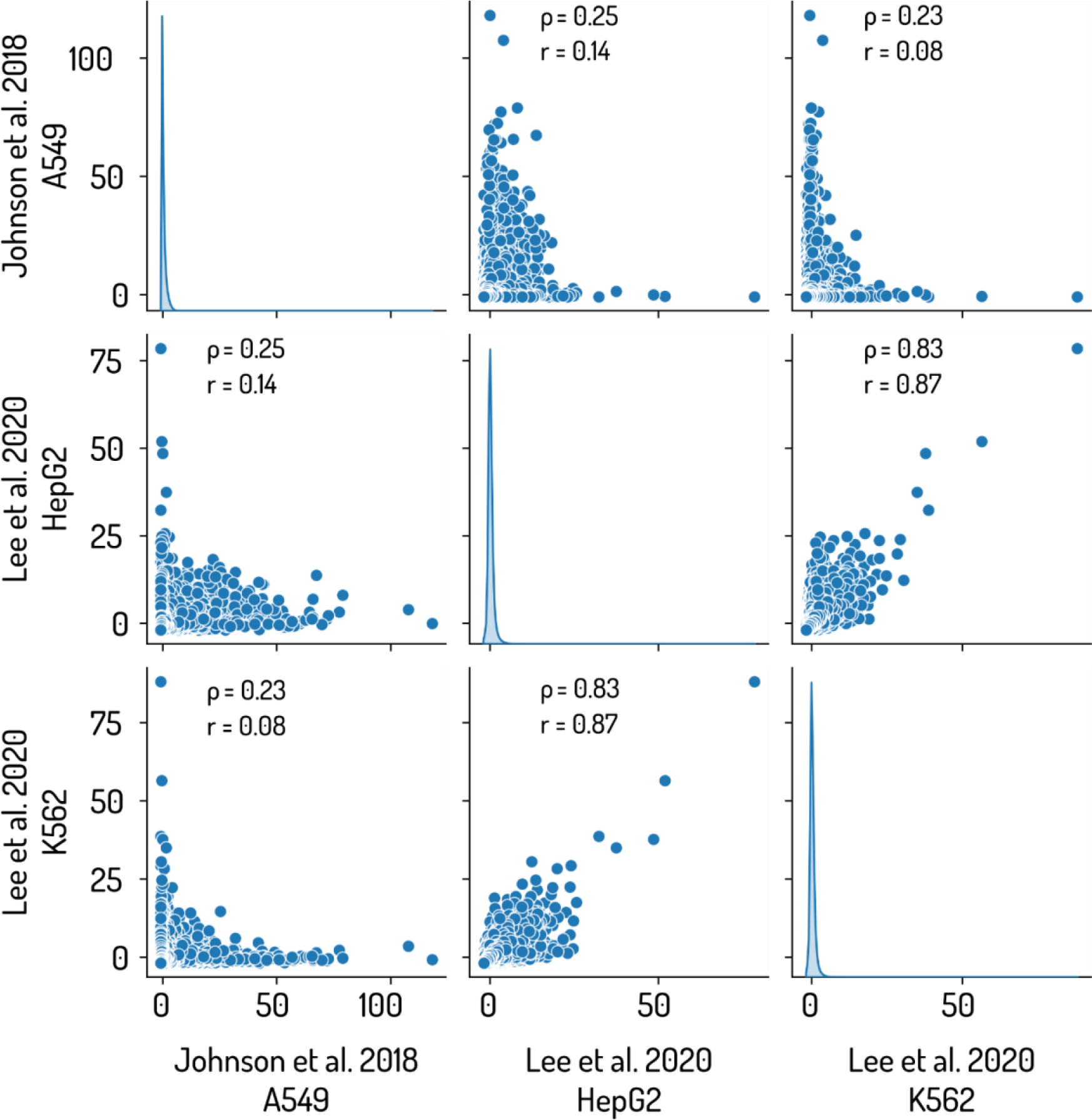
Spearman and Pearson’s correlations for output over input fold changes for three whole genome STARR-seq assays are shown. Two assays were conducted for the same study (Lee et al. 2020) in K562 and HepG2 lines using the modified human STARR-seq vector containing the ORI promoter (Muerdter et al 2018). The third assay was conducted for a different study (Johnson et al. 2018) on A549 cells using the original STARR-seq vector (Arnold et al 2013).

## Supplementary Tables

**Supplemental_Table_S1:** Table shows suggested rationales for core experimental design factors including library size, fragment length, and DNA source for different biological questions, and for different versions of STARR-seq or MPRAs.

|  | Biologically question | Suggested methods | Library size | Fragment length | Source of target DNA for library preparation | References |
| --- | --- | --- | --- | --- | --- | --- |
| 1 | Genome-wide assessment for enhancer potential | Whole genome STARR-seq | Library size is determined by the target species (such as *Drosophila* or human) being assayed. | Longer fragment lengths (>500 bp) will enable easier coverage of target genome.<br>Shorter fragments (300 bp-500 bp) will require more effort and resources to obtain comparable library complexity and coverage.<br>Sensitivity of enhancer activity detected may vary based on the length of the fragment. | Genomic DNA<br>Oligo synthesis not recommended due to limitations of shorter synthesized fragment length. | (Arnold et al. 2013; Liu et al. 2017b; Zabidi et al. 2015; Peng et al. 2020; Sahu et al. 2022) |
| 2 | Assess the effect of changes in a cellular environment such as drugs, hormone treatment, or genomic mutations on enhancer activity | Whole genome STARR-seq | Optimal for assaying global changes in enhancer activity due to host cell perturbations.<br>Harder to achieve high library depth.<br>Higher costs and resources. | Similar guidelines as whole genome libraries mentioned above.<br>Very long fragments (>1000 bp) may not capture smaller changes in activity. | Genomic DNA | (Johnson et al. 2018; Shlyueva et al. 2014) |
|  |  | Focused STARR-seq<br>CHEQ-seq | Useful for assaying the effect on candidate enhancers.<br>Simpler and cost-efficient library preparation and sequencing.<br>Allow for higher sequencing depths that can reveal subtle changes. | Longer lengths can help provide sequence contexts for the observed activity changes.<br>Shorter fragments can help pinpoint active regions responsive to perturbation. | Target DNA can be obtained from different methods such as synthesis (<300 bp) or capture. | (Verfaillie et al. 2016; Schöne et al. 2018) |
| 3 | Validation of enhancer function and assessment of activity in open chromatin regions | • ATAC-STARR-seq<br>• FAIRE-STARR-seq<br>• ChIP-STARR-seq | • Determined by the number of regions captured through ATAC/FAIRE/ChIP capture methods from the target genome.<br>• For ChIP-STARR-seq, target size also depends on the number of TFs or histone modifications assessed. | • Depends on the length of the open chromatin fragments captured.<br>• Lengths of fragments can also vary within the same library but library preparation strategy should be adjusted to account for potential bias caused by variable fragment lengths.<br>• Captured fragments may be sheared and size selected, if required, prior to cloning. | • Genomic DNA | (Wang et al. 2018; Chaudhri et al. 2020; Glaser et al. 2021; Hansen and Hodges 2022; Barakat et al. 2018) |
| 4 | Assessment of non-coding mutations in populations or effect of SNPs and enhancer-flanking regions on enhancer activity | • CapSTARR-seq | • Determined by number of mutations being assessed. CapSTARR-seq is useful for capturing larger libraries from genomic DNA isolated from populations carrying the mutations being tested. | • Fragment lengths can vary with type of mutation assessed and capture method.<br>• For assessment of impact of SNPs, shorter fragment lengths are recommended to remove additional sequence contexts. | • Genomic DNA | (Liu et al. 2017a; Vanhille et al. 2015) |
|  |  | • BiT-STARR-seq<br>• SynSTARR-seq | • Determined by the number of tiled fragments synthesized to cover target regions.<br>• Uniformly tiled fragments designed for target region allow for uniform library representation.<br>• Size limited by synthesis costs and capabilities. | • Fragment length limited to 300 bp.<br>• Fragments can be tailored to contain SNPs, mutations, or even multiple bp configurations of a particular mutation. | • Synthesized oligos. | (Kalita et al. 2018; Schöne et al. 2018) |
|  |  | • LentiMPRA | • Similar to syn-STARR/BiT-STARR-seq<br>• Size additionally limited due to requirement of multiple barcodes per designed fragment and transduction efficiency. | • Length limited by synthesis limitations.<br>• Fragments should also be designed to contain multiple barcodes.<br>• Fragments are integrated into the genome and provides additional context. | • Synthesized oligos. | (Inoue et al. 2017) |
| 5 | Assessment of enhancer activity in tissue systems using explant cultures or in organs in an *ex-vivo* setting. | *In-vivo* or *Ex-vivo* STARR-seq | Library size limited by transduction ability and efficiency, and requires validation of delivery method and library loss. | Longer fragments recommended for testing broader sequence context.<br>Shorter fragments recommended for assessing mutations on the fragments or treatments on the host. | Determined by fragment length | (Lambert et al. 2021) |
| 6 | Assessment of enhancer activity in *Drosophila* cell lines | Fly STARR-seq (Whole genome or focused) | Range from whole genome to focused library based on the question. | Fragment length is flexible | Genomic DNA | (Arnold et al. 2013; Zabidi et al. 2015; Shlyueva et al. 2014) |
| 7 | Assessment of enhancer - promoter associations | ExP-STARR-seq | Library consists of synthesized fragment combinations and therefore library size limited by synthesis costs. Eg: Bergman and colleagues used 1000 enhancer and 1000 promoter combinations. | Fragment length of enhancer, promoter, and spacer element determined during design. The design should also consider barcodes associated with enhancer-promoter pairs. | Synthesized oligos. | (Bergman et al. 2022) |
| 8 | Assessment of enhancer activity integrated into the genome. | LentiMPRA | As mentioned in 4. | Fragment length limited by synthesis limitations.<br>Longer fragment use yet to be reported. | Synthesized oligos. | (Inoue et al. 2017; Klein et al. 2020) |
| 9 | Assessments of multiple base-pair configurations of promoters or enhancers | Classical MPRA | Limited by synthesis constraints and the need to add unique barcodes to each bp configuration. | Fragment length limited by synthesis limitations.<br>Longer fragment use yet to be reported. | Synthesized oligos. | (Patwardhan et al. 2012; Melnikov et al. 2012) |

**Supplemental_Table_S2:** Proposed guidelines for assay scaling based on the library size, fragment length, and estimated read depth are shown. For read depth, if the dynamic range of the library (the ratio of read count between the most active fragment in the output library to the least active fragment) can be estimated, then the total number of reads in both input and output libraries is given by the product of the dynamic range, the minimum number of reads per fragment for it to be considered active, and the total number of fragments in the complete library. However, to uncover the true dynamic range in each output library, input and output libraries should be sequenced using the same number of reads. Additionally, after library construction and validation, studies should report complete library details including the length of the target region, number of unique fragments obtained, and the final sequence depth or fold coverage for each unique fragment after sequencing to enable assay reproducibility.

|  | Library size | Fragment length range | Minimum number of fragments to cover target region | Number of transformation reactions | Reads for sufficient depth |
| --- | --- | --- | --- | --- | --- |
| | Length of target region (in base pairs) | Determined by library size and biological question. | (Library size/frag length)<br>Eg: Assuming upper limit of library size and lower limit of fragment length. | $t = (n * c)/e$<br>t = number of transformations<br>n = number of minimum fragments<br>c = number of copies per fragment<br>e = transformation efficiency (i.e., transformants per reaction) | $r = n * x * z$<br>r = total number of reads<br>x = minimum number of reads required per variant<br>z = presumptive library dynamic range |
| 1 | Up to 100 kb | 200-1200 bp | 100 kb/200 bp = 500 fragments minimum. | $t = (500 * 1000)/10^6 = 0.5$<br>Hence a single transformation reaction would yield sufficient colonies | For both libraries assuming z = 100, x = 10.<br>$r = 500 * 10 * 100 = 500,000$ reads for output library |
| 2 | 100 kb-1 Mb | 200-1200 bp | 5000 fragments minimum | $t = (5000 * 1000)/10^6 = 5$ reactions | $r = 5000 * 10 * 100 = 5,000,000$ for output library |
| 3 | 1-100 Mb | 300-1200 bp | 333,333 fragments minimum | $t = (333,333 * 100)/10^6 = 33$ reactions | $r = 333,333 * 10 * 100 = 333,333,000$ reads for output library |
| 4 | 100-1000 Mb or<br>Whole genome | 400-1200 bp | 2.5 million fragments for 400 bp<br>Therefore, by increasing length, this can be reduced.<br>Eg: 1000 bp fragments would require 1 million unique fragments. | $t = (2.5 * 10^6 * 100)/10^6 = 250$ reactions<br>This can be further optimized to use cells with higher transformation efficiency.<br>Example: a transformation efficiency of $10^7$ would decrease the number of reactions to 25. | $r = 2.5 * 10^6 * 10 * 100 = 2,500,000,000$ reads for output library. |

**Supplemental_Table_S3:**
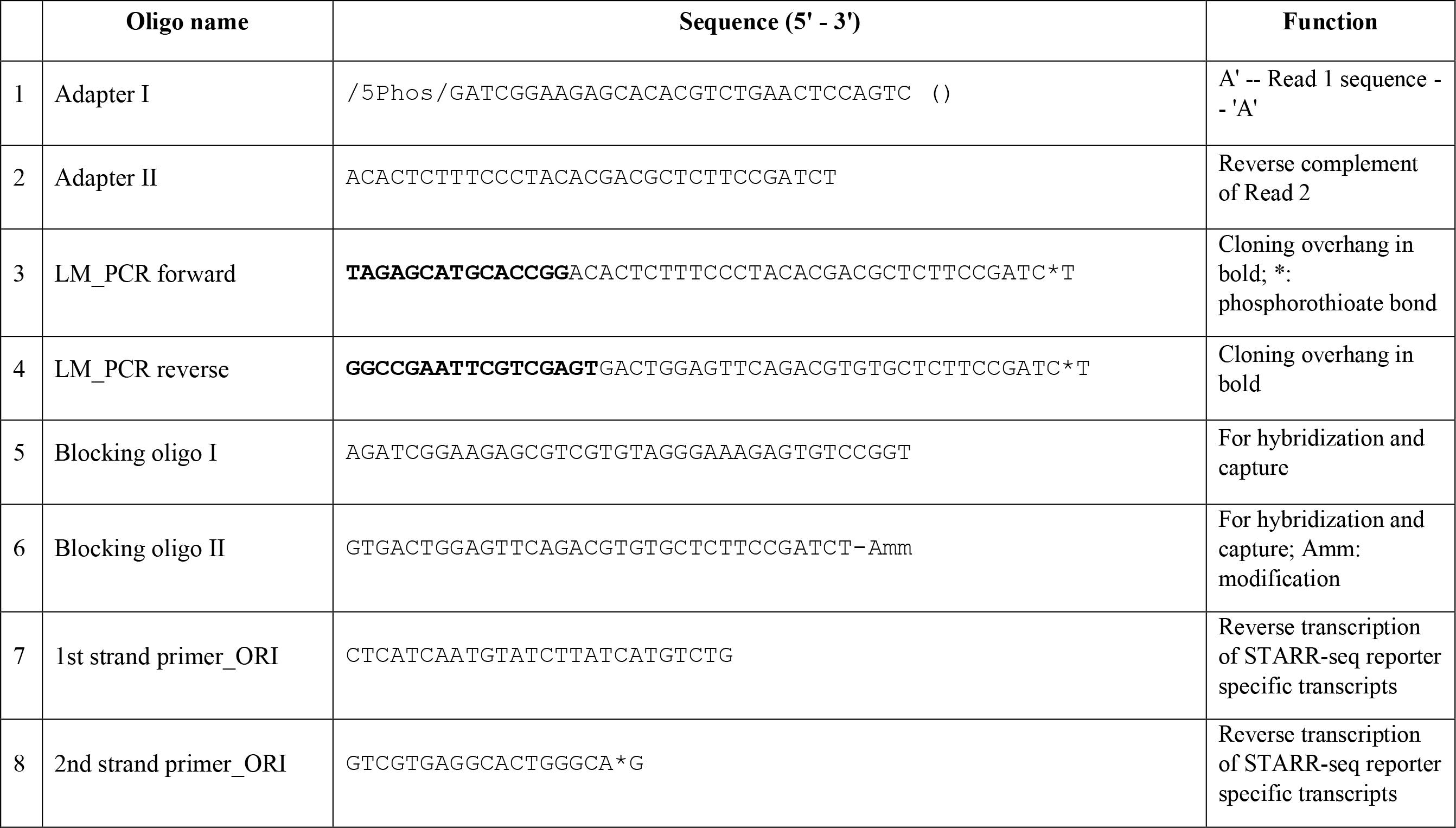
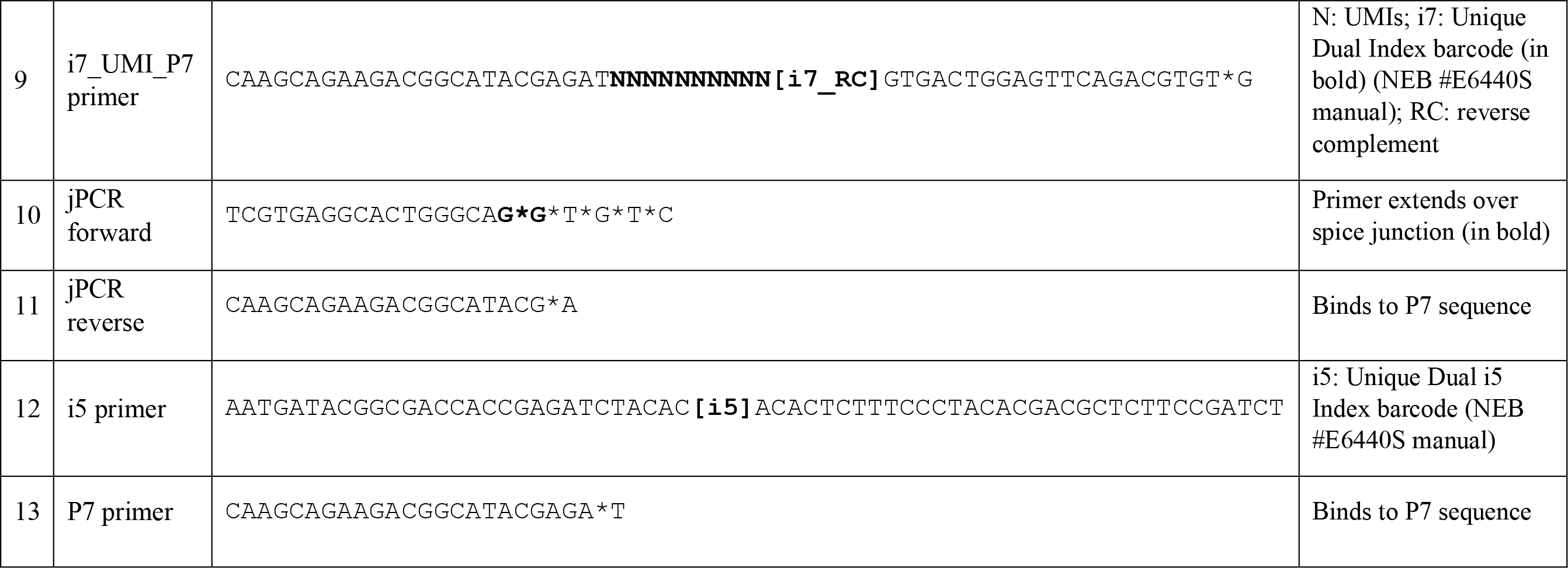
Sequence information for all oligos used for in-house STARR-seq assays is shown. Sequence oligos were ordered from Integrated DNA Technologies, Inc for use. Dilutions required for each oligo is provided in STARR-seq protocol.

**Supplemental_Table_S4A:** Rubric used for scoring each feature of each study for assessing plasmid library information is shown (for Fig. 4 A, B).

| Steps | Rubric for each score based on details provided |  |  |  |  |
| --- | --- | --- | --- | --- | --- |
|  | 0 | 1 | 2 | 3 | 4 |
| **Library size and DNA source** | No detail | Only mentions genome wide/focused. Doesn't connect with goal of experiment | Library target justified and answers research question | Provides details on building target library including source of DNA and enrichment (if focused) | Explains scientific logic and provides complete detail on choosing library size, type, DNA source, enrichment. |
| **Length selection** | No detail | Fragment length | Fragment length and method of size selection | Justification of length and selection | Validation of selection |
| **Insert Preparation** | No detail | Mentions steps involved | Adapter ligation kit and LM_PCR reaction details including primers | Adapter ligation protocol parameters and number of reactions | Complete information on adapter sequences, explanation for choice |
| **Library cloning** | No detail | Insert source and vector name | Kit/reagent used | Total number of reactions, pooling information and purification steps | Insert and vector ratio with concentration and amount and optimization steps |
| **Transformation** | No detail | Transformation methodology | Competent cell name | Amount of competent cell and ligated product per reaction and number of reactions | Validation of transformation, complexity assessment, QC and optimization |
| **Plasmid Library QC** | No detail | Validation of any 1 step or QC checkpoint | Validation of any 2 steps or report both QC checkpoints | Validation of any 3 steps or provide intermediate reports for both QC checkpoints | Validation of all steps and report complete details for both checkpoints |

**Supplemental_Table_S4B:** Rubric used for scoring each feature of each study for assessing library screening information is shown (for Fig. 4 C, D).

|  |  |  |  |  |  |
| --- | --- | --- | --- | --- | --- |
| **Transfection** | No detail | Cell type | Transfection methodology | Transfection protocol | Transfection efficiency, validation and optimization |
| **RNA Isolation** | No detail | Different sub-stages of RNA (total RNA, mRNA prior to RT | RNA extraction kits used for each sub-stage execution | Number of cells used, amount of final RNA, and description of protocol of kits used | Optimization and validation of RNA extracted at each sub-stage, purity and RNA integrity |
| **Reverse Transcription** | No detail | Reaction kit, primer and parameters | Amount of starting material and number of reactions. | Explanation for use of kit parameters | Validation and purification of cDNA obtained and optimizations. |
| **Library screening QC** | No detail | Explanation and validation of any 1 step | Explanation and validation of any 2 steps or report QC checkpoint | Explanation and validation of any 3 steps or report intermediate data for QC checkpoint | Explanation and validation of all steps and provide complete details for checkpoint |

**Supplemental_Table_S4C:** Rubric used for scoring each feature of each study for assessing library sequencing information **(for Fig 4 E, F)**.

|  |  |  |  |  |  |
| --- | --- | --- | --- | --- | --- |
| **Sequencing Library preparation** | No detail | Parameters of sequencing ready PCR reactions | Kit detail and protocol details for sequencing libraries (input and output) | Number of reactions, replicate information | Index information, lane and pooling information, final validation and amount sent for sequencing |
| **Read Information** | No detail | Mention sequencing platform and GEO | Read length, type of sequencing | Adapter type, total number of reads | Complete adapter sequence and reads per replicate |
| **Read QC** | No detail | Read QC steps reported in manuscript or supplementary document | Read QC tools used along with their parameters, raw data processing commands and post processing commands (after peak call) if applicable reported in manuscript or supplementary document | Final read depth or coverage per library reported in manuscript or supplementary document | Read loss due to mapping quality, N bases and bad/off-target reads reported in manuscript or supplementary document and enough information to calculate read depth or library coverage by a reader |
| **Data QC** | No detail | Correlation of replicates within input and/or output libraries reported in manuscript or supplementary document | Correlation across replicates within input and output libraries and details about transfection efficiency reported in manuscript or supplementary document | Correlation of enhancers reads between replicates and/or correlation of fold changes between replicates reported in manuscript or supplementary document | Number of reproducible peaks called across replicates and/or by sub-sampling filtered read file, reported in manuscript or supplementary document |
| **Data transparency** | No detail | Raw data and final enhancer activity data uploaded to public repository e.g. GEO | Filtered/Processed reads (after read QC) uploaded to public repo OR source code provided in public repo to reproduce filtered read file | Intermediate QC data uploaded to public repo OR source code provided in public repo to reproduce intermediate QC files | Complete analysis pipeline starting from raw data processing to analysis post peak calling uploaded to public repo. The source code should be sufficient to reproduce final results and QC data at all intermediate steps |

## STARR-SEQ PROTOCOL (Girirajan lab)

This mammalian (human) STARR-seq protocol is based on the protocol reported by Muerdter and colleagues (Muerdter et al. 2018) and Neumayr and colleagues (Neumayr et al. 2019) from Dr. Alexander Stark’s lab, as well as the SeqCap EZ HyperCap Workflow by Roche (Roche Sequencing Solutions, Inc, CA 94588, USA) <u>along with various modifications and adaptations</u>. The major steps of the protocol are provided below, followed by detailed description of the steps.

**(1)** STARR-seq plasmid library preparation

(A) ) Insert preparation: End repair and dA tailing

(B) ) Insert preparation: Adapter ligation

(C) ) Insert preparation: LM_PCR

(D) Hybridization and Capture

(E) Vector preparation: Vector culturing

(F) Vector preparation: Vector linearization

(G) Library amplification: Library cloning

(H) Library amplification: Transformation

(I) Library amplification: Library storage and extraction

**(2)** STARR Seq screening

(A) ) Culturing of cells

(B) Transfection

(C) Total RNA isolation

(D) mRNA isolation

(E) TURBO DNase treatment

(F) ) RNAClean XP treatment

(G) cDNA library preparation: 1^st^ strand synthesis

(H) ) RNase A treatment and AMPure XP clean-up

(I) UMI addition: 2^nd^ strand synthesis

(J) UMI addition: UMI_PCR

(K) Junction PCR

(L) Sequencing Ready PCR: Output library preparation

(M) Sequencing Ready PCR: Input library preparation

### (1A)#Insert preparation: End repair and dA tailing

#### Before starting

➢ The first section of this protocol follows the SeqCap EZ HyperCap workflow by Roche. This protocol replaces the NEBNext Illumina library preparation protocol reported by Neumayr and colleagues (Neumayr et al. 2019) to incorporate hybridization and capture of the target library using SeqCap EZ Prime Choice XL probes (Roche catalog # 08247510001).

➢ The required reagents for this protocol include KAPA Hyper Prep reaction kit (Roche catalog #KK8500) that consists of End repair and A-tailing buffer, End repair and A-tailing enzyme, ligation buffer, DNA ligase and KAPA HiFi HotStart ReadyMix (2X). This kit also contains primer mixes for standard illumina adapters however, use custom adapter sequences provided in supplementary table S2. Order additional polymerase for subsequent PCR steps KAPA HiFi HotStart ReadyMix (2X) (Roche catalog #07958935001)

➢ Prepare AMPure XP beads (Beckman Coulter catalog #A63881) for sample clean-up in 1.5ml Eppendorf tube (VWR catalog #87003-294) to equilibrate to room temperature for at least 30 mins.

➢ Human whole genome DNA (Promega catalog # G3041) is fragmented through sonication at core facility and then selected at specified length using blue pippin and fragment length distribution is verified on a bioanalyzer and provided in 5 tubes at ∼ 33 ng/µl. Library is sheared and selected to ∼ 500 bp.

➢ Take equal amounts (in ng) from each tube and pool to a total of 260ng DNA starting material for end repair and dA tailing.

➢ Starting DNA: 260 ng made up to 50 µl with ultra-water in 0.2 ml tubes (VWR catalog #20170-012) in 2 replicates.

#### End repair and dA tailing protocol

1. Thaw reagents on ice. Assemble following reaction on ice. Mix thoroughly by pipetting and light tapping. Spin down and keep on ice.

2. Incubate reactions on thermocycler (program: end_repair)

|  | Reagent | Volume |
| --- | --- | --- |
| 1 | KAPA End Repair & A-Tailing Buffer | 7μl |
| 2 | KAPA End Repair & A-Tailing Enzyme mix | 3μl |
| 3 | DNA + water | 50μl |
|  | Total | 60μl |

2. Incubate reactions on thermocycler (program: end\_repair)
|  | Temperature | Time |
| --- | --- | --- |
| 1 | 20°C | 30 mins |
| 2 | 65°C | 30 mins |
| 3 | 4°C | hold |

➢ Continue to adapter ligation without stopping.

### (1B)#Insert preparation: Adapter Ligation

1. Anneal adapter oligos: Resuspend adapter I and II sequences to 100 µM stock (All adapter sequences provided in supplementary table S2). Order fresh oligo sequences from IDT (Integrated DNA Technologies, Inc, IA-52241) for new library. Use NEBuffer2 (New England Biolabs catalog # B7002S).
2. Assemble following reaction in a 0.2 ml tube. 

3. Incubate reaction at 95°C for 5 mins on thermocycler. Keep on bench for 2 hours to cool slowly to room temperature.
4. Add 2 µl ultra-pure water + 3µl adapter mix for final conc at 15 µM for KAPA protocol. (5 µl adapter per reaction in the next step)
5. Thaw out adapter ligation reagents on ice and assemble following reaction in 0.2 ml tubes.

|  | Reagent/Oligo | Volume |
| --- | --- | --- |
| 1 | Adapter I | 10 μl |
| 2 | Adapter II | 10 μl |
| 3 | NEBuffer 2 | 4 μl |
| 4 | Ultra-pure water | 16 μl |
| | Total | 40 μl<br>(25μM) |

6. Mix samples thoroughly by pipetting and spin down. Incubate in thermocycler (program: Adapter lig)

|  | Reagent/Oligo | Volume |
| --- | --- | --- |
| 1 | Ultra-water | 5 μl |
| 2 | KAPA Ligation Buffer | 30 μl |
| 3 | KAPA DNA Ligase | 10 μl |
| 4 | End Repair/dATail product (**Step A**) | 60 μl |
| 5 | Adapters (15 μM) | 5 μl |
| | Total | 110 μl |

7. Clean-up reaction using 0.8X AMPureXP bead clean-up (protocol below).

6. Mix samples thoroughly by pipetting and spin down. Incubate in thermocycler (program: Adapter lig)
|  | Temperature | Time |
| --- | --- | --- |
| 1 | 20°C | 15 mins |

#### AMPure XP Clean-up notes

➢ These are magnetic beads that bind to specific lengths of DNA depending on the ratio of bead volume and sample volume.

➢ Beads must be equilibrated to room temperature prior to use for at least 30 mins.

➢ Use only freshly prepared 80% ethanol (diluted in ultra-pure water) for wash steps.

➢ Ethanol wipe bench and magnetic rack prior to use.

➢ All further AMPure XP clean-up steps will follow the same protocol with different volume ratios and elution reagent type and volume.

#### AMPure XP Clean-up protocol

1. Pipette appropriate amount of beads into sample tube. For 0.8X clean-up, pipette 88 µl beads into each 110 µl adapter ligation reaction and transfer mix to 1.5 ml Eppendorf tube.
2. Mix beads and sample thoroughly by vortexing for 10 secs and pipetting vigorously. Spin down the tube once finished.
3. Incubate mix for 5-10 mins at room temperature. Set-up magnetic rack during incubation and prepare 5 ml 80% ethanol.
4. Place tubes on magnetic rack and incubate for 5 mins (till a clear solution is observed). Carefully remove supernatant liquid without disturbing the beads. (Keep tube on the rack while discarding supernatant)
5. Add 200 µl 80% ethanol and incubate for 30 secs to 1 min and then remove promptly. Repeat this step. After 2 washes, leave tube on rack to dry.
6. If residue ethanol is scattered on the inner tube surface, spin down tube for 1-2 secs and then use P20 pipette to remove trace ethanol. Dry tube for 2-3 mins or till no liquid visible.
7. Do not over-dry. The beads should have a moist liquid coating and not crack up.
8. Remove tube from rack and add 40 µl 10mM Tris-HCl at pH 7.5 for elution. Mix thoroughly with beads and vortex lightly and spin down.
9. Incubate for 5 mins. Place tube back on magnetic rack. Wait till clear solution obtained and pipette solution into a fresh 1.5 ml Eppendorf tube.
10. Measure sample purity and concentration. Typical concentration observed is ∼ 40 ng/µl but purity ratios will be high (>3 for both 260/280 and 260/230).
11. Use 3 µl adapter ligated DNA for subsequent LM_PCR step.

### (1C)#Insert preparation: Ligation Mediated PCR (LM_PCR)

➢ This step adds overhang arms to the adapter ligated fragments to facilitate library cloning.

➢ Use custom primers (LM_PCR forward and LM_PCR reverse) designed for human STARR-seq ORI vector (Muerdter et al. 2018) as provided in supplementary table S2.

➢ Order fresh primers from IDT for new library. Reconstitute oligos to 100 µM.

#### LM_PCR protocol

1. Dilute primers and make primer pool according to the following:

2. Thaw out LM_PCR reagents on ice and assemble following reaction on ice in 0.2 ml PCR strip tubes.

|  | Reagent/Oligo | Volume |
| --- | --- | --- |
| 1 | LM_PCR forward | 2 µl |
| 2 | LM_PCR reverse | 2 µl |
| 3 | Ultra-pure water | 16 µl |
| 4 | Total | 20 µl |

3. Gently mix and spin down and incubate in Thermocycler (Protocol: LM_PCR)

2. Thaw out LM\_PCR reagents on ice and assemble following reaction on ice in 0.2 ml PCR strip tubes.
|  | Reagent/Oligo | Volume |
| --- | --- | --- |
| 1 | Adapter ligated DNA (Previous step) | 3µl |
| 2 | Primer Mix | 5µl |
| 3 | Ultra-water | 17µl |
| 4 | KAPA HiFi HotStart Ready Mix | 25µl |
|  | Total | 50µl |

4. Clean-up reactions using 1.8X AMPure XP bead clean-up. (Use 90 µl beads for each 50 µl reaction. Elute samples in 30 µl of ultra-pure water.
5. Measure concentration and purity. Typical yield is ∼ 10-20 ng/µl.
6. **Checkpoint 1:** Assess samples on bioanalyzer. Send 3 µl of each sample to assess purity and fragment length distribution. Fragments should show a steady increase in length based on the size of the adapter sequences and serves as the first validation of library preparation. This also shows presence of adapter dimers or primer dimers as well as potential PCR biases.
7. Samples can be stored at -20°C.
8. For whole genome libraries, repeat several replicates of LM_PCR as directed in protocol reported by Neumayr and colleagues (Neumayr et al 2019).
9. For focused libraries using hybridization and capture of target sites, perform at least 8 replicates. Assess each replicate for quality and length distribution.
10. Pool equal amounts from each replicate for hybridization and capture.

3. Gently mix and spin down and incubate in Thermocycler (Protocol: LM\_PCR)
|  | Temperature | Time | Step |
| --- | --- | --- | --- |
| 1 | 98°C | 45 secs | Initial denaturation |
| 2 | 98°C | 15 secs | Denaturation |
| 3 | 65°C | 30 secs | Annealing |
| 4 | 72°C | 30 secs | Extension |
| 5 | 72°C | 60 secs | Final extension |
| 6 | 4°C | Hold |  |

### (1D)#Hybridization and Capture

➢ This step allows for selectively capturing target library using custom designed hybridization and capture probes. The required kits needed for this include

(1) SeqCap EZ Prime Choice XL probes (Roche catalog # 08247510001)

(2) SeqCap EZ Hybridization and Wash kit (Roche catalog # 05634261001)

(3) SeqCap EZ Pure Capture Bead Kit (Roche catalog # 06977952001)

(4) Blocking Oligos custom designed (IDT), sequences provided in supplementary table S2

(5) COT Human DNA (Sigma Aldrich catalog #11581074001)

➢ Upon receipt of probes, immediately aliquot into 4.5 µl aliquots in 0.2 ml tubes and store at -20°C. Each aliquot will be sufficient for 1 capture reaction.

➢ To prepare blocking oligos, reconstitute oligos to 400 µM. Use 2.5 µl (1000 pmol) of each for final reaction. Here is a schematic outline on the role of blocking oligos during hybridization and capture.

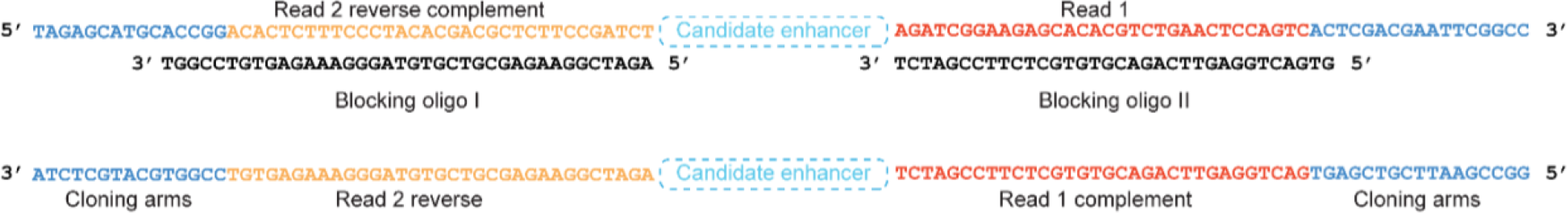

➢ Prepare pooled LM_PCR product from multiple replicates and measure final concentration and purity. Starting material should be ∼ 1 µg of DNA.

➢ Prior to starting, thaw capture reaction aliquot on ice.

➢ Bring AMPure XP beads to room temperature for at least 30 mins for clean-up.

#### Hybridization protocol

1. Prepare following reaction mix in a 1.5 ml Eppendorf tube.

2. Calculate total volume of the mixture and add 2X AMPure XP beads to mix and vortex for 10 seconds and incubate at room temperature for 10 mins.
3. Place tube onto magnetic rack and discard supernatant when it clears.
4. Wash beads with 190 µl 80% ethanol for 30 secs and remove and dry beads for 5 mins.
5. For each reaction, prepare a hybridization buffer mix:
6. µl Hybridization Buffer + 3 µl Hybridization Component A = 10.5 µl
7. Add hybridization buffer mix to beads (per reaction) and vortex and incubate up to 2 mins and place on rack.
8. Pipette complete 10.5 µl solution from tube into fresh 200 µl tube containing 4.5 µl of SeqCap EZ Prime Choice XL probes and mix thoroughly.
9. Incubate on thermocycler (program: Hybridization)

|  | Component | Volume |
| --- | --- | --- |
| 1 | COT Human DNA | 5 µl |
| 2 | Blocking Oligos | 2.5 + 2.5 µl |
| 3 | LM_PCR Pool | 1 µg |

|  | Temperature | Time |
| --- | --- | --- |
| 1 | 95°C | 5 mins |
| 2 | 47°C | 20 hours |

#### Wash and recover of captured library protocol

➢ Follow SeqCap EZ HyperCap workflow (User’s guide v2.3 pages 29 – 34) according to manufacturer’s instructions completely for capturing target library from hybridized beads.

➢ Dilute all buffers according to the protocol.

➢ Prior to starting, allow capture beads and AMPure XP clean-up beads to equilibrate to room temperature for at least 30 mins.

➢ Proceed directly to post capture LM_PCR reaction. All reaction parameters and primers are same as previous LM_PCR.

➢ Use sample and beads as starting material for LM_PCR (∼ 20 µl).

➢ Perform 1.8X AMPure XP clean-up of LM_PCR product. Elute in 50 µl ultra-pure water. Repeat AMPure XP clean-up step and elute in 40 µl. This significantly improves purity ratio of final product.

➢ Product now ready for library cloning.

### (1E)#Vector preparation: Vector culturing

Use human STARR-seq vector ADDgene #99296 as reported by Muerdter and colleagues (Muerdter et al. 2018). The hSTARR-seq_ORI vector was a gift from Alexander Stark (Addgene plasmid # 99296 ; http://n2t.net/addgene:99296 ; RRID:Addgene_99296)

1. Use all standard bacteria culture protocols for preparation of LB broth, LB-agar plates and antibiotic stocks. Prepare fresh broth and plates for use.
2. Prepare LB plates along with both ampicillin (10mg/ml stock) and chloramphenicol (25 mg/ml in ethanol stock) according to standard lab protocol. (final ampicillin concentration is 100 µg/ml and chloramphenicol is 25 µg/ml in plate) (Vector carries both resistance markers)
3. Streak LB plate using a sterile pipette tip with inoculum from vector stab and incubated overnight at 37°C to grow isolated colonies.
4. Following day, pick multiple colonies using sterile pipette tips and inoculate 4 ml LB broth with ampicillin and chloramphenicol in 14 ml bacteria culture tube (VWR catalog #60819-524) (1 culture tube per colony) and shake at 300 rpm at 37°C overnight in shaker-incubator.
5. Use 1 ml of culture broth to make glycerol stock for each colony (Use standard procedure for glycerol stock preparation). Spin down rest of the cultures and extract vector DNA using ZymoPURE plasmid Miniprep kit (Zymo catalog #D4209).
6. Measure purity and concentration and send 2 samples for sanger sequencing at the genomics core facility using multiple primers designed for ORI vector sequence verification to verify sequence. (Sequences available on request)
7. Following sequence verification regrow sample with highest concentration and purity from the glycerol stock. In the morning, pick glycerol stock with a sterile pipette and inoculate 2 ml of LB broth with ampicillin and chloramphenicol and incubate at 300rpm at 37C for 8 hours as a starter culture. In the evening, use 1 ml of starter culture to incubate 150 ml of LB broth with ampicillin and chloramphenicol and incubate overnight at 300 rpm at 37°C.
8. Make fresh glycerol stocks using 1ml of culture. Spin down rest of the culture and extract vector DNA using ZymoPURE II plasmid Midiprep kit (Zymo catalog #D4200) and verify concentration and purity.

### (1F)#Vector preparation: Vector linearization

➢ Reagents required include enzymes SalI-HF (NEB catalog #R3138S) and AgeI-HF (NEB catalog #R3552S) along with supplied Cutsmart buffer and 6X purple loading dye.

➢ Perform gel extraction using Zymoclean Gel DNA Recovery Kit (Zymo catalog #D4001). Further PCR purification or sample concentration can be carried out using DNA clean and concentrator kit (Zymo #D4003).

➢ Bring AMPure XP beads to room temperature for at least 30 mins.

➢ Prepare 1% agarose gel.

#### Vector linearization protocol

1. Setup following reaction for each digest. Set-up 8 such reactions in 0.2ml PCR strip tubes and incubate in thermocycler at 37°C for 2 hours. Approximately 1/3^rd^ of the product will be recovered from the gel so scale reactions accordingly for larger libraries.

2. Run on 1% agarose gel for 1 hour at 100V till bands separate and cut out heavier band (∼2kb) and weigh each slice. Add 3 volumes of ADB buffer and melt gel at 55°C for 30 min. Column purify each gel slice separately and pool 4 samples together and measure concentration and purity. (Purity may be low).
3. Perform secondary clean-up with 1X AMPure XP beads for each pooled sample and confirm concentration. (Should be above 35ng/µl)

|  | Digest mix | Volume |
| --- | --- | --- |
| 1 | Vector | ~ 1µg |
| 2 | SalI | 1µl |
| 3 | AgeI | 1µl |
| 4 | Cutsmart | 5µl |
| 5 | Water | Up to 50µl |
|  | Total | 50µl |

### (1G)#Library amplification: Library cloning

➢ For cloning, use the NEBuilder HiFi DNA Assembly Master Mix (NEB catalog #E2621L) with the 2-3 fragment assembly protocol for efficient cloning. Alternatives include In-fusion HD cloning and Gibson Assembly.

➢ Try multiple ligation ratios to optimize library cloning. Repeat cloning using multiple replicates to maximize library complexity.

➢ Use DNA clean and concentrator kit or AMPure XP beads for reaction clean-up prior to transformation.

#### Library cloning protocol

1. Calculation of [insert: vector ratio] for ligation Considerations:

a. Vector and insert mass and associated molarity
b. Maximum molar capacity per reaction (30 fmol – 200 fmol per reaction for ligating 2-3 fragments)
c. Final volume of reaction
d. Number technical replicates to be performed For calculations:

➢ Insert length: Use average insert fragment length as observed on bioanalyzer in step C.

➢ Vector length: 2543 bp

Ratio calculations can be done using the NEBioCalculator (https://nebiocalculator.neb.com/#!/ligation) to calculate amount of DNA for insert and vector required for final reaction assembly. Example parameter calculation for cloning:

Starting material:

| DNA | conc ng/ul | 260/280 | 260/230 |
| --- | --- | --- | --- |
| Insert_1 | 29.8 | 1.81 | 2.32 |
| Insert_2 | 25.7 | 1.81 | 2.22 |
| Vector | 38.2 | 1.87 | 2.33 |

Cloning parameters:

| Ratio | Vector (ng) | Vector (fmol) | Insert (ng) | Insert (fmol) | total fmol |
| --- | --- | --- | --- | --- | --- |
| 1:2 | 100 | 63.64 | 51.91 | 127.3 | 190.94 |
| 1:2 | 50 | 31.82 | 25.95 | 63.62 | 95.44 |
| 1:2 | 70 | 44.54 | 36.34 | 89.1 | 133.64 |
| 1:7 | 39.29 | 25 | 71.38 | 175 | 200 |

1. Set up cloning reactions according to the following volumes. Perform at least 4 reactions for each condition.

2. Incubate reactions at 50°C for 1 hour. Pool 4 reactions of same conditions and purify using 1X AMPure XP beads and measure concentration and purity.

- 625 2. Set up cloning reactions according to the following volumes. Perform at least 4 reactions for each condition.
|  | Sample | Amount (volume) |
| --- | --- | --- |
| 1 | Insert | X (depends on ratio) |
| 2 | Vector | Y (depends on ratio) |
| 3 | NEBuilder HiFi Assembly mix | Up to 10ul |
| 4 | Water | 10 |
| 5 | total | 20 ul |

- 628 3. Incubate reactions at 50°C for 1 hour. Pool 4 reactions of same conditions and purify using 1X AMPure XP beads and measure concentration and purity.
| Ligation reactions | conc ng/ul | 260/280 | 260/230 | Ratio | # Reactions |
| --- | --- | --- | --- | --- | --- |
| HiFi Assembly_1 | 24.1 | 1.92 | 1.5 | 1:7 | 4 |
| HiFi Assembly_2 | 36 | 2 | 1.87 | 1:2 | 4 |
| HiFi Assembly_3 | 33.4 | 1.87 | 1.48 | 1:2 | 4 |
| HiFi Assembly_4 | 31.8 | 1.88 | 2.22 | 1:2 | 4 |
| HiFi Assembly_5 | 42.5 | 1.91 | 2.21 | 1:2 | 4 |
| HiFi Assembly_6 | 41.6 | 1.87 | 2.21 | 1:2 | 4 |
| HiFi Assembly_7 | 42 | 1.89 | 2.27 | 1:2 | 4 |

### (1H)#Library amplification: Library transformation

➢ Use either NEB5 alpha (NEB #C2989, currently discontinued) and NEB10 beta (NEB #C3020K) electrocompetent cells. Following protocol has been tested with only NEB electrocompetent cells.

➢ Perform test transformation for each cloning ratio in 25 μl competent cells. Vary between 2 –5 μl of cloned product per transformation reaction.

➢ For control, use supercoiled plasmid of choice, either plasmid provided with NEB electrocompetent cells (pUC19 Vector at 50 pg/μl or plasmid PX459 (Ran et al. 2013) (ADDgene catalog #62988). Vector pSpCas9(BB)-2A-Puro (PX459) V2.0 was a gift from Feng Zhang (Addgene plasmid # 62988 ; http://n2t.net/addgene:62988 ; RRID:Addgene_62988) For PX459, dilute stock to 1:10 and use 2 μl per transformation.

➢ Prior to starting: Electroporation is a very time sensitive step and must be carried as quickly and efficiently as possible. Make sure everything is kept ready, labeled and within reach. After cells have thawed out, work as quickly as possible to proceed. If there are a large number of reactions, perform and thaw in batches of 4 or 8 reactions.

➢ Prepare adequate number of sterile LB-agar ampicillin plates according standard laboratory protocol for estimating cloning efficiency and CFU/μg. Prepare >20 plates from 500 ml LB-agar. Pre-warm to 37°C during transformation.

➢ Prepare adequate amount of sterile LB broth with ampicillin for inoculating and amplifying transformed library. Prepare 500 ml to 1 L LB broth depending on number of transformations performed. Use >600 ml for 100 μl (4 reactions) to 200 (8 reactions) μl of competent cells.

➢ Pre-chill electroporation cuvettes 200 μl and 20 μl tips **overnight** (or at least 1 hour) at 4°C.

➢ Tightly pack ice into an ice bucket and spray with ethanol to make a ‘chilled ice sludge’.

Move cuvettes into ice bucket.

➢ Pre-warm SOC media provided with electrocompetent cells to 37°C for 1 hour and place back on bench.

➢ Take out required number of sterile 1.5ml Eppendorf tubes (1 for each transformation reaction) and place on ice to chill.

➢ Thaw out electrocompetent cells on the ice. Gently flick tube to check thawing.

➢ Label each tube with name of transformation reaction.

➢ Label 14 ml round bottomed bacterial culture tubes with similar labels as tubes.

➢ Make sure to work inside a sterile environment (under a lamp or inside a biosafety cabinet).

Wipe everything down with ethanol prior to starting transformation.

➢ Ensure electroporation machine is within reach and set to correct parameters depending on competent cells. Use 1700V for NEB5 alpha or NEB10 beta cells.

#### Transformation procedure

1. After cells have completely thawed out, pipette out 25 μl of cells onto the pre-chilled and labeled Eppendorf tube (DO NOT pipette more than once or necessary). Use pre-chilled tips while pipetting.
2. Add appropriate amount of cloned product (2 – 5 μl) to cell. Gently mix by flicking tube from side by holding the tube around the cap a few times. Use pre-chilled tips. (High number of unique colonies observed for 5 μl of product in 25 μl of cells)
3. Transfer mix of cells and cloned product into electroporation cuvette without any bubbles. Wipe cuvette from the side and electroporate cells with 2 pulses. Record the time constants for each transformation. (Typical range is 4.4 – 4.6 milliseconds).
4. Immediately add 975 ml of pre-warmed SOC media to cuvette and pipette up and down to mix.
5. Transfer the transformant mixture into pre-labeled 14 ml bacterial culture tubes and incubate in a shaker-incubator at 300 rpm at 37°C for 1 hour.
6. Repeat this for each transformation reaction one by one.
7. During 1 hour incubation, prepare LB broth and plates for overnight culture. Label plates with name of cloning ratio being tested and serial dilution number.
8. Label flasks with cloning ratio being amplified. Keep separate flasks for different ratios. If ratios show similar efficiency, products can be pooled during extraction.
9. Prepare 100% ethanol and sterile spreader for plating. Prepare 1.5 ml Eppendorf tubes for serial dilution. Add 900 μl of LB broth (no antibiotics) to each tube.
10. After 1 hour incubation, take 14 ml tubes off the shaker-incubator. Transformant sample should be cloudy.
11. Use 1 reaction per ratio to test efficiency in dilution series. Take 100 μl of transformant and add to 900 μl of LB broth in the 1.5ml Eppendorf tube for the 1^st^ dilution (labeled as [-2]) and mix thoroughly.
12. Change tips and take 100 μl of 1^st^ dilution and add to another 900 μl LB broth in the next tube for the 2^nd^ dilution (labeled as [-3]. Repeat this up to 5^th^ or 6^th^ dilutions (labeled [-6] and [-7] respectively). Repeat this for each cloning ratio and control plasmid.
13. From each tube, take 100 μl of diluted transformant and spread evenly onto the pre-warmed plates and incubate overnight at 37°C.
14. Add rest of the transformants to the LB broth in conical flasks with ampicillin. Make sure volume of broth is ∼ ¼ of maximum volume of flask. Use 500 ml flasks for 150 ml cultures. Use 1 L flasks for 250 ml cultures. Pool replicates of same cloning ratios.
15. Incubate cultures overnight on shaker-incubator at 300 rpm and 37°C.

### (1I)#Library amplification: Library storage and extraction

1. After incubation between 12-16 hours check for colonies on LB plates. Depending on the cloning and transformation efficiency, there should be at least >1 colony on the [-6] plate for all successfully transformed libraries. DH10B should show increased number of colonies.
2. This can be used to compare efficiency of each cloning ratio and show overall estimates of library complexity. A single colony on [-6] is equivalent to at least 1 million unique colonies. Higher number of colonies indicates greater library complexity.
3. Control plasmid should show significantly more colonies than cloned products and can be used to calculate the Colony Forming Units (CFU)/ μg of plasmid transformed using the formular provided by NEB. This helps estimate the transformation efficiency of the competent cell being used.
4. After validation of transformation, check the optical density of the cultures (∼ 2.6).

#### Library Storage

1. Pipette 5 ml of culture into a sterile round-bottomed 15 ml tube and spin down culture for 5 mins.
2. Pipette out LB broth and resuspend pellet in 750 ml of fresh LB broth. Mix with 750 ml of sterile glycerol and transfer to 2 ml screw-capped cryotube. Label tubes and transfer to -80°C for long term storage.
3. Make up to 5 glycerol stocks for each successfully transformed ratio.

#### Library extraction

1. Pipette remaining cultures into screw capped 500 ml centrifuge tubes and spin down. Aspirate media and use pellet to extract DNA using ZymoPURE II Plasmid Maxiprep kit (Zymo catalog #D4203) according to manufacturer’s protocol.
2. Use up to 150 ml of culture per maxiprep reaction. Use centrifuge version of the protocol.
3. Perform optional endotoxin treatment on the extracted library using column provided in kit.
4. Verify concentration and purity of final library ready for transfection or input library preparation.

### (2)#STARR-seq Screening (2A) Culturing of cells

➢ Follow standard cell culture techniques for all experiments. Conduct all cell culture inside biosafety cabinet. Use only cell culture grade labware including sterile serological pipettes (VWR catalog #89130-896 for 5 ml, catalog #89130-898 for 10 ml, catalog #89130-900 for 25 ml and catalog #414004-265 for 2 ml aspiration pipettes) and standard cell culture plates and culture dishes.

➢ Use HEK293T cells or cell line of choice for STARR-seq. Do appropriate literature review to see evidence of IFN response from cell line upon transfection. HEK293T cells have not shown to be highly susceptible.

➢ Make appropriate media for HEK293T cells using following recipes.

|  | Media | percentage | Vol (500 ml) | Catalog number |
| --- | --- | --- | --- | --- |
| 1 | DMEM (High glucose) | 87 | 435 | Sigma Aldrich #D6429-500ML |
| 2 | PenStrep | 1 | 5 | Sigma Aldrich #P4333 |
| 3 | FBS | 10 | 50 | Sigma Aldrich #F2442 |
| 4 | Non-essential amino acids | 1 | 5 | Sigma Aldrich #M7145-100ML |
| 5 | HEPES | 1 | 5 | Sigma Aldrich #H0887-100ML |

➢ For 500 ml solutions, take out 65 ml of DMEM from the bottle and prepare media inside DMEM bottle.

➢ Store FBS in 40-50 ml stocks in 50 ml tubes (VWR catalog #89039-656) and store at -20°C.

Thaw out FBS overnight at 4°C.

➢ Store PenStrep at -20°C in 5-10 ml stocks in 15 ml tubes (VWR catalog #89039-666) and thaw out at room temperature.

➢ Scale recipe as needed.

➢ Thaw cells in to 6-welled plates (VWR catalog #10861-696) or a 6 cm dish (VWR catalog

#25382-100) using standard lab protocol.

➢ Passage cells every 2-3 days or when ∼80% confluent and track cell morphology. Pass cell through at least 3 passages prior to transfection.

➢ For transfection of STARR-seq library, use at least 30 to 50 million cells per replicate (1 x 15 cm dish (VWR catalog #430599). For efficiency, passage cells from 2 or 3, 80% confluent 10 cm dishes (VWR catalog #25382-166) per 15 cm dish.

### (2B)#Transfection

Use Lipofectamine 3000 reagent (Thermo Fisher catalog #L3000008) for HEK293T cells with OptiMEM (Thermo Fisher catalog #31985070). Transfection optimized and shown to be >70% efficient and up to 90%.

1. Ensure ∼70-80% confluency and healthy morphology of cells prior to starting transfection.
2. Pre-warm optiMEM media to 37°C in water bath. Label 1.5 ml sterile Eppendorf tubes to be used for transfection. Keep everything inside biosafety cabinet.
3. Aspirate DMEM culture media from the dishes to be transfected, wash cells with 10 ml sterile 1X PBS (Sigma Aldrich catalog #806552-500ML), aspirate and add 30 ml of OptiMEM media to cells, 30 mins prior to transfection and place cells back into the incubator. (While aspirating, be careful not to detach cells from the surface. Add media dropwise or from the side of dish to prevent detachment.)
4. Scale lipofectamine 3000 protocol according to number of cells or size of culture dish/plate. For 15 cm dish, use following reagent volumes. Remaining protocol steps are unchanged.
5. Perform up to 3 transfections per cell line. To avoid batch effects, transfect replicates on separate days.

6. Prepare tubes A and B and add mix from tube B into tube A dropwise, gently mix by tapping (DO NOT vortex) and incubate for 15 mins.
7. After incubation, gently pipette combined mix onto the cells and incubate for 24 hours.

|  | Tube A | Vol |
| --- | --- | --- |
| 1 | Lipofectamine 3000 | 57 μl |
| 2 | OptiMEM | 1443 μl |
| 3 | total | 1.5 ml |

|  | Tube B | Vol |
| --- | --- | --- |
| 1 | STARR-seq plasmid library | 38 μg |
| 2 | P3000 enhancer reagent | 76 μl |
| 3 | OptiMEM | 1375 μl |
| 4 | Total | 1.5 ml |

### (2C)#Total RNA Isolation

➢ Any RNA isolation kit may be used and should be scaled according to number of cells and culture volume. Use Trizol plus RNA purification kit (Thermo Fisher Catalog #12183555) for HEK293T cells. This kit combines trizol and chloroform extraction with PureLink RNA Mini Kit.

➢ RNA isolation should be carried out inside fume hood. Wipe everything down with RNaseZap RNase Decontamination Solution (Thermo Fisher Catalog # AM9782) and use separate tips and pipettes for RNA. Eppendorf tubes should be clean, RNAse free and labeled for each replicate.

➢ Move tabletop centrifuge to cold room if temperature-controlled centrifuge not available at least 1 hour prior to starting and wipe down with RNAseZap.

➢ Harvest cells in trizol in the biosafety cabinet and then move to fume hood. Trizol and chloroform are biohazard chemicals and all associated waste should be discarded separately and disposed only in satellite waste area inside fume hood.

#### Total RNA Isolation Protocol

1. For 15 cm dish, use up to 6 ml of trizol.
2. To collect cells, aspirate out all media from cells and wash the cells with 10 ml of PBS.
3. Add 6 ml trizol to cells directly and collect ∼1 ml of cells in trizol per 1.5 ml Eppendorf tube (6 tubes per replicate) and incubate for 5 mins at room temperature.
4. Add 0.2 ml of chloroform to each 1 ml of cells and trizol mix vigorously by shaking the tube for 15 secs. Make sure tube cap is firmly closed and not leaking. Incubate at room temperature for 3 mins.
5. Centrifuge tubes at 12,000 g for 15 mins at 4°C. Gently place tubes back inside fume hood. Sample will form 2 phases inside tube. Lower phase will be red phase comprising of phenol- chloroform, an interphase and upper aqueous phase will comprise of RNA.
6. Carefully pipette 450 – 500 μl of aqueous phase into fresh Eppendorf tubes without disturbing other phases. Pipette 150 μl at a time for ease.
7. Add equal volume (450 – 500 μl) of 70% ethanol to each tube, vortex and spin down.
8. Add up to 700 μl of mix into spin column and centrifuge at 12,000 g for 30 secs at room temperature and discard flow-through. Repeat process until all sample has been processed using the same column for all tubes per replicate. (Use the same column for all 6 tubes per replicate).
9. After all sample has been processed per replicate add 700 μl wash buffer I and centrifuge at 12,000 g for 30 secs and discard flow-through.
10. Add 500 μl of wash buffer II and repeat centrifugation and discard flow-through. Repeat step.
11. Perform a final dry centrifuge spin at 12,000 g for 1 min to remove trace ethanol from the column.
12. Remove collection tube and place column into fresh tube for RNA collection. Add 100 μl of DEPC water to column and incubate sample for 1 min at room temperature.
13. Centrifuge tube at 12,000 g for 2 mins at room temperature and measure concentration and purity.
14. Add 100 μl DEPC water for 2^nd^ elution and 3^rd^ elution by repeating process and measure concentration and purity of each elution. If values are consistent, pool eluates and measure final concentration and purity. Purity ratio for 260/280 and 260/230 should be above 2.0.
15. Send samples for RIN analysis to genomics core facility for quality assessment.
16. RNA can be stored at -80°C or used for mRNA isolation.
17. Dilute total RNA to 750 ng/μl for mRNA isolation. Typical yield is ∼1200 μl of dilute total RNA (∼900 μg of total RNA).

### (2D)#mRNA Isolation

➢ Use dynabeads mRNA Purification Kit (Thermo Fisher catalog #61006) for mRNA isolation. Each kit contains reagents for 10 mRNA isolations.

➢ However, binding buffer and wash buffer B (Thermo fisher catalog #11900D) and elution buffer (Thermo Fisher catalog #A33566) will not be enough if beads are to be reused and can be ordered or prepared additionally. Binding buffer is not available separately and needs to be prepared or ordered with kit.

➢ Additional Lysis/Binding buffer (Thermo Fisher catalog #A33562) also needs to be ordered for reusing beads). Instructions are provided on manufacturer website as well as in STARR-seq protocol by Neumayr and colleagues (Neumayr et al 2019).

➢ Dilute total RNA sample to 750 ng/μl prior to starting mRNA isolation. (Beads can process up to 75 μg of RNA at a time and can be regenerated for reuse. Process 100 μl of diluted total RNA at a time.

➢ Use 200 μl of dynabeads per 100 μl of total RNA. Regenerate beads up to 6 times. Therefore, use ∼ 600 μl of total RNA for 1 x 200 μl dynabead reaction. Reuse beads within same replicate to avoid cross contamination.

➢ Work inside RNA hood and wipe everything with RNAseZap.

➢ Set heat block A to 65°C and heat block B to 80°C.

➢ Set tabletop shaker incubator to 25°C (room temperature) and 60 rpm.

➢ Thaw out total RNA on ice before proceeding.

#### mRNA Isolation Protocol

1. Separate the diluted total RNA into 100 μl batches in individual 1.5 ml Eppendorf tubes and keep on ice.
2. Transfer tube containing the sample being processed to 65°C for 2 mins (to remove secondary structures) and then immediately replace on ice.
3. To prepare beads, pipette 200 μl of the beads to a RNAse free 1.5 ml Eppendorf tube and place on magnetic rack for 30 secs. Carefully discard the supernatant without disturbing the beads adhering to the magnet.
4. Remove tubes from the rack and add 100 μl of binding buffer to the tube, lightly mix and place tubes back on rack. Remove supernatant. Add another 100 μl of binding buffer (or same volume of total RNA being processed) and mix. Beads are not ready for mRNA isolation.
5. Add 100 μl of total RNA to 100 μl of beads and mix by pipetting. Incubate on shaker incubator at room temperature and low rpm (∼60 rpm) for 5 mins.
6. Transfer tubes to magnetic rack and remove supernatant. Add 200 μl of wash buffer B, mix and place tubes on magnetic rack. Remove supernatant and repeat wash step.
7. Remove the tubes from the rack and add 6-10 μl of elution buffer per 100 μl total RNA and incubate samples at 80°C for 2 mins and immediately place tubes back on rack.
8. Pipette supernatant (mRNA) into a fresh tube and measure concentration and purity.
9. To reuse the beads, remove the beads from the rack and add 300 μl of Lysis/Binding buffer to wash beads. Place back on the rack, remove the supernatant and add 100 μl binding buffer to proceed with the next round of mRNA isolation for the same replicate. (Note, this is only for reuse of beads and not regeneration which has a separate protocol provided on manufacturer’s website).
10. Repeat until all total RNA has been processed to mRNA. Pool multiple batches for each replicate and measure final concentration and purity. Typical yield is between 45 – 70 μl of mRNA at ∼ 400 ng/μl or 18 – 28 μg of mRNA and depends on starting amount of total RNA (around 1 – 5% of total RNA).
11. Store mRNA at -80°C or proceed with TURBO DNase treatment, RNA clean-up and reverse transcription.

### (2E)#TURBO DNase treatment

➢ This is required to remove any residual DNA from sample prior to reverse transcription. Use TURBO DNA-free Kit (Thermo Fisher catalog #AM1907).

➢ Starting material is around ∼400 ng/μl or ∼400 μg/ml and thus requires the rigorous protocol for TURBO DNase treatment and thus use 2 μl or 4 units of TURBO DNase enzyme per reaction.

➢ Process entire sample together per replicate. Scale protocol according to volume of mRNA.

➢ Set heat block to 37°C prior to starting.

#### TURBO DNase treatment protocol

1. Add 0.1 volume of 10X TURBO DNase buffer to sample.
2. Add 2 μl (4 units) of TURBO DNase enzyme and mix gently (DO NOT vortex).
3. Incubate at 37°C for 60 mins.
4. Add 0.2 volumes of DNase inactivation reagent and mix gently (DO NOT vortex) and incubate at room temperature for 5 mins. Lightly flick tube to redistribute reagent every 1 min.
5. Centrifuge tubes at 10,000 g for 1.5 mins at room temperature and transfer supernatant to fresh 1.5 ml Eppendorf tube for the next step.

### (2F)#RNAClean XP treatment

➢ Perform clean-up of each TURBO DNase reaction using RNAClean XP beads (Beckman Coulter catalog #A63987)

➢ Warm beads to room temperature for at least 30 mins.

➢ Prepare fresh 80% ethanol for sample wash.

➢ Set tabletop shaker incubator to 37°C and 60 rpm (low rpm).

➢ Process 1 replicate per clean-up reaction. Scale volume of beads according to volume of mRNA.

#### RNAClean XP protocol

1. Add 1.8 volume of bead per 1 volume of mRNA sample and mix by thoroughly pipetting >20 times and incubate at room temperature for 15 mins.
2. Transfer mix to magnetic rack and incubate for 10 mins or till when all beads have adhered to magnet.
3. Remove supernatant (should be clear liquid) and add 500 μl of 80% ethanol for 1^st^ wash. Incubate for up to 2 mins, remove supernatant and add 500 μl of 80% ethanol for 2^nd^ wash. Incubate for 2 mins and remove without disturbing beads.
4. Dry beads completely at room temperature for 5 mins. Check for any residual ethanol inside tube and remove.
5. Remove tubes from the rack, and add 20 μl DEPC water to beads for elution, mix by pipetting and vortexing and incubate in tabletop shaker incubator at 37°C at low rpm for 3 mins and place tubes back on magnetic rack.
6. Incubate for 1 min on the rack and pipette pure mRNA into a fresh 1.5 ml Eppendorf tube for further processing.
7. Assess final concentration and purity of mRNA. This is the final checkpoint for samples prior to cDNA library preparation.
8. Typical yield is ∼12 μg of mRNA per replicate (600 – 900 ng/μl of mRNA in 19 μl of water (1 μl used for purity and concentration assessment)

### (2G)#cDNA library preparation: 1^st^ Strand Synthesis

➢ For 1^st^ strand synthesis, use Superscript III kit

➢ Divide total sample amount by 5 and round off to nearest multiple of 5 to determine number of reactions required per sample. Use 5 reactions using 2.4 μg of mRNA per reaction for ∼12 μg samples.

➢ Adjust all samples to 12 μg in 20 μl (600 ng/μl) of DEPC water for uniformity across replicates. Use 4 μl (2.4 μg mRNA) per RT reaction.

➢ Though protocol suggests using 500 ng of mRNA at a time for 1^st^ strand synthesis, we are performing reverse transcription only for fraction of mRNA that was self-transcribed due to enhancer activity and so taking larger amounts of mRNA is permissible.

➢ Dilute STARR-seq reporter specific primer to 2 μM. Dilute dNTP solution to 10 μM.

#### 1^st^ Strand Synthesis Protocol

1. Assemble following reaction as reaction 1:

2. Incubate at 65°C for 5 mins followed by 4°C for 1 min on a thermocycler (program RT_1)
3. Assemble following reaction as reaction II:

|  | Reagent/sample | Vol (µl) |
| --- | --- | --- |
| 1 | mRNA | 4 |
| 2 | dNTP | 1 |
| 3 | Primer | 1 |
| 4 | Water | 7 |
| 5 | Total | 13 |

4. Add reaction II to reaction I for each RT reaction (final volume 20 μl)
5. Incubate according to following conditions on thermocycler (program RT_II):

2. Incubate at 65°C for 5 mins followed by 4°C for 1 min on a thermocycler (program RT\_1) 3. Assemble following reaction as reaction II:
|  | Reagent | Vol (µl) |
| --- | --- | --- |
| 1 | 1<sup>st</sup> strand buffer | 4 |
| 2 | DTT | 1 |
| 3 | RNaseOUT | 1 |
| 4 | Superscript III | 1 |
| 5 | total | 7 |

|  |  |  |
| --- | --- | --- |
| 1 | 50°C | 60 mins |
| 2 | 75°C | 15 mins |
| 3 | 4°C | hold |

### (2H)#cDNA clean-up: RNase A treatment and AMPure XP clean-up

➢ Treat all reverse transcribed samples with RNase A to remove residual RNA.

➢ Pool 5 RT reactions (total = 100 μl per replicate) (5 reactions carried out per replicate)

➢ Move to DNA bench to avoid pipetting RNase A in RNA isolation bench

➢ Prepare 80% ethanol for AMPure XP bead clean-up wash steps

➢ Bring beads to room temperature for at least 30 mins prior to starting.

#### RNase A treatment and AMPure XP bead clean-up Protocol

1. Add 1 μl of RNase A (10mg/ml) per 5 RT reactions.
2. Incubate at 37°C for 1 hour in thermocycler (program: RNase A treatment)
3. Perform **1.4X AMPure XP bead clean-up** protocol
4. Elute cDNA in 43 μl DEPC water for next step.
5. Measuring cDNA concentration is advised but is not accurate and hence it is also okay to proceed to either 2^nd^ strand synthesis and UMI_PCR or directly to jPCR steps.

### (2I)#UMI addition: 2^nd^ strand synthesis

➢ Unique Molecular Identifiers (UMIs) are essential to label and filter out PCR duplicates from the libraries that occur during library preparation.

➢ For addition of UMIs, follow UMI-STARR-Seq protocol reported by Neumayr and colleagues (Neumayr et al. 2019) including a 2^nd^ strand synthesis reaction and UMI_PCR reaction prior to jPCR.

➢ Start 2^nd^ strand synthesis using 2^nd^ strand primer as shown by Neumayr and colleagues (Neumayr et al. 2019). Dilute primer to 10 μM in DEPC water for use.

➢ Use KAPA 2X HiFi Hot Start Ready mix polymerase (Roche catalog #07958927001)

#### 2^nd^ strand synthesis protocol

1. Process entire replicate cDNA (∼43 μl) per reaction. Set-up following reaction for each replicate:

2. Incubate reactions in thermocycler (program: 2^nd^ strand synth)

| | Sample/reagent | Vol (μl) |
| --- | --- | --- |
| 1 | cDNA | 42.5 |
| 2 | KAPA HiFi 2X Ready mix | 50 |
| 3 | 2<sup>nd</sup> strand primer | 7.5 |
| 4 | Total | 100 |

3. Reaction purification using 1.4X AMPure XP bead clean-up.
4. Elute library in 43 μl of DEPC water for UMI_PCR step.

Incubate reactions in thermocycler (program: 2<sup>nd</sup> strand synth)
|  | Temperature | Time | Step |
| --- | --- | --- | --- |
| 1 | 98°C | 60 secs | Denaturation |
| 2 | 65°C | 30 secs | Annealing |
| 3 | 72°C | 30 secs | Extension |

### (2J)#UMI addition: UMI_PCR

➢ To add UMI, use custom designed i7-UMI-P7 primer to add i7 index sequence and UMI sequence simultaneously to cDNA. This helps retain unique dual indexing to mitigate index hopping as well as allow for PCR duplicate filtration as opposed to Neumayr and colleagues protocol where the i7 is replaced by the UMI.

➢ Dilute primers to 10 μM for use. Keep record of which index is being added to which sample and label all tubes.

➢ Process complete cDNA for each reaction.

#### UMI_PCR protocol

1. Set-up following reaction for each replicate:

2. Incubate reaction in thermocycler (program: 2^nd^ strand synth) Same conditions as 2^nd^ strand synthesis.

| | Sample/reagent | Vol (μl) |
| --- | --- | --- |
| 1 | cDNA | 42.5 |
| 2 | KAPA HiFi 2X Ready mix | 50 |
| 3 | i7-UMI-P7 primer | 7.5 |
| 4 | Total | 100 |

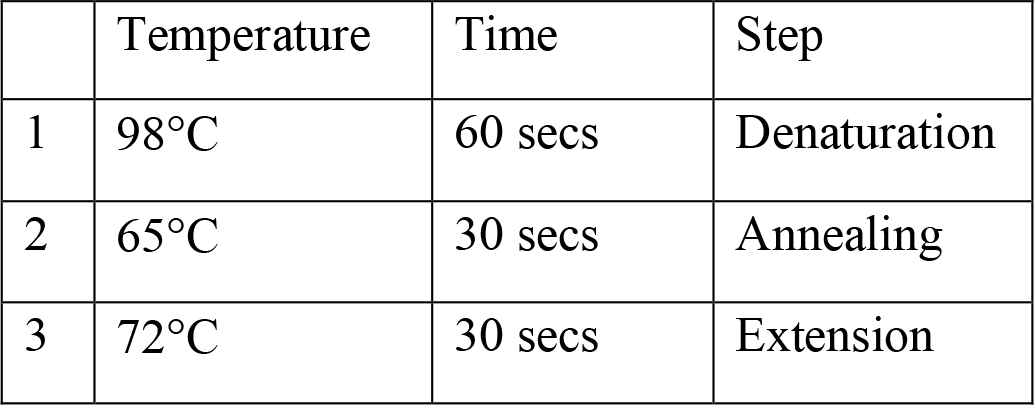
3. Clean-up reactions using 1.4X AMPure XP bead clean-up and elute in 50 μl of DEPC water per replicate. These samples are now ready for jPCR step.

### (2K)#Junction PCR (jPCR)

➢ Perform same number jPCR reactions as RT reactions. Use 10 μl of cDNA library per jPCR reaction.

➢ Use modified jPCR primers as provided by Neumayr and colleagues in their protocol (Neumayr et al. 2019).

➢ jPCR allows amplification of only self-transcribed fragments and filters out plasmid transcripts

#### jPCR protocol

1. Set-up following reactions for each replicate. (5 reactions per replicate)

2. Incubate reactions in thermocycler (program: jPCR)

|  | Sample/reagent | Vol (µl) |
| --- | --- | --- |
| 1 | cDNA library | 10 |
| 2 | DEPC water | 10 |
| 3 | jPCR forward primer | 2.5 |
| 4 | jPCR reverse primer | 2.5 |
| 5 | KAPA 2X HiFi Ready Mix | 25 |
|  | Total | 50 |

3. Clean-up jPCR reactions by pooling all 5 reactions per replicate (250 μl per replicate) and perform a 0.8X AMPure XP clean-up using 200 μl of beads per replicate.
4. Elute in 50 μl DEPC water. Measure concentration and purity of final jPCR products prior to performing final sequencing ready PCR step.
5. Typical yield is around 50 – 130 ng/μl of 50 μl product. Yields may vary based on STARR-seq activity and starting amount for samples.

- 1132 2. Incubate reactions in thermocycler (program: jPCR)
|  | Temperature | Time | Step |
| --- | --- | --- | --- |
| 1 | +98°C | 45 secs | Initial denaturation |
| 2 | +98°C | 15 secs | Denaturation |
| 3 | +65°C | 30 secs | Annealing |
| 4 | +72°C | 30 secs | Extension |
| 5 | +72°C | 60 secs | Final extension |
| 6 | +4°C | Hold |  |

### (2L)#Sequencing ready PCR: Output library preparation

➢ This step is carried out to add i5 and P5 index adapters to the library before sending samples for sequencing.

➢ Use standard i5 primer as forward primer and P7 primer as shown in Neumayr et al 2019 and supplementary table S2.

➢ Dilute all primers to 10 μM prior to use.

➢ Bring AMPure XP beads to room temperature for at least 30 mins for PCR clean-up

➢ Prepare 1% agarose gel for test PCR run to determine number of cycles for PCR. Start with low cycle number. 5 cycles are sufficient for jPCR products obtained in the previous step.

➢ If performing test PCR, test on all samples since they may have different starting amounts.

#### Sequencing Ready PCR protocol

1. Setup the following reaction for each replicate:

2. Incubate reactions on thermocycler (program: SeqReady PCR)

|  | Sample/reagent | Vol (μl) |
| --- | --- | --- |
| 1 | jPCR product | 10 |
| 2 | DEPC water | 10 |
| 3 | i5 primer | 2.5 |
| 4 | P7 primer | 2.5 |
| 5 | KAPA 2X HiFi Ready Mix | 25 |
|  | Total | 50 |

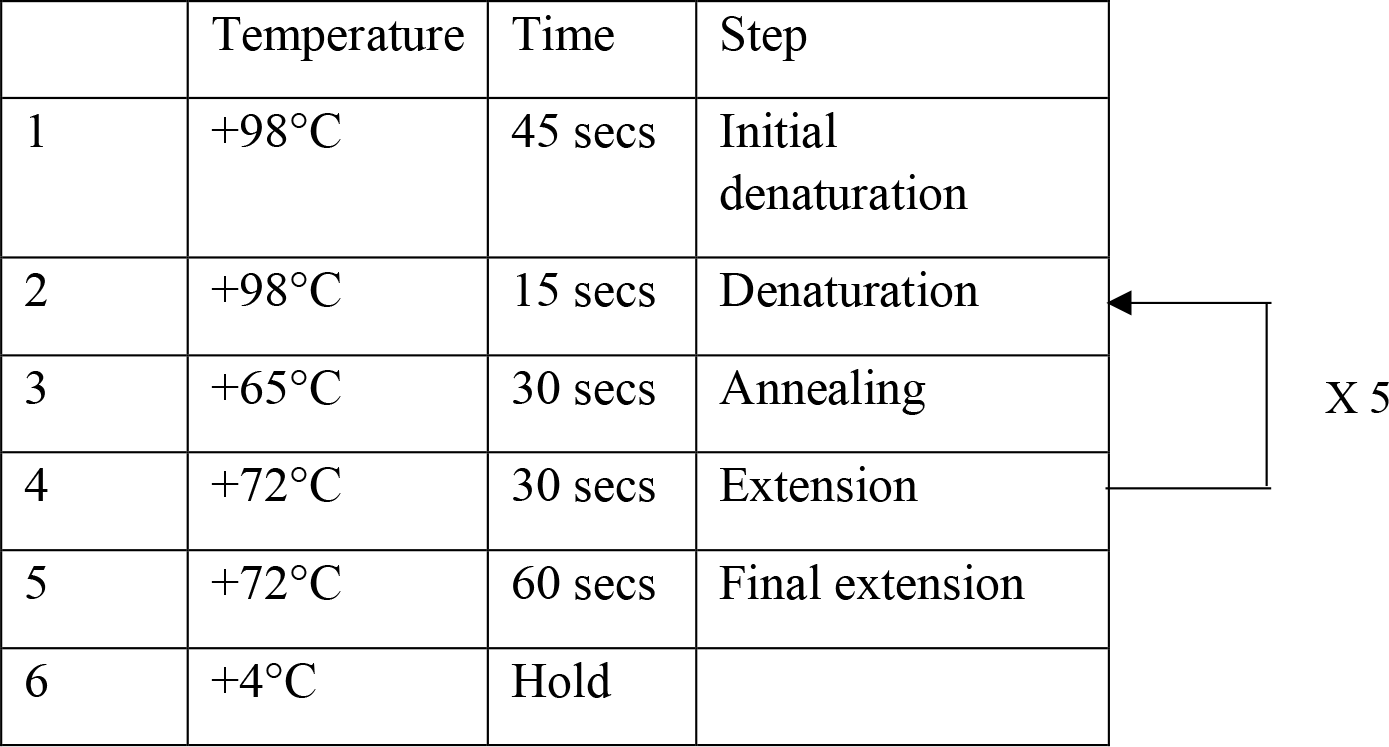
3. Run 10 μl sample with 2 μl of 6X loading dye (for each replicate) on 1% agarose gel for 30 mins at 130V and observe band. Check for characteristic smear for each band at right size as shown by Neumayr and colleagues (Neumayr et al. 2019). If band is concentrated, repeat PCR with lower cycles (if cycle >5) or try lower starting material (5 μl of jPCR).
4. Repeat PCR for correct starting amount and number of cycles as determined.
5. Clean up each reaction with 1X AMPure XP beads and elute in 20 μl. Typical yield can vary and may need to be diluted prior to sequencing.
6. Samples once sent to core facility will be reassessed for quality, checked for distribution on tapestation, further diluted and pooled into an equimolar pool for sequencing.

### (2M)#Sequencing ready PCR: Input library preparation

➢ To assess enhancer activity, plasmid library needs to be sequenced alongside output libraries for normalization.

➢ Library sequence architecture should be identical to enable pooling of library along with output libraries.

➢ Input library also goes through PCR steps and so also important to add UMIs to these libraries prior to sequencing similar to output libraries.

➢ Use custom i7-UMI-P7 primers designed for UMI addition. Use i5 primer to add i5 index and P5 adapter. Assign unique dual index pairs to each replicate of input library and use at least 3 replicates. Record indexes assigned. Dilute primers to 10 μM prior to use.

➢ Bring AMPureXP clean-up beads to room temperature for at least 30 mins for clean-up.

➢ Prepare 1% agarose gel for final library fragment length verification and purification.

➢ For starting material, use 1 µl of STARR-seq plasmid library (∼750 ng/ µl) added with 39 µl of DEPC water.

#### UMI_PCR protocol for input library preparation

1. Setup following reaction for each input replicate:

2. Incubate reaction in thermocycler (program: UMI_PCR)
3. Perform 1.4X AMPure XP clean-up of each replicate reaction. Elute in 20 μl DEPC water.

| | Sample/reagent | Vol (μl) |
| --- | --- | --- |
| 1 | STARR-seq plasmid library | 40 |
| 2 | KAPA HiFi 2X Ready mix | 50 |
| 3 | i7-UMI-P7 primer | 5 |
| 4 | i5 primer | 5 |
| 5 | Total | 100 |

#### Sequencing Ready PCR protocol for input library preparation

1. Setup following reaction for each input replicate: (use complete product)

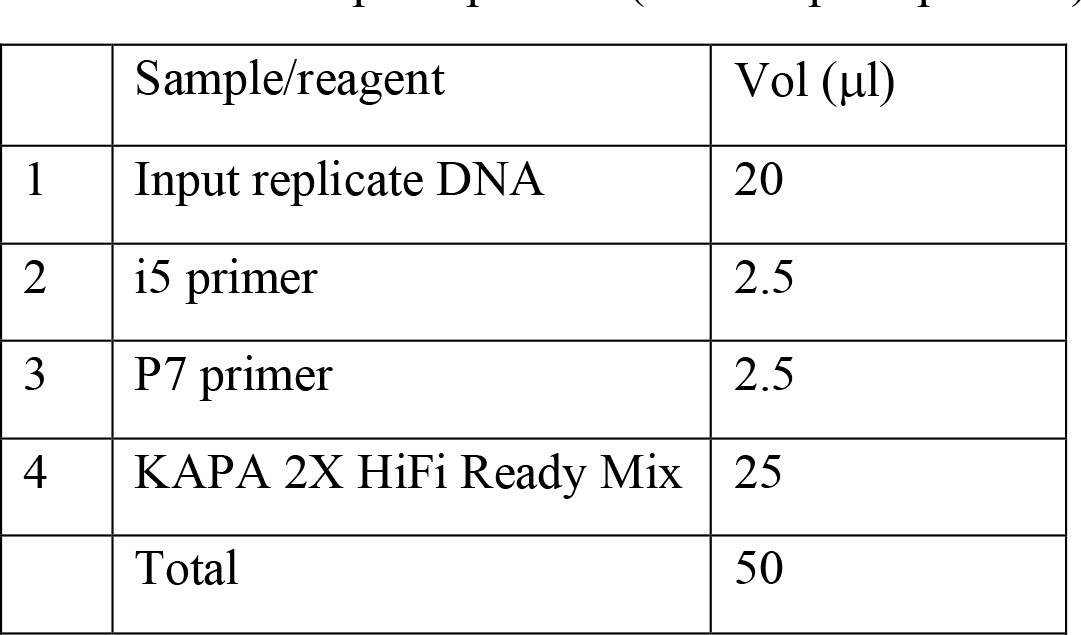
2. Incubate reactions in thermocycler (program: SeqReady PCR)
3. Run entire sample (50 μl) with 10 μl 6X loading dye on a 1% agarose gel for 30 mins at 130V.
4. Visualize gel and verify fragment length. Slice out band of correct length and weigh gel slice.
5. Extract DNA form gel slice using Zymoclean Gel DNA recovery kit.
6. Clean-up extracted DNA using 1X AMPureXP bead clean-up and elute in 25 µl of DEPC water and measure concentration and purity. Typical yield is ∼ 10 – 20 ng/µl.

|  | Sample/reagent | Vol (μl) |
| --- | --- | --- |
| 1 | Input replicate DNA | 20 |
| 2 | i5 primer | 2.5 |
| 3 | P7 primer | 2.5 |
| 4 | KAPA 2X HiFi Ready Mix | 25 |
|  | Total | 50 |

➢ All input and output libraries are sent to sequencing core facility for sequencing.

## Notes

### Competing Interest Statement

The authors have declared no competing interest.

### Summary of Updates

Summary of revisions (1) Rationales for selecting STARR-seq designs: We have added Supplemental_Table_S1 to provide rationales and recommendations on selecting different STARR-seq design factors based on the goal of the assay. This table provides recommendations for selecting library size, fragment length, and DNA source, as well as shows examples of published STARR-seq studies that investigated similar questions. This table will help distinguish between different STARR-seq designs and applications, understand their strengths and limitations, and guide researchers design more informed STARR-seq experiments. (2) Assay scaling guidelines: We have added Supplemental_Table_S2 to provide guidance on assay scaling. This table uses library size and fragment length as a starting point and allows readers to calculate the required minimum library complexity, the number of transformation reactions, and the minimum number sequencing reads necessary to achieve sufficient read depth. (3) Design considerations for choice of vector and promoter: We have expanded the section on experimental design factors to include key considerations for selecting the appropriate STARR-seq vector and a compatible basal promoter. Based on extensive literature review, we describe adaptations and applications of different vectors and promoters for biological contexts. (4) Selection of biological controls: We have added a new sub-section on selecting biological controls for STARR-seq that includes selection criteria, associated challenges, and alternatives. As an example, we have added Supplemental_Fig_S6 differentiating STARR-seq enhancer activity from activity within an exonic region, treated as a negative control region. We have also emphasized the use of control reactions during library preparation in order to assess library complexity. (5) Reanalysis of published datasets: We have reanalyzed four datasets from three studies representing whole-genome and focused STARR-seq experiments. We compared various read filtering and quality control metrics, such as read loss during filtering, correlations between library replicates and assessment of batch effects between the published studies and results from our data. We provide detailed descriptions of these analyses in Supplemental Methods and in the accompanying figures (Supplemental Fig_S2-S5). We also compare STARR-seq activity of similar regions across published datasets and identify inconsistencies in results, illustrating challenges with reproducibility across studies (Supplemental_Fig_S7). (6) Descriptions for newly generated data use: We now provide descriptions for our STARR-seq experimental design and analysis in Supplemental Text and Supplemental Methods as well as data quality assessments (Supplemental_Fig_S3) to provide readers with a clearer understanding of our data.

## References

1. Arnold. Cosmas D, Gerlach. Daniel, Stelzer. Christoph, Boryń. Łukasz M, Rath. Martina SA. 2013. Genome-Wide Quantitative Enhancer Activity Maps Identified by STARR-seq. Science (80- ) 339.

2. Arnold CD, Gerlach D, Stelzer C, Boryń ŁM, Rath M, Stark A. 2013. Genome-wide quantitative enhancer activity maps identified by STARR-seq. Science (80- ) 339: 1074–1077.

3. Bailey TL, Elkan C. 1994. Fitting a mixture model by expectation maximization to discover motifs in biopolymers. Proc Int Conf Intell Syst Mol Biol 2: 28–36.

4. Banerji J, Rusconi S, Schaffner W. 1981. Expression of a β-globin gene is enhanced by remote SV40 DNA sequences. Cell 27: 299–308.

5. Barakat TS, Halbritter F, Zhang M, Rendeiro AF, Perenthaler E, Bock C, Chambers I. 2018. Functional Dissection of the Enhancer Repertoire in Human Embryonic Stem Cells. Cell Stem Cell 23: 276–288.e8.

6. Bergman DT, Jones TR, Liu V, Ray J, Jagoda E, Siraj L, Kang HY, Nasser J, Kane M, Rios A, et al. 2022. Compatibility rules of human enhancer and promoter sequences. Springer US.

7. Birney E, Stamatoyannopoulos JA, Dutta A, Guigó R, Gingeras TR, Margulies EH, Weng Z, Snyder M, Dermitzakis ET, Thurman RE, et al. 2007. Identification and analysis of functional elements in 1% of the human genome by the ENCODE pilot project. Nature 447: 799–816.

8. Boyle AP, Davis S, Shulha HP, Meltzer P, Margulies EH, Weng Z, Furey TS, Crawford GE. 2008. High-resolution mapping and characterization of open chromatin across the genome. Cell 132: 311–22. http://www.ncbi.nlm.nih.gov/pubmed/18243105 (Accessed August 9, 2019).

9. Brandt MM, Meddens CA, Louzao-Martinez L, Van Den Dungen NAM, Lansu NR, Nieuwenhuis EES, Duncker DJ, Verhaar MC, Joles JA, Mokry M, et al. 2018. Chromatin Conformation Links Distal Target Genes to CKD Loci. J Am Soc Nephrol 29: 462–476.

10. Broad Institute. 2019. “Picard Toolkit.” GitHub Repository. https://broadinstitute.github.io/picard/; Broad Institute

11. Buenrostro JD, Giresi PG, Zaba LC, Chang HY, Greenleaf WJ. 2013. Transposition of native chromatin for fast and sensitive epigenomic profiling of open chromatin, DNA-binding proteins and nucleosome position. Nat Methods 10: 1213–1218.

12. Chaudhri VK, Dienger-Stambaugh K, Wu Z, Shrestha M, Singh H. 2020. Charting the cis-regulome of activated B cells by coupling structural and functional genomics. Nat Immunol 21: 210–220. http://dx.doi.org/10.1038/s41590-019-0565-0.

13. Conesa A, Madrigal P, Tarazona S, Gomez-Cabrero D, Cervera A, McPherson A, Szcześniak MW, Gaffney DJ, Elo LL, Zhang X, et al. 2016. A survey of best practices for RNA-seq data analysis. Genome Biol 17: 1–19.

14. Daley T, Smith AD. 2013. Predicting the molecular complexity of sequencing libraries. Nat Methods 10: 325–327.

15. de Almeida BP, Reiter F, Pagani M, Stark A. 2022. DeepSTARR predicts enhancer activity from DNA sequence and enables the de novo design of synthetic enhancers. Nat Genet 54: 613– 624.

16. Errington TM, Denis A, Perfito N, Iorns E, Nosek BA. 2021. Challenges for assessing replicability in preclinical cancer biology. Elife 10: 1–32.

17. Freedman LP, Cockburn IM, Simcoe TS. 2015. The economics of reproducibility in preclinical research. PLoS Biol 13: 1–9.

18. Glaser L V., Steiger M, Fuchs A, Van Bömmel A, Einfeldt E, Chung HR, Vingron M, Meijsing SH. 2021. Assessing genome-wide dynamic changes in enhancer activity during early mESC differentiation by FAIRE-STARR-seq. Nucleic Acids Res 49: 12178–12195.

19. Gross DS, Garrard WT. 1988. Nuclease hypersensitive sites in chromatin. Annu Rev Biochem 57: 159–197.

20. Haberle V, Arnold CD, Pagani M, Rath M, Schernhuber K, Stark A. 2019. Transcriptional cofactors display specificity for distinct types of core promoters. Nature 570: 122–126. http://dx.doi.org/10.1038/s41586-019-1210-7.

21. Haberle V, Stark A. 2018. Eukaryotic core promoters and the functional basis of transcription initiation. Nat Rev Mol Cell Biol 19: 621–637. http://dx.doi.org/10.1038/s41580-018-0028-8.

22. Halper SM, Hossain A, Salis HM. 2020. Synthesis Success Calculator: Predicting the Rapid Synthesis of DNA Fragments with Machine Learning. ACS Synth Biol 9: 1563–1571.

23. Hansen TJ, Hodges E. 2022. ATAC-STARR-seq reveals transcription factor–bound activators and silencers within chromatin-accessible regions of the human genome. Genome Res 32: 1529–1541.

24. Heintzman ND, Stuart RK, Hon G, Fu Y, Ching CW, Hawkins RD, Barrera LO, Van Calcar S, Qu C, Ching KA, et al. 2007. Distinct and predictive chromatin signatures of transcriptional promoters and enhancers in the human genome. Nat Genet 39: 311–318.

25. Heinz S, Benner C, Spann N, Bertolino E, Lin YC, Laslo P, Cheng JX, Murre C, Singh H, Glass CK. 2010. Simple Combinations of Lineage-Determining Transcription Factors Prime cis- Regulatory Elements Required for Macrophage and B Cell Identities. Mol Cell 38: 576– 589. http://dx.doi.org/10.1016/j.molcel.2010.05.004.

26. Hong CKY, Cohen BA. 2022. Genomic environments scale the activities of diverse core promoters. Genome Res 32: 85–96.

27. Hossain A, Lopez E, Halper SM, Cetnar DP, Reis AC, Strickland D, Klavins E, Salis HM. 2020. Automated design of thousands of nonrepetitive parts for engineering stable genetic systems. Nat Biotechnol 38: 1466–1475. http://dx.doi.org/10.1038/s41587-020-0584-2.

28. Inoue F, Kircher M, Martin B, Cooper GM, Witten DM, McManus MT, Ahituv N, Shendure J. 2017. A systematic comparison reveals substantial differences in chromosomal versus episomal encoding of enhancer activity. Genome Res 27: 38–52.

29. Johnson GD, Barrera A, McDowell IC, D’Ippolito AM, Majoros WH, Vockley CM, Wang X, Allen AS, Reddy TE. 2018. Human genome-wide measurement of drug-responsive regulatory activity. Nat Commun 9: 1–9. http://dx.doi.org/10.1038/s41467-018-07607-x.

30. Jores T, Tonnies J, Wrightsman T, Buckler ES, Cuperus JT, Fields S, Queitsch C. 2021. Synthetic promoter designs enabled by a comprehensive analysis of plant core promoters. Nat Plants 7: 842–855. http://dx.doi.org/10.1038/s41477-021-00932-y.

31. Kalita CA, Brown CD, Freiman A, Isherwood J, Wen X, Pique-Regi R, Luca F. 2018. High-throughput characterization of genetic effects on DNA-protein binding and gene transcription. Genome Res 28: 1701–1708.

32. Kim YS, Johnson GD, Seo J, Barrera A, Cowart TN, Majoros WH, Ochoa A, Allen AS, Reddy TE. 2021. Correcting signal biases and detecting regulatory elements in STARR-seq data. Genome Res 31: 877–889.

33. Kircher M, Sawyer S, Meyer M. 2012. Double indexing overcomes inaccuracies in multiplex sequencing on the Illumina platform. Nucleic Acids Res 40: 1–8.

34. Klein JC, Agarwal V, Inoue F, Keith A, Martin B, Kircher M, Ahituv N, Shendure J. 2020. A systematic evaluation of the design and context dependencies of massively parallel reporter assays. Nat Methods 17: 1083–1091. http://dx.doi.org/10.1038/s41592-020-0965-y.

35. Klein JC, Keith A, Agarwal V, Durham T, Shendure J. 2018. Functional characterization of enhancer evolution in the primate lineage. Genome Biol 19: 1–13.

36. LaFleur TL, Hossain A, Salis HM. 2022. Automated model-predictive design of synthetic promoters to control transcriptional profiles in bacteria. Nat Commun 13.

37. Lajoie BR, Dekker J, Kaplan N. 2015. The Hitchhiker’s guide to Hi-C analysis: Practical guidelines. Methods 72: 65–75.

38. Lambert JT, Su-Feher L, Cichewicz K, Warren TL, Zdilar I, Wang Y, Lim KJ, Haigh J, Morse SJ, Canales CP, et al. 2021. Parallel functional testing identifies enhancers active in early postnatal mouse brain. Elife 10: 1–27.

39. Landt SG, Marinov GK, Kundaje A, Kheradpour P, Pauli F, Batzoglou S, Bernstein BE, Bickel P, Brown JB, Cayting P, et al. 2012. ChIP-seq guidelines and practices of the ENCODE and modENCODE consortia. Genome Res 22: 1813–1831.

40. Lee D, Shi M, Moran J, Wall M, Zhang J, Liu J, Fitzgerald D, Kyono Y, Ma L, White KP, et al. 2020. STARRPeaker: uniform processing and accurate identification of STARR-seq active regions. Genome Biol 21: 1–24.

41. Liu S, Liu Y, Zhang Q, Wu J, Liang J, Yu S, Wei GH, White KP, Wang X. 2017a. Systematic identification of regulatory variants associated with cancer risk. Genome Biol 18: 1–14.

42. Liu Y, Yu S, Dhiman VK, Brunetti T, Eckart H, White KP. 2017b. Functional assessment of human enhancer activities using whole-genome STARR-sequencing. Genome Biol 18: 1– 13.

43. Love MI, Huber W, Anders S. 2014. Moderated estimation of fold change and dispersion for RNA-seq data with DESeq2. Genome Biol 15: 1–21.

44. MacConaill LE, Burns RT, Nag A, Coleman HA, Slevin MK, Giorda K, Light M, Lai K, Jarosz M, McNeill MS, et al. 2018. Unique, dual-indexed sequencing adapters with UMIs effectively eliminate index cross-talk and significantly improve sensitivity of massively parallel sequencing. BMC Genomics 19: 1–10.

45. Martinez-Ara M, Comoglio F, van Arensbergen J, van Steensel B. 2022. Systematic analysis of intrinsic enhancer-promoter compatibility in the mouse genome. Mol Cell 82: 2519–2531.e6. https://doi.org/10.1016/j.molcel.2022.04.009.

46. Melnikov A, Murugan A, Zhang X, Tesileanu T, Wang L, Rogov P, Feizi S, Gnirke A, Callan CG, Kinney JB, et al. 2012. Systematic dissection and optimization of inducible enhancers in human cells using a massively parallel reporter assay. Nat Biotechnol 30: 271–277.

47. Montana CL, Myers CA, Corbo JC. 2011. Quantifying the activity of cis-regulatory elements in the mouse retina by explant electroporation. J Vis Exp 1–7.

48. Muerdter F, Boryn ŁM, Woodfin AR, Neumayr C, Rath M, Zabidi MA, Pagani M, Haberle V, Kazmar T, Catarino RR, et al. 2018. Resolving systematic errors in widely used enhancer activity assays in human cells. Nat Methods 15: 141–149.

49. Mulvey B, Lagunas T, Dougherty JD. 2021. Massively Parallel Reporter Assays: Defining Functional Psychiatric Genetic Variants Across Biological Contexts. Biol Psychiatry 89: 76–89. https://doi.org/10.1016/j.biopsych.2020.06.011.

50. Neumayr C, Pagani M, Stark A, Arnold CD. 2019. STARR-seq and UMI-STARR-seq: Assessing Enhancer Activities for Genome-Wide-, High-, and Low-Complexity Candidate Libraries. Curr Protoc Mol Biol 128: e105.

51. Orabi B, Erhan E, McConeghy B, Volik S V., Le Bihan S, Bell R, Collins CC, Chauve C, Hach F. 2019. Alignment-free clustering of UMI tagged DNA molecules. Bioinformatics 35: 1829–1836.

52. Patwardhan RP, Hiatt JB, Witten DM, Kim MJ, Smith RP, May D, Lee C, Andrie JM, Lee SI, Cooper GM, et al. 2012. Massively parallel functional dissection of mammalian enhancers in vivo. Nat Biotechnol 30: 265–270.

53. Peng T, Zhai Y, Atlasi Y, Ter Huurne M, Marks H, Stunnenberg HG, Megchelenbrink W. 2020. STARR-seq identifies active, chromatin-masked, and dormant enhancers in pluripotent mouse embryonic stem cells. Genome Biol 21: 1–27.

54. Pennacchio LA, Ahituv N, Moses AM, Prabhakar S, Nobrega MA, Shoukry M, Minovitsky S, Dubchak I, Holt A, Lewis KD, et al. 2006. In vivo enhancer analysis of human conserved non-coding sequences. Nature 444: 499–502.

55. Robertson G, Hirst M, Bainbridge M, Bilenky M, Zhao Y, Zeng T, Euskirchen G, Bernier B, Varhol R, Delaney A, et al. 2007. Genome-wide profiles of STAT1 DNA association using chromatin immunoprecipitation and massively parallel sequencing. Nat Methods 4: 651– 657.

56. Sahu B, Hartonen T, Pihlajamaa P, Wei B, Dave K, Zhu F, Kaasinen E, Lidschreiber K, Lidschreiber M, Daub CO, et al. 2022. Sequence determinants of human gene regulatory elements. Nat Genet 54: 283–294.

57. Schöne S, Bothe M, Einfeldt E, Borschiwer M, Benner P, Vingron M, Thomas-Chollier M, Meijsing SH. 2018. Synthetic STARR-seq reveals how DNA shape and sequence modulate transcriptional output and noise. PLoS Genet 14: 1–24.

58. Selvarajan I, Toropainen A, Garske KM, López Rodríguez M, Ko A, Miao Z, Kaminska D, Õunap K, Örd T, Ravindran A, et al. 2021. Integrative analysis of liver-specific non-coding regulatory SNPs associated with the risk of coronary artery disease. Am J Hum Genet 108: 411–430.

59. Shen SQ, Myers CA, Hughes AEO, Byrne LC, Flannery JG, Corbo JC. 2016. Massively parallel cis-regulatory analysis in the mammalian central nervous system. Genome Res 26: 238–255.

60. Shlyueva D, Stelzer C, Gerlach D, Yáñez-Cuna JO, Rath M, Boryń ŁM, Arnold CD, Stark A. 2014. Hormone-Responsive Enhancer-Activity Maps Reveal Predictive Motifs, Indirect Repression, and Targeting of Closed Chromatin. Mol Cell 54: 180–192.

61. Sims D, Sudbery I, Ilott NE, Heger A, Ponting CP. 2014. Sequencing depth and coverage: Key considerations in genomic analyses. Nat Rev Genet 15: 121–132.

62. Smith T, Heger A, Sudbery I. 2017. UMI-tools: Modeling sequencing errors in Unique Molecular Identifiers to improve quantification accuracy. Genome Res 27: 491–499.

63. Su Z, Łabaj PP, Li S, Thierry-Mieg J, Thierry-Mieg D, Shi W, Wang C, Schroth GP, Setterquist RA, Thompson JF, et al. 2014. A comprehensive assessment of RNA-seq accuracy, reproducibility and information content by the Sequencing Quality Control Consortium. Nat Biotechnol 32: 903–914.

64. Tarazona S, García-Alcalde F, Dopazo J, Ferrer A, Conesa A. 2011. Differential expression in RNA-seq: A matter of depth. Genome Res 21: 2213–2223.

65. Tewhey R, Kotliar D, Park DS, Liu B, Winnicki S, Reilly SK, Andersen KG, Mikkelsen TS, Lander ES, Schaffner SF, et al. 2016. Direct identification of hundreds of expression- modulating variants using a multiplexed reporter assay. Cell 165: 1519–1529. http://dx.doi.org/10.1016/j.cell.2016.04.027.

66. Van Ouwerkerk AF, Bosada FM, Liu J, Zhang J, Van Duijvenboden K, Chaffin M, Tucker NR, Pijnappels D, Ellinor PT, Barnett P, et al. 2020. Identification of Functional Variant Enhancers Associated with Atrial Fibrillation. Circ Res 127: 229–243.

67. van Weerd JH, Mohan RA, Duijvenboden K van, Hooijkaas IB, Wakker V, Boukens BJ, Barnett P, Christoffels VM. 2020. Trait-associated noncoding variant regions affect tbx3 regulation and cardiac conduction. Elife 9: 1–26.

68. Vanhille L, Griffon A, Maqbool MA, Zacarias-Cabeza J, Dao LTM, Fernandez N, Ballester B, Andrau JC, Spicuglia S. 2015. High-throughput and quantitative assessment of enhancer activity in mammals by CapStarr-seq. Nat Commun 6.

69. Verfaillie A, Svetlichnyy D, Imrichova H, Davie K, Fiers M, Atak ZK, Hulselmans G, Christiaens V, Aerts S. 2016. Multiplex enhancer-reporter assays uncover unsophisticated TP53 enhancer logic. Genome Res 26: 882–895.

70. Visel A, Blow MJ, Li Z, Zhang T, Akiyama JA, Holt A, Plajzer-Frick I, Shoukry M, Wright C, Chen F, et al. 2009. ChIP-seq accurately predicts tissue-specific activity of enhancers. Nature 457: 854–858.

71. Vockley CM, Guo C, Majoros WH, Nodzenski M, Scholtens DM, Hayes MG, Lowe WL, Reddy TE. 2015. Massively parallel quantification of the regulatory effects of noncoding genetic variation in a human cohort. Genome Res 25: 1206–1214.

72. Wang X, He L, Goggin SM, Saadat A, Wang L, Sinnott-Armstrong N, Claussnitzer M, Kellis M. 2018. High-resolution genome-wide functional dissection of transcriptional regulatory regions and nucleotides in human. Nat Commun 9. http://dx.doi.org/10.1038/s41467-018-07746-1.

73. White MA, Myers CA, Corbo JC, Cohen BA. 2013. Massively parallel in vivo enhancer assay reveals that highly local features determine the cis-regulatory function of ChIP-seq peaks. Proc Natl Acad Sci U S A 110: 11952–11957.

74. Yan F, Powell DR, Curtis DJ, Wong NC. 2020. ATAC-seq data analysis. Genome Biol 21: 1–16.

75. Zabidi MA, Arnold CD, Schernhuber K, Pagani M, Rath M, Frank O, Stark A. 2015. Enhancer- core-promoter specificity separates developmental and housekeeping gene regulation. Nature 518: 556–559. http://dx.doi.org/10.1038/nature13994.

76. Zhang P, Xia JH, Zhu J, Gao P, Tian YJ, Du M, Guo YC, Suleman S, Zhang Q, Kohli M, et al. 2018. High-throughput screening of prostate cancer risk loci by single nucleotide polymorphisms sequencing. Nat Commun 9. http://dx.doi.org/10.1038/s41467-018-04451-x.

## Reference

77. Arnold. Cosmas D, Gerlach. Daniel, Stelzer. Christoph, Boryń. Łukasz M, Rath. Martina SA. 2013. Genome-Wide Quantitative Enhancer Activity Maps Identified by STARR-seq. Science (80- ) 339.

78. Barakat TS, Halbritter F, Zhang M, Rendeiro AF, Perenthaler E, Bock C, Chambers I. 2018. Functional Dissection of the Enhancer Repertoire in Human Embryonic Stem Cells. Cell Stem Cell 23: 276–288.e8.

79. Bergman DT, Jones TR, Liu V, Ray J, Jagoda E, Siraj L, Kang HY, Nasser J, Kane M, Rios A, et al. 2022. Compatibility rules of human enhancer and promoter sequences. Springer US.

80. Birney E, Stamatoyannopoulos JA, Dutta A, Guigó R, Gingeras TR, Margulies EH, Weng Z, Snyder M, Dermitzakis ET, Thurman RE, et al. 2007. Identification and analysis of functional elements in 1% of the human genome by the ENCODE pilot project. Nature 447: 799–816.

81. Broad Institute. 2019. “Picard Toolkit.” GitHub Repository. https://broadinstitute.github.io/picard/; Broad Institute

82. Chaudhri VK, Dienger-Stambaugh K, Wu Z, Shrestha M, Singh H. 2020. Charting the cis- regulome of activated B cells by coupling structural and functional genomics. Nat Immunol 21: 210–220. http://dx.doi.org/10.1038/s41590-019-0565-0.

83. Glaser L V., Steiger M, Fuchs A, Van Bömmel A, Einfeldt E, Chung HR, Vingron M, Meijsing SH. 2021. Assessing genome-wide dynamic changes in enhancer activity during early mESC differentiation by FAIRE-STARR-seq. Nucleic Acids Res 49: 12178–12195.

84. Hansen TJ, Hodges E. 2022. ATAC-STARR-seq reveals transcription factor–bound activators and silencers within chromatin-accessible regions of the human genome. Genome Res 32: 1529–1541.

85. Inoue F, Kircher M, Martin B, Cooper GM, Witten DM, McManus MT, Ahituv N, Shendure J. 2017. A systematic comparison reveals substantial differences in chromosomal versus episomal encoding of enhancer activity. Genome Res 27: 38–52.

86. Johnson GD, Barrera A, McDowell IC, D’Ippolito AM, Majoros WH, Vockley CM, Wang X, Allen AS, Reddy TE. 2018. Human genome-wide measurement of drug-responsive regulatory activity. Nat Commun 9: 1–9. http://dx.doi.org/10.1038/s41467-018-07607-x.

87. Kalita CA, Brown CD, Freiman A, Isherwood J, Wen X, Pique-Regi R, Luca F. 2018. High- throughput characterization of genetic effects on DNA-protein binding and gene transcription. Genome Res 28: 1701–1708.

88. Kim YS, Johnson GD, Seo J, Barrera A, Cowart TN, Majoros WH, Ochoa A, Allen AS, Reddy TE. 2021. Correcting signal biases and detecting regulatory elements in STARR-seq data. Genome Res 31: 877–889.

89. Klein JC, Agarwal V, Inoue F, Keith A, Martin B, Kircher M, Ahituv N, Shendure J. 2020. A systematic evaluation of the design and context dependencies of massively parallel reporter assays. Nat Methods 17: 1083–1091. http://dx.doi.org/10.1038/s41592-020-0965-y.

90. Lambert JT, Su-Feher L, Cichewicz K, Warren TL, Zdilar I, Wang Y, Lim KJ, Haigh J, Morse SJ, Canales CP, et al. 2021. Parallel functional testing identifies enhancers active in early postnatal mouse brain. Elife 10: 1–27.

91. Lee D, Shi M, Moran J, Wall M, Zhang J, Liu J, Fitzgerald D, Kyono Y, Ma L, White KP, et al. 2020. STARRPeaker: uniform processing and accurate identification of STARR-seq active regions. Genome Biol 21: 1–24.

92. Li H. 2013. Aligning sequence reads, clone sequences and assembly contigs with BWA-MEM. 00: 1–3. http://arxiv.org/abs/1303.3997.

93. Li H, Handsaker B, Wysoker A, Fennell T, Ruan J, Homer N, Marth G, Abecasis G, Durbin R. 2009. The Sequence Alignment/Map format and SAMtools. Bioinformatics 25: 2078–2079.

94. Liu S, Liu Y, Zhang Q, Wu J, Liang J, Yu S, Wei GH, White KP, Wang X. 2017a. Systematic identification of regulatory variants associated with cancer risk. Genome Biol 18: 1–14.

95. Liu Y, Yu S, Dhiman VK, Brunetti T, Eckart H, White KP. 2017b. Functional assessment of human enhancer activities using whole-genome STARR-sequencing. Genome Biol 18: 1– 13.

96. Love MI, Huber W, Anders S. 2014. Moderated estimation of fold change and dispersion for RNA-seq data with DESeq2. Genome Biol 15: 1–21.

97. Melnikov A, Murugan A, Zhang X, Tesileanu T, Wang L, Rogov P, Feizi S, Gnirke A, Callan CG, Kinney JB, et al. 2012. Systematic dissection and optimization of inducible enhancers in human cells using a massively parallel reporter assay. Nat Biotechnol 30: 271–277.

98. Muerdter F, Boryn ŁM, Woodfin AR, Neumayr C, Rath M, Zabidi MA, Pagani M, Haberle V, Kazmar T, Catarino RR, et al. 2018. Resolving systematic errors in widely used enhancer activity assays in human cells. Nat Methods 15: 141–149.

99. Neumayr C, Pagani M, Stark A, Arnold CD. 2019. STARR-seq and UMI-STARR-seq: Assessing Enhancer Activities for Genome-Wide-, High-, and Low-Complexity Candidate Libraries. Curr Protoc Mol Biol 128: e105.

100. Patwardhan RP, Hiatt JB, Witten DM, Kim MJ, Smith RP, May D, Lee C, Andrie JM, Lee SI, Cooper GM, et al. 2012. Massively parallel functional dissection of mammalian enhancers in vivo. Nat Biotechnol 30: 265–270.

101. Peng T, Zhai Y, Atlasi Y, Ter Huurne M, Marks H, Stunnenberg HG, Megchelenbrink W. 2020. STARR-seq identifies active, chromatin-masked, and dormant enhancers in pluripotent mouse embryonic stem cells. Genome Biol 21: 1–27.

102. Quinlan AR, Hall IM. 2010. BEDTools: A flexible suite of utilities for comparing genomic features. Bioinformatics.

103. Ran FA, Hsu PD, Wright J, Agarwala V, Scott DA, Zhang F. 2013. Genome engineering using the CRISPR-Cas9 system. Nat Protoc 8: 2281–2308.

104. Sahu B, Hartonen T, Pihlajamaa P, Wei B, Dave K, Zhu F, Kaasinen E, Lidschreiber K, Lidschreiber M, Daub CO, et al. 2022. Sequence determinants of human gene regulatory elements. Nat Genet 54: 283–294.

105. Schöne S, Bothe M, Einfeldt E, Borschiwer M, Benner P, Vingron M, Thomas-Chollier M, Meijsing SH. 2018. Synthetic STARR-seq reveals how DNA shape and sequence modulate transcriptional output and noise. PLoS Genet 14: 1–24.

106. Shlyueva D, Stelzer C, Gerlach D, Yáñez-Cuna JO, Rath M, Boryń ŁM, Arnold CD, Stark A. 2014. Hormone-Responsive Enhancer-Activity Maps Reveal Predictive Motifs, Indirect Repression, and Targeting of Closed Chromatin. Mol Cell 54: 180–192.

107. Vanhille L, Griffon A, Maqbool MA, Zacarias-Cabeza J, Dao LTM, Fernandez N, Ballester B, Andrau JC, Spicuglia S. 2015. High-throughput and quantitative assessment of enhancer activity in mammals by CapStarr-seq. Nat Commun 6.

108. Verfaillie A, Svetlichnyy D, Imrichova H, Davie K, Fiers M, Atak ZK, Hulselmans G, Christiaens V, Aerts S. 2016. Multiplex enhancer-reporter assays uncover unsophisticated TP53 enhancer logic. Genome Res 26: 882–895.

109. Wang X, He L, Goggin SM, Saadat A, Wang L, Sinnott-Armstrong N, Claussnitzer M, Kellis M. 2018. High-resolution genome-wide functional dissection of transcriptional regulatory regions and nucleotides in human. Nat Commun 9. http://dx.doi.org/10.1038/s41467-018-07746-1.

110. Zabidi MA, Arnold CD, Schernhuber K, Pagani M, Rath M, Frank O, Stark A. 2015. Enhancer- core-promoter specificity separates developmental and housekeeping gene regulation. Nature 518: 556–559. http://dx.doi.org/10.1038/nature13994.

111. Zhang Y, Liu T, Meyer CA, Eeckhoute J, Johnson DS, Bernstein BE, Nussbaum C, Myers RM, Brown M, Li W, et al. 2008. Model-based analysis of ChIP-Seq (MACS). Genome Biol 9.

